# Early Treatment with Oral Pirfenidone Improves Bladder Function after Contusive Spinal Cord Injury in Mice

**DOI:** 10.64898/2026.08.19.745817

**Authors:** Alonso Carlos Agustin Isidro, Murugapoopathy Vasikar, Curran Laura, Rivard Leane, Bharti Anamitra, Kassouf Wassim, Janzen Jan, David Samuel, R Gupta Indra

## Abstract

Spinal cord injury (SCI) disrupts innervation to the lower urinary tract, resulting in bladder dysfunction that predisposes to urinary infections and renal impairment. While inflammation is central to bladder pathology after SCI, the molecular events linking acute to chronic remodeling are poorly defined. We hypothesized that early treatment with pirfenidone, an anti-inflammatory and anti-fibrotic drug, would attenuate bladder pathology after SCI. Adult female C57BL/6J mice underwent contusive SCI or sham laminectomy, and bladders were collected at 2, 7, 16, and 45 days later. SCI induced bladder hypertrophy, edema, hemorrhage, neutrophil infiltration, cell proliferation and loss of voiding function in the first 48 hours. Transcriptomic profiling at this timepoint was characterized by activation of inflammatory and cytokine pathways including TNF-α, IL-6, the complement cascade, and TGFβ. Although bladder function partially recovered by day 7, inflammatory pathways persisted and extracellular matrix (ECM) remodeling programs emerged. By day 16, robust activation of ECM-remodeling pathways was evident in all bladders. Treatment with pirfenidone during the acute inflammatory phase (day 2–7) reduced bladder hypertrophy and suppressed expression of pro-fibrotic, inflammatory, and neuroplasticity-associated genes including *Bdnf* and *Chrm2* that encodes muscarinic receptor 2 (M2). Mechanistically, pirfenidone attenuated TGFβ signaling as shown by downregulation of phosphoSmad2 protein in whole bladders and decreased M2 receptor expression in the urothelium. These molecular changes correlated with improved function in pirfenidone-treated mice as shown by fewer voiding events with larger urine volumes up until 45 days after SCI. Early treatment with pirfenidone limits inflammation and fibrosis, normalizes neural signaling, and improves bladder function after SCI.

**New & Noteworthy:** Using a mouse model of traumatic spinal cord injury, we showed that bladder inflammation and dysfunction precede fibrosis, with TGFβ serving as a key regulatory hub. Transient pirfenidone treatment during the acute inflammatory phase (days 2-7) attenuated bladder pathology, limited *Tgfb2/3*, *Bdnf*, and *Chrm2* expression, and reduced M2 receptor and phospho-Smad2 protein levels. These molecular events correlated with improved bladder storage function, highlighting an early therapeutic window and TGFβ signaling as a target for neurogenic bladder remodeling.

## Introduction

Spinal cord injury (SCI) disrupts the innervation to multiple organ systems, including the urinary tract. Quality of life after SCI is severely affected by bladder dysfunction. There is a lack of synchronization between contractions of the detrusor muscle and opening and closing of the external urethral sphincter that results in impaired storage and emptying of urine. Bladder dysfunction after SCI thus causes high bladder pressures, and an increased risk of urinary tract infections, hydronephrosis, and long-term kidney damage. Chronic urinary retention and inflammation promote a pro-fibrotic environment within the bladder wall, characterized by excessive deposition of extracellular matrix proteins and a decrease in bladder compliance. Current treatment for bladder dysfunction, also called the neurogenic bladder, is focused on the use of anti-cholinergic medication to improve storage function and/or bladder catheterization to facilitate emptying, but both are of limited benefit. There is therefore a compelling need to elucidate the inflammatory pathways in the bladder after SCI to identify novel approaches to prevent or mitigate fibrosis and preserve bladder function.

In this work, we defined bladder dysfunction in a mouse model of spinal cord injury induced by a moderate contusion. The model recapitulates human SCI secondary to trauma and was characterized in-depth using bulk RNA sequencing, RT-qPCR, histology and immunofluorescent studies to identify the molecular pathways involved in bladder dysfunction. We found that early inflammation, within 48 hours after SCI, was driven by neutrophils and TNF-α, while later stages were characterized by sustained TNF-α and TGFβ signaling, and cytokine release. We, therefore hypothesized that early treatment with the anti-inflammatory and anti-fibrotic drug known as pirfenidone would limit bladder inflammation and fibrosis and improve bladder function after SCI.

## Materials and Methods

### Mice

Adult C57BL/6J female mice, aged 8–16 weeks old and a minimum of 20 g in body weight were used in this study. Mice were housed in a 12 h light/dark cycle, temperature-controlled environment with ad libitum access to food and water as per the ARRIVE guidelines (1). Laminectomy (sham) or spinal cord injury (SCI) was performed at the T9/T10 level using combined anesthesia with ketamine, xylazine, and acepromazine. A moderate contusion injury of 50 kdyn was induced using an Infinite Horizon impactor with displacements ranging between 400-600 µm (2). After the surgery, all animals were given analgesia with 0.1 mg/kg buprenorphine twice per day for 48 hours, and hydration as needed. Manual bladder compression was performed twice a day for the first 7 days after SCI. For the pirfenidone experiments, female mice were randomized to control or treatment groups. Pirfenidone (Sigma, P2116) was administered via oral gavage at a dose of 200 mg/kg/day beginning on day 2 after injury and continuing for 7 days. Vehicle (0.5% carboxymethylcellulose, CMC)-treated mice received an equivalent volume of 200 μl. Mice were euthanized at d2, d7, d16, and d45 and bladders were weighed and processed for subsequent analyses. All animal studies were performed in accordance with the regulations of the Canadian Council on Animal Care and approved by the Animal Care Committee of the Research Institute of the McGill University Health Centre (AUP 8204).

### Human Sections

Histological sections of human bladders from SCI patients were kindly provided by Dr. med. Jan Janzen, VascPath Bern, Switzerland. Both samples belonged to male patients (40 and 52 years old) and were obtained 13 and 10 years after spinal injury, respectively. Bladder control biopsies from patients without bladder disease were kindly provided by Dr. Wassim Kassouf, Department of Surgery, McGill University, Montreal, Canada.

### Bulk RNA-Sequencing

Bladders from sham and SCI mice were collected at 2 and 7 days post-injury, as well as from sham, SCI + CMC and SCI + pirfenidone mice at 16 days post-injury. Tissue was snap-frozen in liquid nitrogen and stored at −80°C until needed. Frozen tissue was pulverized using a mortar and pestle and RNA was extracted using the RNeasy Mini Kit (Qiagen 74104) following the manufacturer’s instructions. The purity and concentration of the RNA samples was evaluated using a NanoDrop One spectrophotometer (ThermoFisher Scientific). Libraries were prepared using the NEB poly A mRNA magnetic isolation module and the NEB Ultra II directional RNA kits and sequenced on an Illumina platform NovaSeq 6000 to generate paired-end reads. Raw reads were demultiplexed using Trimmomatic v0.39 (3) and aligned to Mm10 (grcm38/2011) of the mouse genome using Hisat2. SAM files were converted to BAM using Samtools. Aligned reads were counted using Featurecounts, and counts per gene were analyzed using DESeq2 readily available on the Galaxy platform (https://usegalaxy.org). DEGs were defined with significance set at an FDR adjusted p-value < 0.01 and a log2fold change of (<-1; >1), representing a 2-fold change.

### Real-Time Quantitative PCR

RNA was reverse transcribed using Moloney murine leukemia virus reverse transcriptase and random hexamer primers. qPCR reactions were performed using Blastaq Master mix (Applied Biological Materials G891) using a LightCycler® 96 Instrument (Roche). Expression of *Bdnf* (GeneID: 12064)*, Chrm2*(GeneID:1129), *Chrm3* (GeneID:12671)*, Trpv1 (*GeneID: 193034) *Col1a1* (GeneID: 12842)*, Col3a1* (GeneID:12825), *Il6* (GeneID:14573)*, Il1b* (GeneID :16176)*, Tnf* (GeneID: 21926)*, Rpl13* (GeneID:6137), *Serpine1* (GeneID :57344)*, Tgfb1 (*GeneID*:21803) Tgfb2* (GeneID:19713), and *Tgfb3* (GeneID:20472) were analyzed. Gene expression per sample was analyzed in duplicate, averaged and normalized to the reference gene *Rpl13* using the 2-ΔΔCt method. Primer sets are listed in Table 1.

**Table 1.**
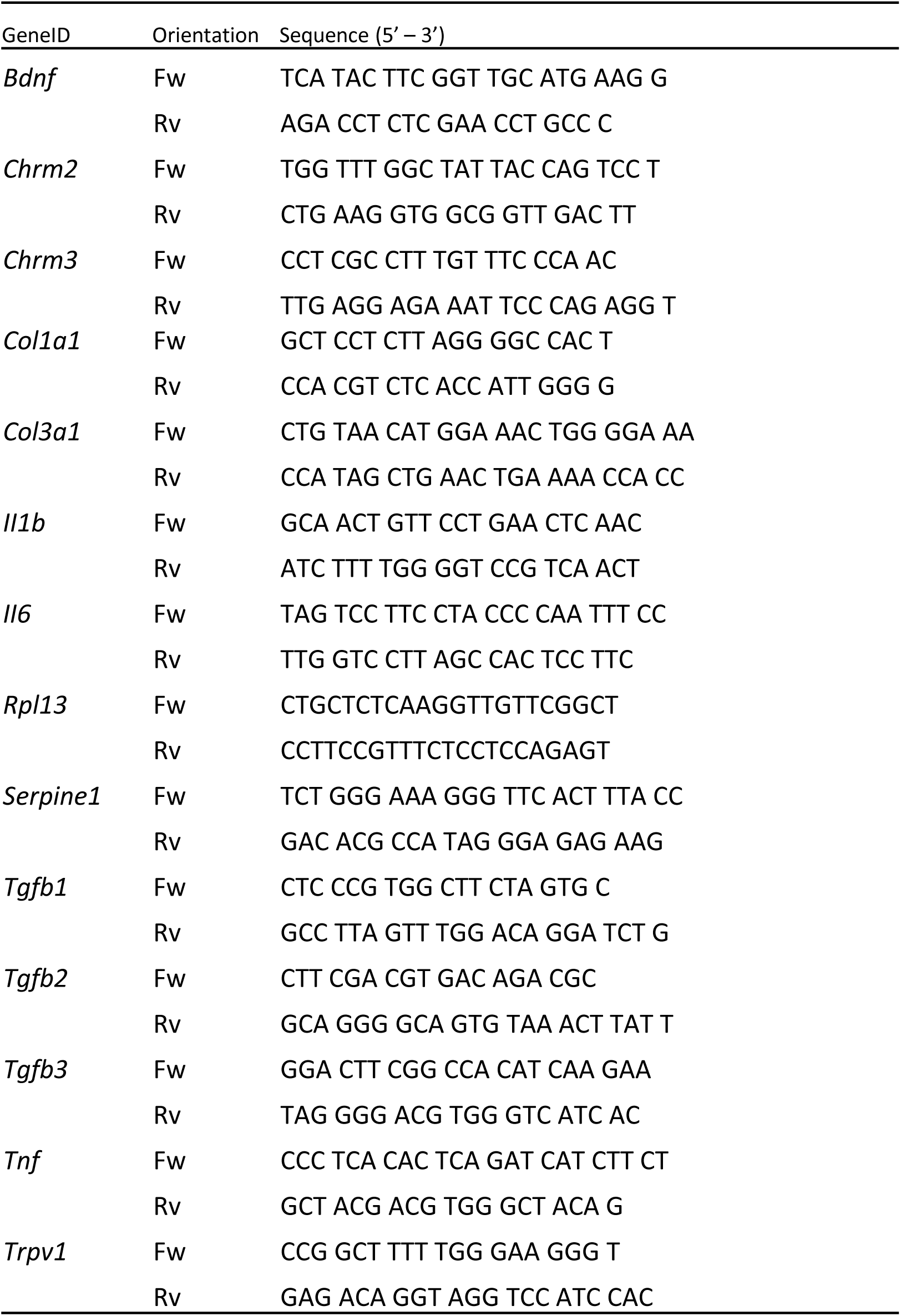
List of primers.

### Histology

Mouse bladders were fixed in 4% PFA overnight, embedded in paraffin, and sectioned at 5 μm thickness. Sections were stained with hematoxylin and eosin or Masson’s trichrome or Verhoeff-van Gieson following standard protocols. Images were acquired using a Evos7000M system (Invitrogen).

### Immunofluorescence

Paraffin-embedded tissue sections were deparaffinized, rehydrated in phosphate-buffered saline (PBS), and subjected to heat-induced antigen retrieval in citrate buffer. After cooling to room temperature, sections were blocked with 1% bovine serum albumin (BSA) in PBS for 1 hour. Primary antibodies were applied at a 1:100 dilution and incubated overnight at 4 °C. The antibodies used, commercial suppliers and catalogue numbers are listed in Table 2. The following day, sections were washed and incubated with secondary antibodies (Invitrogen) for 1 hour at room temperature at a 1:500 dilution. Secondary antibodies included Alexa Fluor 488-conjugated anti-rabbit, Alexa Fluor 555-conjugated anti-rabbit, and Alexa Fluor 555-conjugated anti-mouse. Nuclear counterstaining was performed using DAPI. After final washes, sections were mounted using Fluorsave (Fisher Scientific) and imaged with a Leica LSM780 confocal microscope (Richmond Hill, ON, Canada). Image processing and quantification were performed using Zen Black software (Zeiss) and Fiji (ImageJ).

**Table 2.**
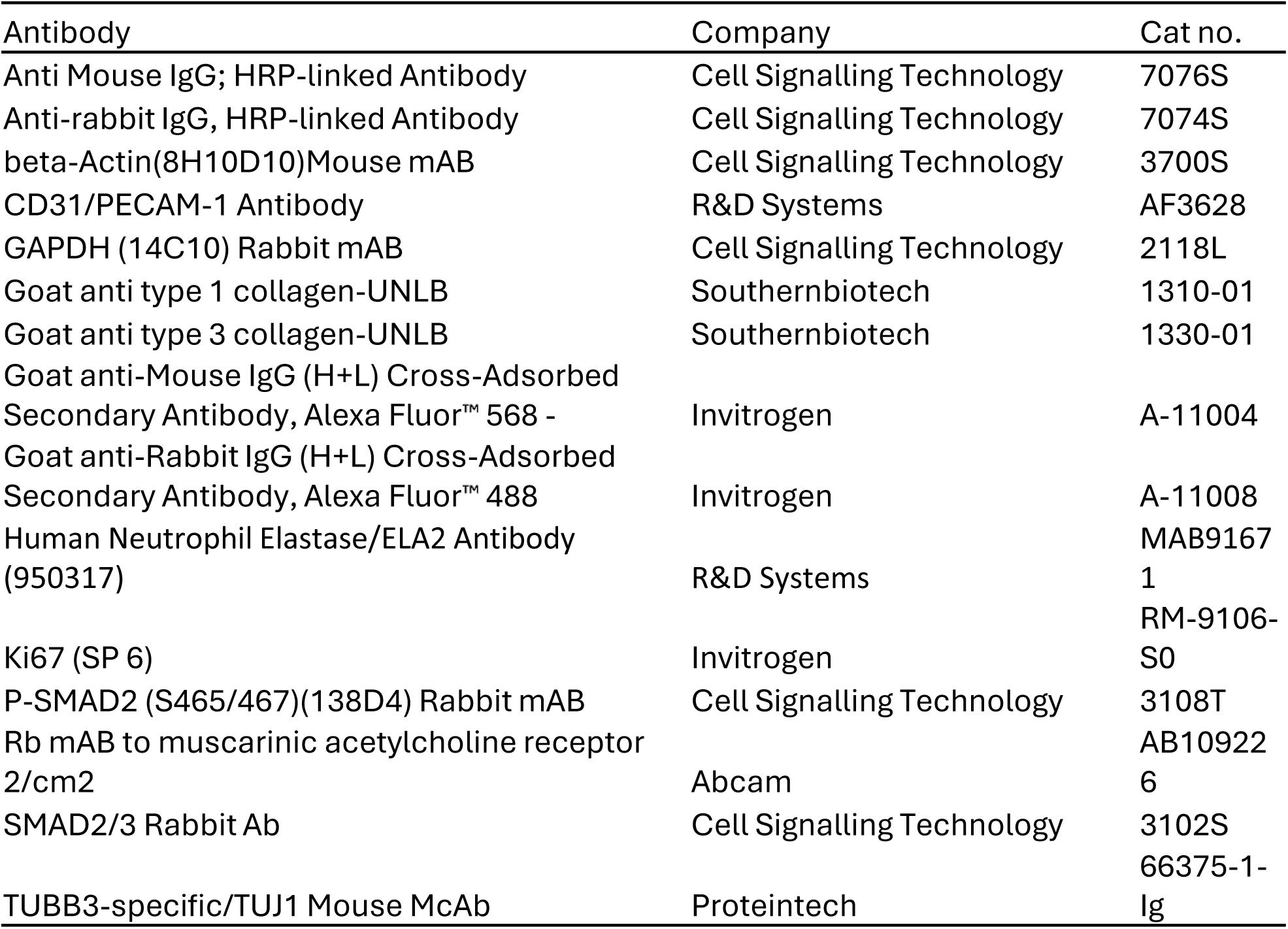
List of antibodies and vendors.

For whole mount bladder staining, animals were perfused with 4% PFA and further fixed in 4% PFA after dissection at 4°C overnight. Bladders were washed with PBS 5x 1 h each and then dehydrated and cleared using the Transluscence tissue clearing kit (Transluscence Biosystems) -based on the iDISCO protocol- and following the manufacturer’s instructions. TUJ1 and CD31 primary antibody were incubated at a concentration of 1:100 for 3 days at 37°C on a nutator. After clearing, the samples were maintained in ethyl cinnamate. Imaging for TUJ1 was performed on a Zeiss LSM780 confocal microscope using the Z stack function (50-60 slices spaced 1 µm apart). Imaging for CD31 was performed on Zeiss Lightsheet Z.1 microscope (Jena, Germany). The whole bladder was imaged with a Z stack distance of 14 µm. Rendered images were obtained from Zen software ZEN2010 (TUJ1) or Imaris 10.2.0 (CD31).

### Western Blot

Bladder halves were snap frozen and triturated with a mortar and pestle. The pulverized tissue was resuspended in 300 μl of RIPA buffer supplemented with protease inhibitors and further sheared using a 20 G syringe, followed by 3 sonication pulses (10 s at 50Hz). Lysates were cleared by centrifugation at 12,000 x g for 20 min at 4°C. Thirty µg of protein was mixed with 5X Laemmli buffer (250 mM Tris, 10% w/v SDS, 0.2 % w/v bromophenol blue, 50 % v/v glycerol) and proteins were resolved using a 10% PAGE gel. Proteins were transferred to nitrocellulose membranes, which were blocked with 3% BSA in TBS-T (0.05% Tween-20) for 1 h at room temperature. Membranes were then incubated with antibodies against pSMAD2 (1:2000), SMAD2 (1:1000), or COL1A1 (1:5000), COL3A1 (1:1000), M2 (1:500), TUJ1 (1:1000), β-actin (1:5000) or GAPDH (1:3000) overnight at 4°C. The following day, membranes were washed for 10 min with TBS-T for 3 washes and incubated with secondary antibodies against mouse or rabbit (1:5000) conjugated to horseradish peroxidase. Bands were visualized using a commercial ECL kit (ZmTech Scientifique, Cat. E208082) as per the manufacturer’s instructions and imaged using an Amersham Imager 600 (GE Healthcare). Between each primary antibody incubation, membranes were stripped with 0.2 M NaOH for 40 min at room temperature and re-blocked with 3% BSA. For epithelial/mesenchymal separation, bladder halves were incubated in 500 µl Dispase in HBSS (STEMCELL technologies, 07913) on an orbital shaker for 30 minutes at room temperature. Epithelia were then pulled from the mesenchyme using tweezers and both tissues were moved to PBS and minced on a cold surface using a scalpel. Minced tissues were resuspended in 500 μl of ice-cold RIPA buffer with protein inhibitors and placed in 2 ml tubes for lysis using a TissueLyser II (Qiagen) at 30 Hz for 1 min. Lysates were subsequently sheared with a 20 G syringe and sonicated and cleared as described above.

### Hydroxyproline Assay

Collagen assessment was performed using the hydroxyproline assay kit by Sigma (MAK0008). Briefly, bladder halves were weighed, snap frozen and pulverized with a mortar and pestle. Samples were resuspended in 100 μL of water and further disrupted with a 20 G blunt needle. 100 μl of the bladder homogenate was mixed with 100 μl of 12 N HCl and incubated for 3 h at 120°C. Fifty μl of each sample was seeded in a 96 well plate and evaporated. A concentration curve (0, 0.2, 0.4, 0. 6, 0.8 and 1 μg) of collagen was used to assess collagen content in the samples. The assay was performed following the manufacturer’s instructions.

### Void Spot Assay

Voiding behavior was assessed using a void spot assay (VSA) as previously described with minor modifications (4). Mice were individually housed in standard cages lined with Whatman grade 1 filter paper (15 x 30 cm). Bladders were manually expressed and mice were administered 250 μl of saline subcutaneously to standardize their hydration status before testing. Water access was removed during the 2h assay period. At the conclusion of the assay, filter papers were collected. Urine spots were revealed after immersing the filter papers in ninhydrin solution (2% w/v), followed by drying overnight and then scanning at 300 ppi. Images were thresholded and particles were analyzed using Fiji. The total number of urine spots, total voided area and average spot size were quantified. Spots smaller than 10 mm^2^ were excluded from analysis.

### Statistical Analysis

Data are presented as mean ± SEM unless otherwise stated. Statistical significance was determined using the unpaired t-test or one-way ANOVA with Tukey’s post hoc test from GraphPad Prism version 8.0.1 for Windows, GraphPad Software (Boston, Massachusetts USA, www.graphpad.com). A p-value < 0.05 was considered statistically significant.

## Results

### Spinal cord injury induces bladder inflammation, early neutrophil infiltration and proliferation

We used a well-established mouse model of spinal cord injury (SCI) that employs controlled contusion to reproduce traumatic spinal cord injury, which is the most common form of injury in humans. While others have used the mouse to model traumatic SCI (5–8), a full transcriptomic and histological analysis of bladder disease has not been reported. From humans and animal models, loss of innervation in the bladder leads to inflammation with bladder hypertrophy, increased collagen deposition and unsynchronized bladder contractions (9). To define the molecular events involved in the acute phase of SCI, we performed transcriptomic analyses on bladders of mice two (d2) and seven days (d7) after a moderate contusion spinal cord injury (SCI) or laminectomy (sham) using bulk RNA sequencing as shown (Fig. 1 and 3, respectively). Differential expression analysis of d2 samples identified 3141 significantly altered genes (log2 fold change < -1; >1, FDR-adjusted p < 0.01) with 1634 upregulated and 1507 downregulated genes (Fig. 1A). Principal component analysis (PCA, Fig. 1B) and sample-to-sample distance metrics (Fig. S1A) revealed two distinct clusters that corresponded to sham and SCI samples. Table 3 lists the top 10 up- and downregulated genes by p-value, with a full list of DEGs available in Supplementary Table 1. During the period of “spinal shock” when manually assisted bladder evacuation is needed (d0-7), KEGG pathway enrichment analysis highlighted terms associated with cell division and inflammation, complement and coagulation cascades, cytokine production and binding, Il-17 signaling, TNF-α signaling, and neuroactive ligand-receptor interaction in bladders from SCI mice (Fig. 1D). KEGG terms reflecting extracellular matrix deposition were not the most highly ranked at day 2 but were included in the heatmaps to track progression from inflammation to fibrosis. A subset of genes belonging to these categories are shown in heatmaps (Fig. 1E). The transcriptomic changes were accompanied by an increase in weight in bladders from SCI mice compared to those from sham mice (Fig. 1C). Additionally, SCI bladders looked distended and visibly red from vascular congestion. Histological analyses of bladder sections from SCI mice revealed edema, blood extravasation and the presence of polymorphonucleated cells in the suburothelium and lamina propria (Fig. 1F). This was consistent with the bulk RNA sequencing results that showed an upregulation of cytokine and TNF-α signaling transcripts that activate the innate immune response. Immunohistochemistry of the myeloid marker, Lymphocyte antigen 6 complex locus G6D (Ly-6G), confirmed that polymorphonucleated cells found in the bladder wall of SCI samples were neutrophils (Fig 1F middle panel). Interestingly, in 3 out of 5 rodent SCI samples, Ly6G-positive node-like structures were identified that expressed the cell cycle protein Ki67 and exhibited DNA fragmentation, as shown by the TUNEL assay (Fig. S1B). On closer examination, the TUNEL signal revealed filament-like structures surrounding the nodes suggesting NETosis from activated neutrophils. Neutrophils are terminally differentiated but can activate cell cycle signaling proteins as shown by the expression of Ki67, however they do not synthesize DNA and re-enter the cell cycle (10). The enrichment of cell cycle proteins from the bulk RNA sequencing analyses therefore, prompted the assessment of Ki67 expression throughout the bladder. There were significantly increased Ki67-positive nuclei in the uroepithelium of SCI bladders compared to shams at d2 (Fig 1F third column). Ki67-positive cells were also detected in the lamina propria of SCI bladders. Only a few proliferative cells were detected in the uroepithelium of sham bladders, consistent with its low cell turnover at baseline (11). In summary, there is an induction of inflammation associated genes, neutrophil invasion and strong proliferative response in the bladder as early as two days after SCI.

**Figure 1.**
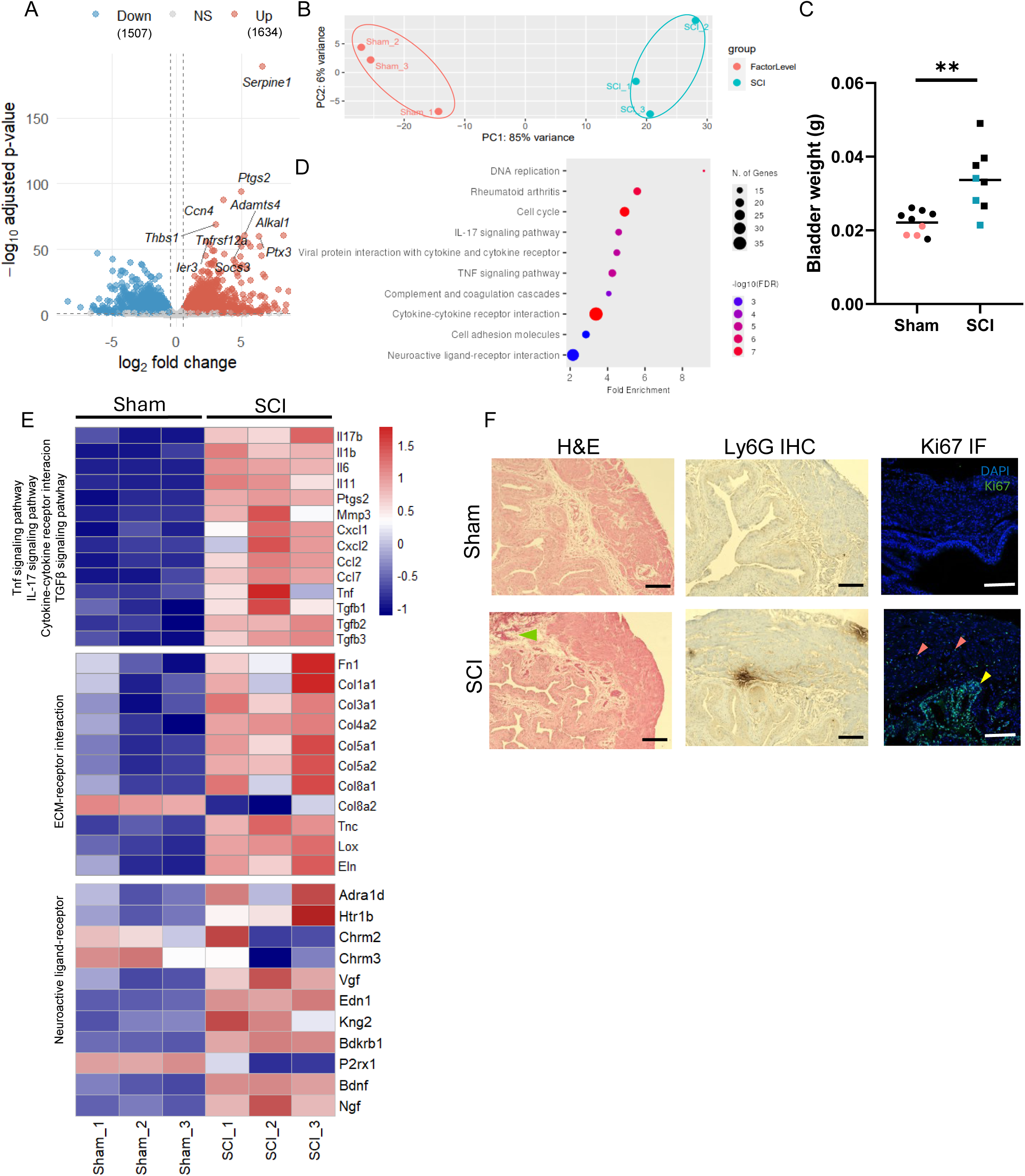
Spinal cord injury induces bladder inflammation, early neutrophil infiltration and proliferation during the first 48 hours. (A) Volcano plot of differentially expressed genes (DEGs) in bladders from SCI vs. sham mice at d2, identified by bulk RNA sequencing (log2FC <-1 or >1, FDR-adjusted p<0.01; n=3 sham, n=3 SCI). (B) Principal component analysis (PCA) of RNAseq samples. (C) Bladder weight at d2 (line indicates mea; t-test, n=9 sham, n=8 SCI). Coloured dots indicate the samples that were subjected to bulk RNA sequencing (pink: sham; blue: SCI). (D) KEGG pathway enrichment analysis of upregulated DEGs. (E) Heatmaps of representative genes from TNF-α signaling/IL17 signaling/cytokine-cytokine receptor interaction (top), extracellular matrix receptor-interaction (middle), and neuroactive ligand-receptor interaction (bottom) pathways, z-score normalized, at d2. (F) Representative bladder sections showing histology using staining with hematoxylin and eosin, protein expression of Ly6G and Ki67 from sham and SCI mice (n=5/group). Arrowheads indicate Ki67+ nuclei in uroepithelium (pink) and lamina propria (yellow). Scale bars: 150 µm. **: *p* < 0.01.

**Table 3.**
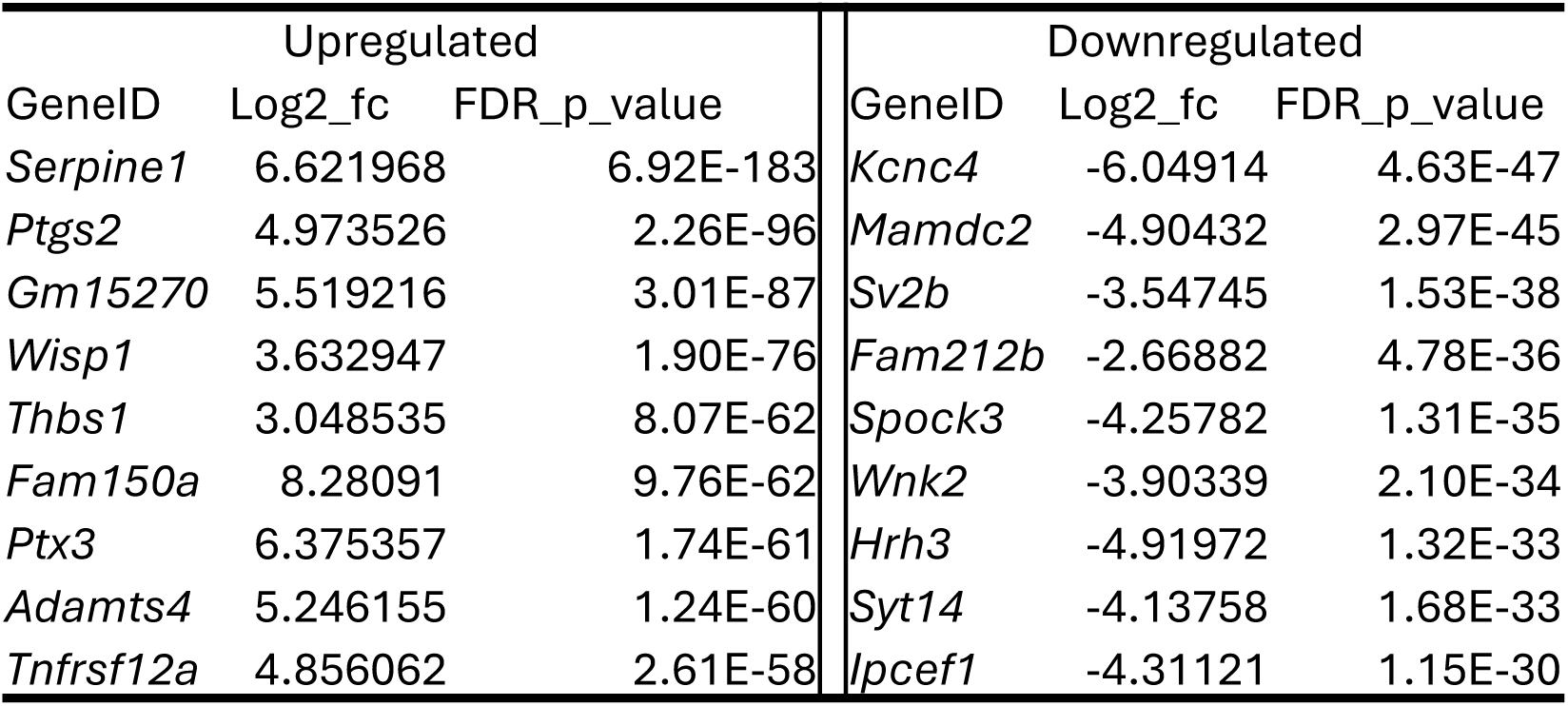
Top ten upregulated and downregulated genes between SCI and sham bladders 2 days after the surgery.

### Histopathology of the neurogenic bladder in patients with SCI

Cystectomy specimens were obtained from 2 adult males undergoing bladder augmentation surgeries. Their spinal cord injuries were at least 10 years prior, and thus, the sections represent findings from a chronic neurogenic bladder from one of the patients (Fig. 2). These were compared with healthy margins from adult male patients with bladder cancer. Severe fibrosis was noted in the suburoepithelial tissue and muscularis in the SCI bladder. Some regions were rich in collagen fibres, as visualised by the Masson-Trichrome stain. A paucicellular hyalinisation of the bladder wall was also observed. Smooth muscle bundles were disorganized and on a background of fibrosis as well as leiomyomatous-like hyperplasia. Neutrophil invasion was also observed in bladder samples from SCI patients as detected using an antibody using an antibody to neutrophil elastin (NE) (Fig. 2 bottom panel) with prominence in the suburothelium.

**Figure 2.**
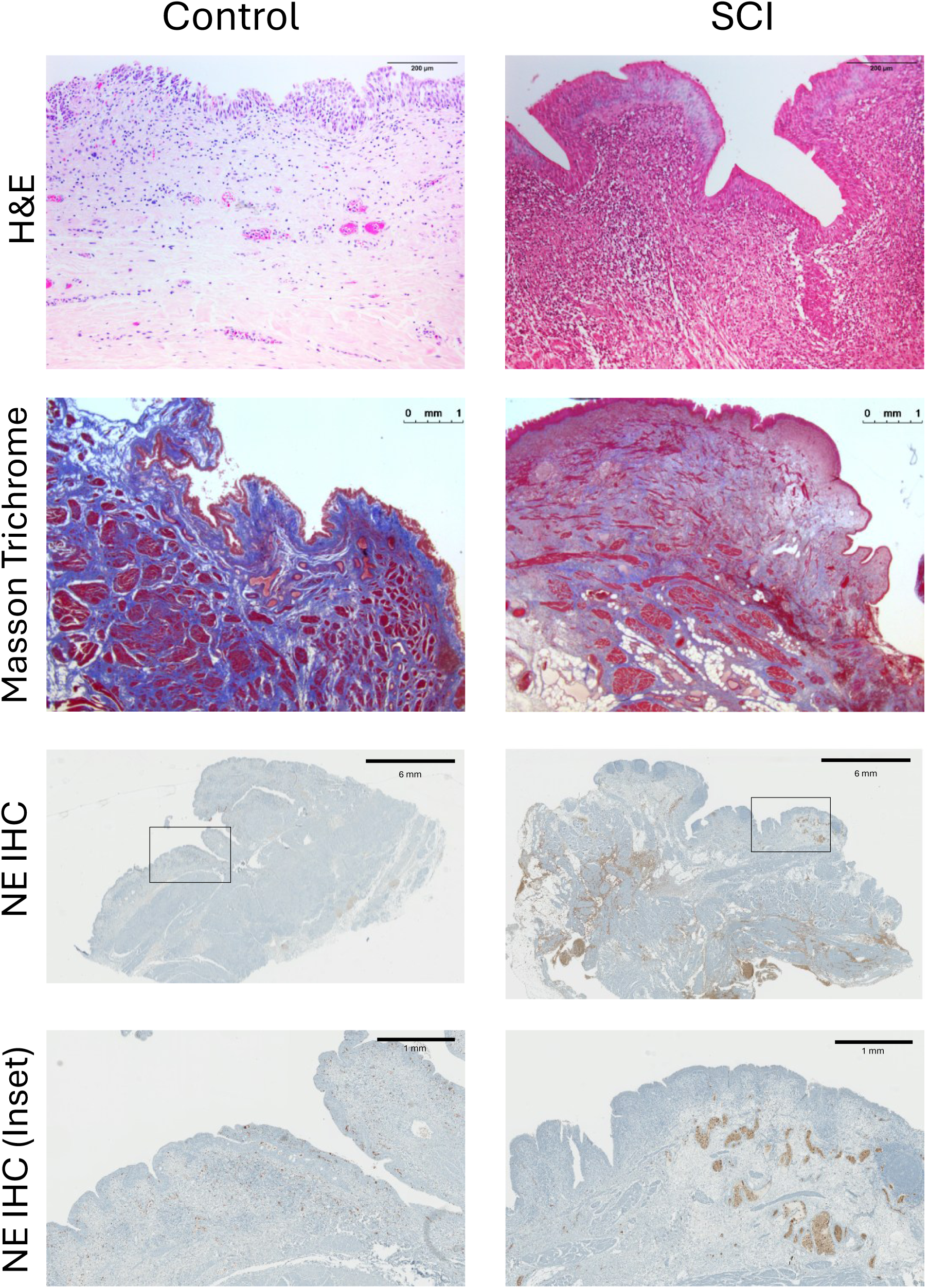
Histological changes in human bladder samples from control and SCI patients. Representative images of bladder sections from biopsies of patients with and without SCI. Top row: Hematoxylin-eosin staining. Second row: Masson’s Trichrome staining showing increased collagen deposition (blue) and marked thickening of the bladder wall following SCI, consistent with extracellular matrix remodeling and fibrosis. Middle row: Low-magnification immunohistochemistry (IHC) for CD14 demonstrating increased infiltration of CD14-positive neutrophils (brown) in the SCI bladder compared with controls. Black boxes indicate the regions shown at higher magnification below. Bottom row: Higher-magnification views of the boxed regions highlighting the increased accumulation of CD14-positive neutrophils within the bladder wall after SCI. Images are representative of n = 2 samples per group. Scale bars: top row, 200 μm; second row, 1 mm; third row, 6 mm; bottom row, 1 mm.

### Bladder inflammation in SCI progresses to an ECM-secretory phenotype

All mice re-gained independent voiding between 7 to 10 days after SCI. Bulk RNA sequencing and differential expression analysis at d7 identified 769 DEGs of which 534 were upregulated and 235 were downregulated (Fig. 3A, Supplementary Table 2). The top 10 most up- and downregulated genes are shown in Table 4. KEGG enrichment analysis revealed overlap with terms from the d2 samples, notably Il-17 and TNF-α signaling, cytokine-cytokine receptor interaction, DNA replication and cell cycle. Other terms like the P53 signaling pathway and cellular senescence were specific to d7 (Fig. 3E). Sample-to-sample distance metrics separated sham and SCI samples (Fig. S1C) however PCA analyses revealed greater variability in the d7 bladders from SCI mice with two distinct groups: one clustered farther and the other closer to the sham samples (Fig. 2B). The transcriptomic profiles revealed that SCI mouse 1 and 2 showed upregulation in genes associated with inflammation, while the bladders from SCI mice (SCI3, 4, and 5) demonstrated an ECM-secretory phenotype, as shown by the upregulation of collagen family and matrix-associated (Fig. 3D) genes. This suggests that some bladders exhibit ongoing injury, while others have initiated a transition to a fibrotic response at d7.

**Figure 3.**
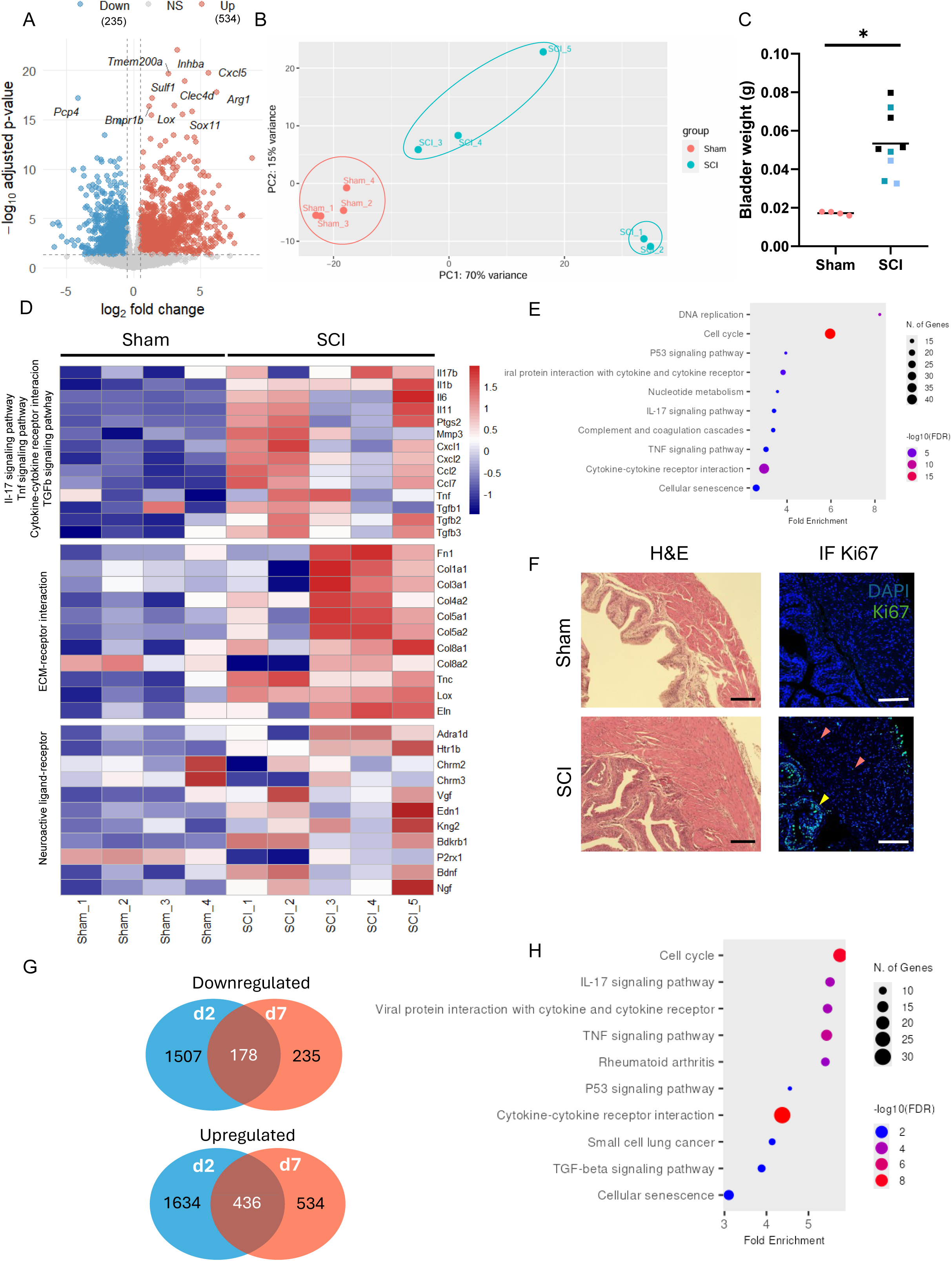
Sustained cell proliferation and emergence of an ECM-secretory transcriptional program at d7 after SCI. (A) Volcano plot of DEGs in SCI vs. sham bladders at d7 (n=3 sham, n=5 SCI). (B) PCA of d7 RNAseq samples. (C) Bladder weight at d7 (line indicates mean; t-test, n=4 sham, n=7 SCI). Coloured dots indicate the samples that were subjected to bulk RNA sequencing (pink: sham; dark blue: SCI 1 and 2; light blue: SCI 3, 4 and 5). (D) Heatmaps of inflammation/cytokine/TGFβ pathway genes (top) extracellular matrix receptor-interaction (middle), and neuroactive ligand-receptor interaction (bottom) pathways, z-score normalized, at d7. (E) KEGG pathway enrichment of DEGs at d7. (F) Representative bladder sections showing staining with hematoxylin and eosin and immunofluorescent detection of Ki67 for sham and SCI bladders at d7 (n=4 sham, n=7 SCI). Scale bars, 150 µm. (G) Venn diagrams of downregulated (left) and upregulated (right) DEGs shared between d2 and d7 in SCI versus sham samples. (H) KEGG pathway enrichment of overlapping DEGs in both d2 and d7 datasets. *: *p* < 0.05.

**Table 4.** Top ten upregulated and downregulated genes between SCI and sham bladders 7 days after the surgery.

| Upregulated |  |  | Downregulated |  |  |
| --- | --- | --- | --- | --- | --- |
| GeneID | Log2_fc | FDR_p_value | GeneID | Log2_fc | FDR_p_value |
| Inhba | 3.282637 | 8.35E-23 | Pcp4 | -4.09649 | 2.00E-18 |
| Cxcl5 | 5.61521 | 1.52E-21 | Rims1 | -2.66958 | 5.94E-12 |
| Tmem200a | 2.637814 | 2.68E-20 | Kcng3 | -2.3807 | 2.90E-11 |
| Clec4d | 3.808857 | 7.80E-20 | Spock3 | -3.71393 | 2.99E-11 |
| Arg1 | 6.196392 | 2.01E-19 | Cpne6 | -4.4746 | 3.92E-11 |
| Lox | 3.072982 | 3.48E-17 | Igsf5 | -1.70987 | 4.76E-11 |
| Sox11 | 4.3975 | 3.48E-17 | Pitpnm3 | -1.41003 | 9.08E-11 |
| AA467197 | 3.653695 | 1.54E-16 | Ankrd63 | -3.69411 | 1.27E-10 |
| Sulf1 | 1.411077 | 4.16E-16 | Deptor | -1.04393 | 2.12E-10 |
| Cemip | 4.546743 | 1.98E-14 | Tmem200c | -2.68979 | 3.13E-10 |

At d7, all bladders from SCI mice were heavier than sham mice (Fig. 3C), however, unlike the bladders at d2, there was now a thickened uroepithelium and markedly expanded detrusor muscle layer without vasculature congestion. Although inflammation-related terms were enriched in the KEGG analysis, there were no Ly-6G positive cells at this timepoint (Fig S1E). Concomitant with the cell cycle and DNA replication terms identified, proliferation was still active at d7 with Ki67-positive cells mostly detected in the uroepithelium, with some in the lamina propria and muscle layer (Fig. 3F). After combining d2 and d7 datasets, an overlap of 436 upregulated genes and 178 downregulated genes were noted (Fig. 3G). Enrichment analysis of upregulated genes in both datasets revealed upregulation of TGFβ signaling effectors that emerged as a novel term (Fig. 3H). Overlapping downregulated genes in d2 and d7 datasets were related to muscle contraction, adrenergic signaling and cyclic nucleotide-associated signaling pathways (Fig. S1D). In summary, proliferation is maintained, and there is a transition occurring amongst the SCI bladders from an inflammatory to an ECM-secretory state at d7.

### Pirfenidone treatment blunts bladder inflammation and hypertrophy

To target inflammation and TGFβ-associated pathways, mice were treated with 5-methyl-1-phenyl-2-[1H]-pyridone, pirfenidone, a compound with known anti-inflammatory and anti-fibrotic properties. Although its precise mechanism of action remains elusive, pirfenidone inhibits TNF-α, TGFβ, and PDGF signaling pathways (12). Mice with SCI received daily oral pirfenidone (200 mg/kg/day) or 0.5% carboxymethyl cellulose (CMC, vehicle) starting at d2 and continuing for 5 days (d2-7, Fig. 4A). This preventive treatment approach is distinct from other therapeutic regimens in which treatment has been initiated after the onset of bladder fibrosis (13–15).

**Figure 4.**
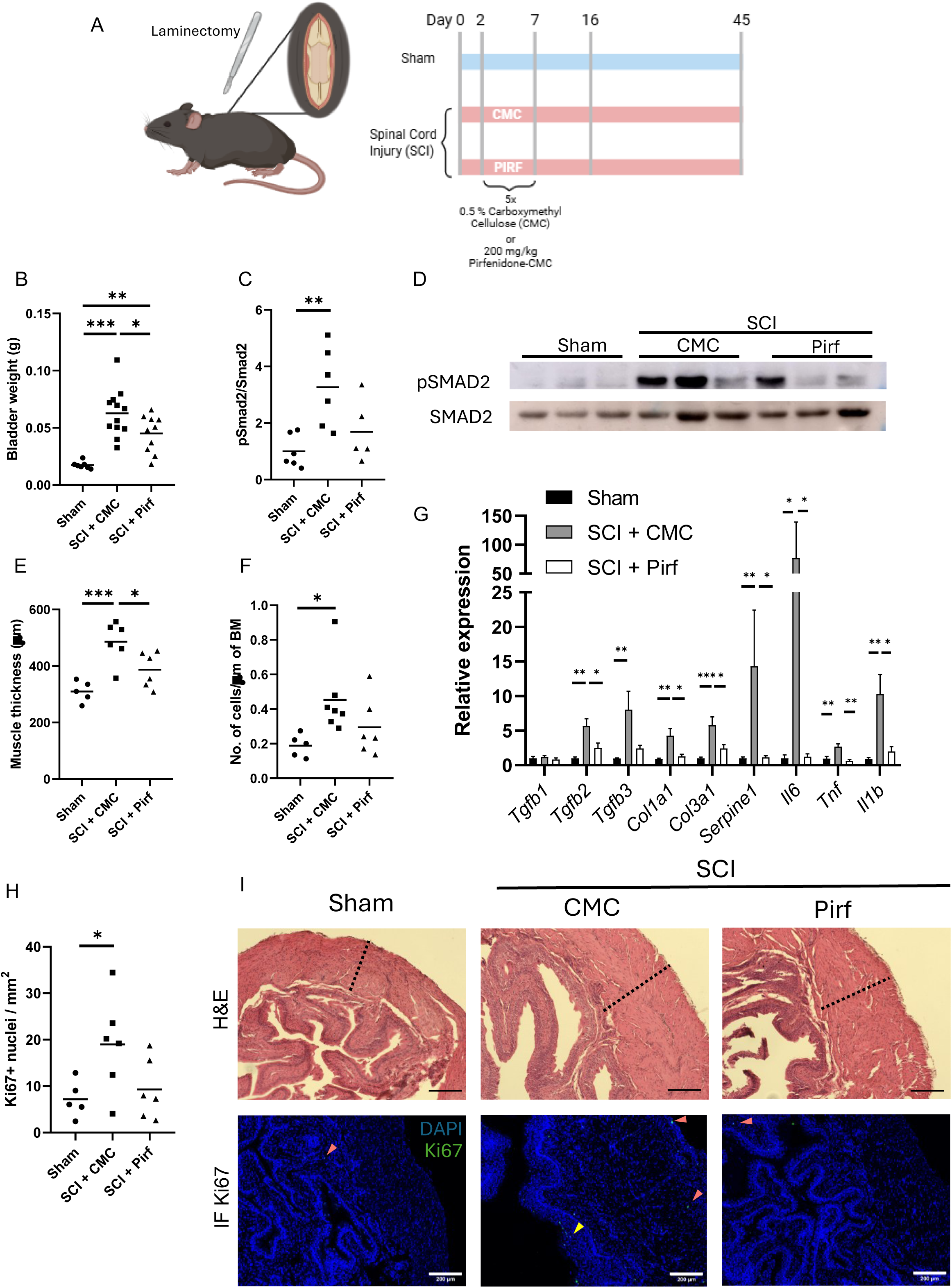
Pirfenidone treatment from day 2 to day 7 reduces bladder weight, the thickness of the detrusor muscle, and inflammatory and TGFβ pathway gene expression. (A) Schematic of treatment paradigm: spinal cords from mice were exposed (laminectomy) and injured with an impactor (SCI) or left intact (sham). Two days later, SCI mice were divided in two groups and administered daily 0.5% CMC (vehicle) or 200 mg/kg/day pirfenidone-CMC (Pirf) orally from day 2 to day 7 (d2–d7); bladders were collected at d7, d16, or d45. Created with BioRender. (B) Bladder weight at d7 in sham, SCI+CMC, and SCI+Pirf groups (line indicates mean; one way ANOVA, n=7 sham, n=12 SCI+CMC, n=10 SCI+Pirf). (C) Quantification of pSMAD2/SMAD2 ratio from densitometry of western blot (n=6/group). (D). Representative western blot of phospho-SMAD2 (pSMAD2) and total SMAD2 from whole bladder lysates. (E) Quantification of detrusor muscle thickness (line indicates mean; ANOVA, sham n=4, n=6 SCI+CMC and n=6 SCI+Pirf). (F) Number of nuclei per µm of basement membrane (line indicates mean; one way ANOVA, n=5 sham, n=7 SCI+CMC, n=6 SCI+Pirf). (G) RT-qPCR of TGFβ pathway components, collagens, and cytokines from whole bladder homogenates [bar indicates mean ± SEM; ANOVA (*Tgfb1*,*Tgfb2, Tgfb3, Col3a1, Tnf-α, Il1b*) or Kruskal-Wallis (*Col1, Serpine1, Il6*) n>6/group]. (H) Quantification of Ki67+ nuclei per mm^2^ of bladder wall (line indicates mean; one way ANOVA, n=5 sham, n=6 SCI+CMC, n=6 SCI+Pirf). (I) Representative images of bladder sections stained with hematoxylin and eosin (upper panel) and Ki67 staining (lower panel). Arrowheads denote Ki67 positive signal. Scale bars, 200 µm. *: *p* < 0.05; **: *p* < 0.01; ***: *p* < 0.001.

The increase in bladder weight in SCI mice at d7 was blunted in pirfenidone-treated mice (Fig. 3B) and accompanied by a decrease in thickness of the detrusor muscle layer (Fig. 3E). There was a trend towards less cellularity (Fig. 4F) and less Ki67-positive cells in the bladders of pirfenidone-treated mice (Fig. 4H), but neither reached statistical significance. To test whether pirfenidone was targeting the TGFβ signaling pathway, TGFβ signaling components and fibrosis-related genes were interrogated using RT-qPCR. *TGFb2/3, Col1a1 and Col3a1* were increased in the bladders from SCI + CMC mice, hereafter referred to as SCI and downregulated in the SCI + pirfenidone mice (Fig. 4G). In contrast, *Tgfb1* was unaltered at d7 in the bladders after SCI, consistent with our transcriptomics results and similar to what has been shown in a genetic mouse model of dilated cardiomyopathy (16). Expression levels of cytokines *Il1b, Il6* and *Tnf-α* were increased in SCI mice and downregulated by pirfenidone treatment. A similar trend was also observed for *Serpine1*, the top upregulated gene at d2 (Table 3) and a target and biomarker of TGFβ activity (16) that was downregulated in the bladders of pirfenidone-treated compared to SCI mice (Fig. 4G). The level of phosphorylated SMAD2 (pSMAD2) protein was also assessed as a readout of TGFβ signaling. Phospho-SMAD2 levels were increased in SCI mice, and this increase was blunted in 2 of 3 bladders from SCI mice (Fig 4C, D). Pirfenidone inhibited inflammation-associated genes and the TGFβ response during the acute phase of SCI, and this correlated with lower bladder weights with less muscle hypertrophy.

### Pirfenidone treatment selectively decreases collagen subtypes following SCI

To determine whether treatment with pirfenidone could limit progression to bladder fibrosis after SCI, analyses were performed at d16. Similar to d7, bladder weight remained elevated in SCI mice compared to sham mice, but it was significantly reduced in pirfenidone-treated mice (Fig. 5A). Although pirfenidone reduced bladder weight at this timepoint, detrusor thickness was comparable between pirfenidone and SCI mice (Fig. 5B, C left panel). To determine whether the extracellular matrix composition changed with bladder weight, four assays were employed including Masson trichrome and Verhoeff Van Gieson staining of bladder sections and hydroxyproline and Western blot analyses of whole bladders. Masson trichrome staining revealed increased collagen deposition in SCI mice and trended towards reduced collagen fiber deposition in pirfenidone-treated mice (Fig. 5C middle panel, E), although this was not statistically significant. Verhoeff-van Gieson staining showed decreased elastin deposition in SCI mice with no apparent differences between pirfenidone-treated mice compared to shams, suggesting that SCI mice had stiffer bladders that were improved with pirfenidone. As expected, total collagen content, measured by hydroxyproline assay, was increased in SCI relative to sham mice, however there was no significant difference between pirfenidone-treated and SCI mice (Fig. 5D). Analysis of specific collagen isoforms revealed a selective effect: at d16 COL1A1 expression was variable in SCI and pirfenidone-treated mice (Fig. 5F, H), whereas COL3A1 protein levels were significantly reduced in pirfenidone-treated mice compared to SCI mice (Fig. 5G, H). The selective reduction in type III collagen, which is associated with early fibrotic remodeling and tissue stiffness, suggests that pirfenidone may modulate the composition rather than the total abundance of collagen. Together, these findings indicate that short-term pirfenidone treatment modifies the ECM composition and reduces overall bladder hypertrophy, consistent with partial attenuation of the fibrotic response.

**Figure 5.**
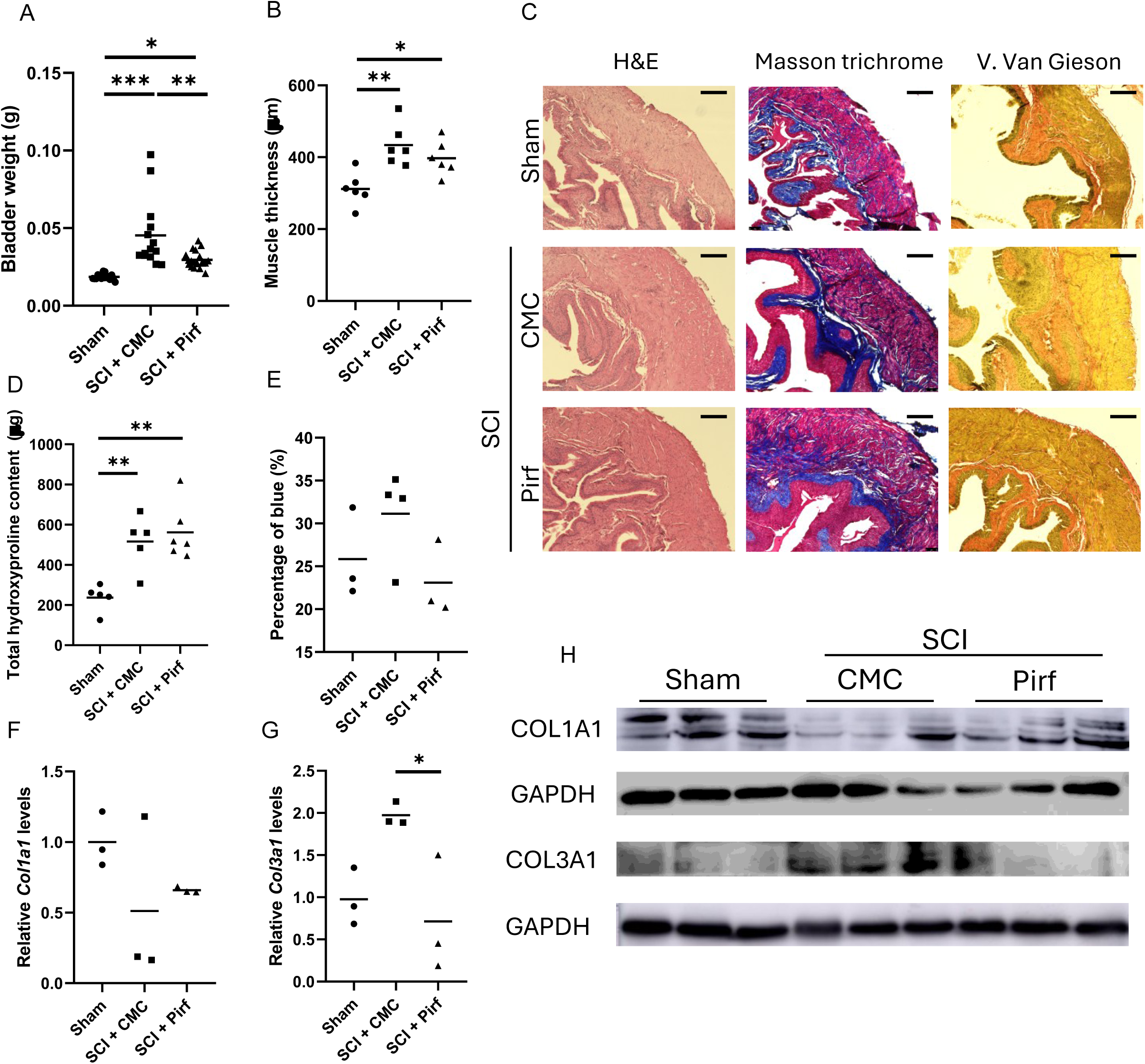
Pirfenidone partially attenuates bladder hypertrophy and selectively reduces type III collagen at d16. (A) Bladder weight at d16 (line indicates mean; ANOVA, sham n=16, n=14 SCI+CMC, n=16 SCI+Pirf). (B) Detrusor muscle thickness at d16 (ANOVA, n=16/group). (C) Representative H&E (left) and Masson trichrome (middle) and Verhoeff-van Gieson (right) staining of bladder cross-sections from sham, SCI+CMC, and SCI+Pirfenidone groups. Scale bars: 150 µm. (D) Total bladder collagen content measured by hydroxyproline assay (line indicates mean; ANOVA, sham n=5, n=6 SCI+CMC, n=6 SCI+Pirf). (E) Quantification of Masson trichrome, blue-stained area (% of total area) (n=4/group, not significant). (F) Quantification and relative COL1A1 and (G) COL3A1 protein expression from densitometry (ANOVA, n=3/group). Densitometry values were normalized to GAPDH and compared to sham bladders. (H) Representative western blots of COL1A1 and COL3A1 protein with corresponding GAPDH loading controls. *: *p* < 0.05; **: *p* < 0.01; ***: *p* < 0.001.

To further define the impact of pirfenidone treatment, we performed transcriptomic analyses of bladders from sham, SCI, and SCI + pirfenidone-treated mice at d16. Comparison between SCI and sham mice revealed 428 DEGs with 374 upregulated and 54 downregulated genes (Fig. 6A, Supplementary Table 3). PCA analysis showed a dispersion of SCI samples along PC1, likely reflecting variability in bladder recovery secondary to the spine contusion (Fig. 6C, green dots). Sham samples aggregated closer together and apart from SCI (Fig. 6C, red dots). The KEGG and GO BP (biological process) enrichment analysis revealed changes in inflammation, ECM remodeling, and cytoskeleton in muscle cells (Fig. 6D). DEG analysis showed upregulation of multiple collagen isoforms (*Col3a1, Col4a2, Col5a1, Col8a1*), fibronectin (*Fn1*), and ECM modifiers such as *Lox* and *Tnc*, suggesting robust fibrotic remodeling of bladder tissue in response to SCI (Fig. 6F). Table 5 shows the top 10 up- and downregulated genes in the bladders of SCI + CMC vs. sham mice. Likelihood-ratio test (LRT) analyses were also performed to assess the interaction between treatment (SCI vs sham) and timepoints post SCI (d2, d7, d16) (Fig. S2). This revealed 335 genes with significant time-dependent changes in expression across conditions (log2 fold change <-1; > 1, p < 0.05 FDR corrected) (Supplementary Table 4). PCA of the resulting expression profiles showed separation of samples primarily driven by time after injury (PC1) (Fig. S2A). Early and late post-injury timepoints showed distinct transcriptional states. Hierarchical clustering of LRT-significant genes with distinct temporal patterns clustered genes in 6 categories (Fig. S2B). Genes associated with inflammatory response or cellular homeostasis and cell cycle were upregulated early and downregulated later. Notably, modules related to extracellular matrix organization and protein maturation exhibited progressive activation at later timepoints, consistent with ongoing tissue remodeling. We inferred immune cell–associated signatures using gene set scoring approaches. SCI samples exhibited increased enrichment of innate immune cell populations, including neutrophils and macrophages, particularly at d2. Adaptive immune signatures, including T cell–related programs, showed more modest and delayed changes (Fig. S2C). These patterns suggest a shift from acute inflammatory infiltration to immune resolution and later-stage tissue remodeling.

**Figure 6.**
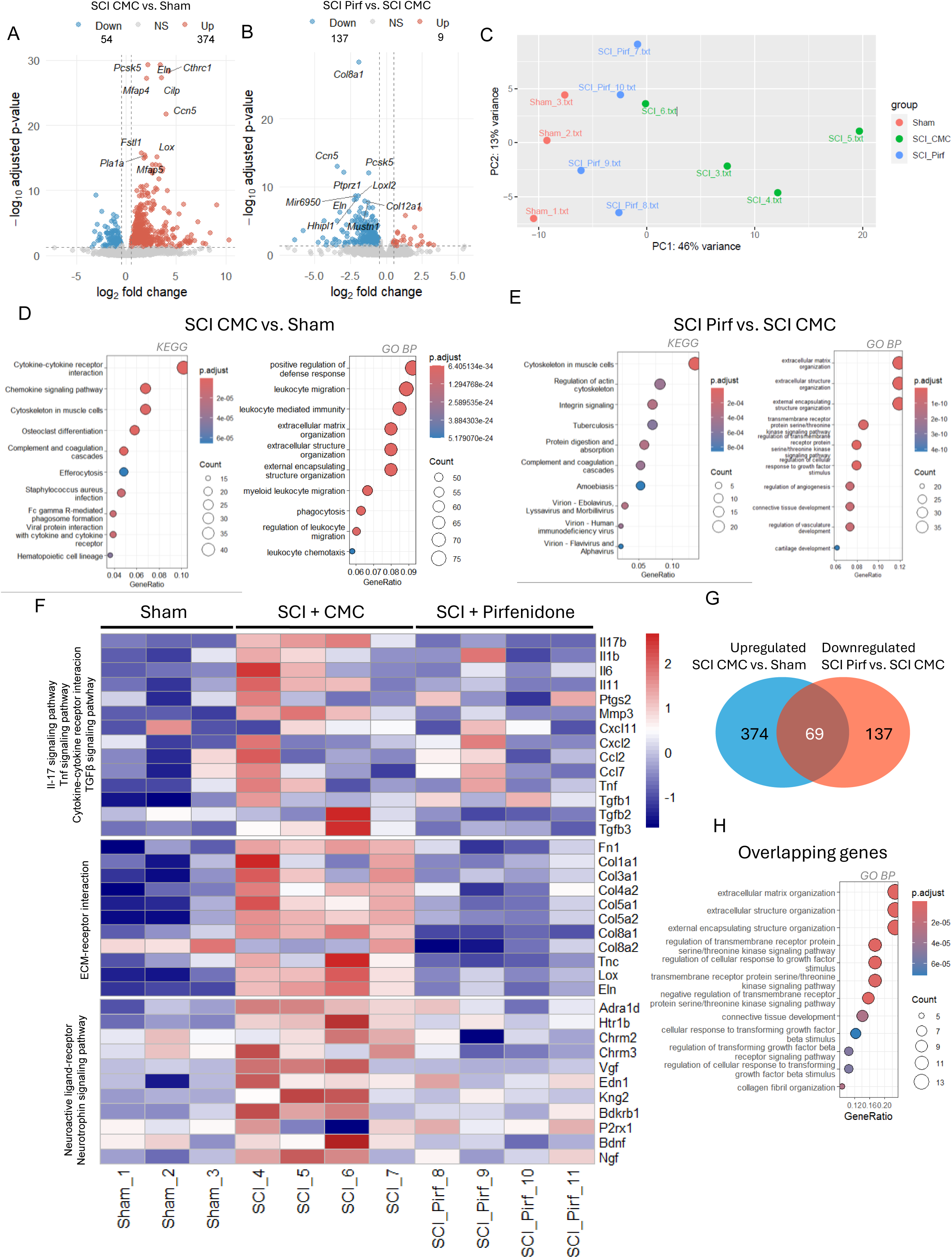
Transcriptomic profiling at d16 reveals persistent fibrotic and inflammatory signatures partially reversed by pirfenidone. Volcano plot of DEGs, SCI+CMC vs. sham (A) or SCI+Pirfenidone vs. SCI+CMC at d16 (B) (n=3 sham, n=4 SCI+CMC, n=4 SCI+Pirf). (C) PCA of all d16 RNAseq samples (sham, SCI+CMC, SCI+Pirfenidone). KEGG and GO BP pathway enrichment of DEGs, SCI+CMC vs. sham(D) or SCI+Pirf vs. SCI+CMC (E). (F) Heatmaps of representative of genes in *Il-17*, *Tnf-α* and TGFβ signaling pathways, and cytokine-cytokine receptor interaction (top), ECM-receptor interaction (middle), and neuroactive ligand-receptor/neurotrophin signaling (bottom) across sham, SCI+CMC, and SCI+Pirf groups. (G) Venn diagram of upregulated genes in SCI vs. sham mice and downregulated genes in SCI Pirf vs. SCI CMC mice. (H) GO BP pathway enrichment of overlapping DEGs in SCI vs. sham comparison, and downregulated genes in SCI Pirf vs. SCI CMC comparison.

**Table 5.**
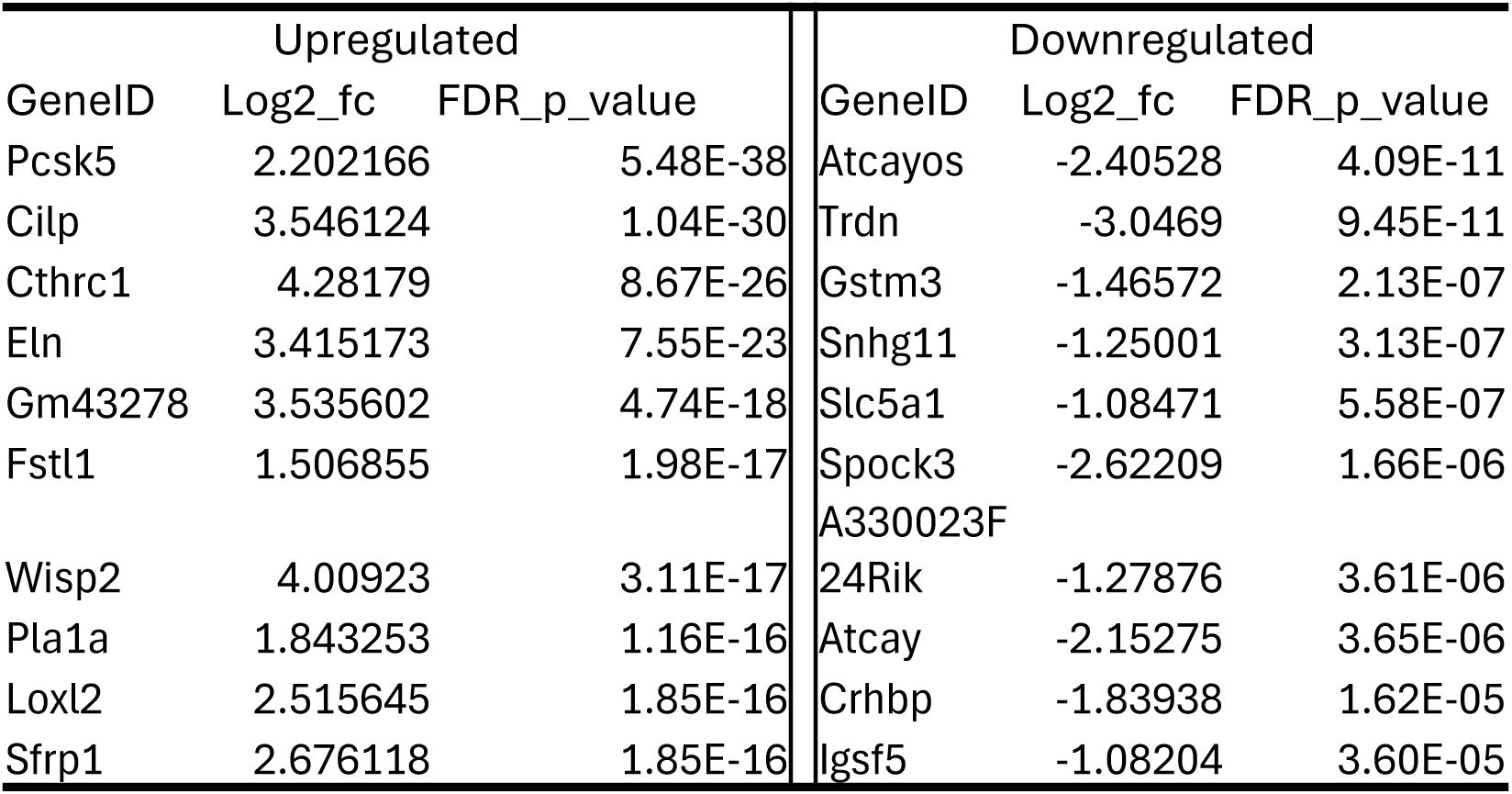
Top ten upregulated and downregulated genes between SCI + CMC and sham bladders 16 days after the surgery.

Comparison of transcriptional changes of bladders from SCI- and pirfenidone-treated mice revealed 146 DEGs with 9 upregulated and 137 downregulated (Fig. 6B, Supplementary Table 4). Table 6 shows the top 10 up- and downregulated genes in the bladders of SCI + CMC vs. SCI + pirfenidone mice. DEGs in the pirfenidone group revealed downregulation of pathways related to ECM organization, including many that were upregulated in the SCI vs. sham mice, specifically, *Fn1, Col4a2, Col5a1/2, Col8a1, and Lox* (Fig. 6F). Genes associated with pro-fibrotic and pro-inflammatory cascades like *Tgfb3*, *Il17b*, and macrophage markers *Arg1* and *Cd163* were also downregulated (Supplementary Table 5), suggesting modulation of pro-fibrotic signaling cascades. Additionally, genes related to neuronal function and contractility including *Bdnf, Chrm2, Actg2*, and *Myh2* were downregulated in pirfenidone-treated mice. A total of 69 genes that were upregulated in SCI vs. sham mice overlapped with those downregulated in SCI + pirfenidone vs. SCI mice (Fig. 6G). GO BP terms associated with this subset of genes suggested changes in ECM, regulation of the activity of serine threonine kinases, and TGFβ signaling suggesting that pirfenidone effectively targeted this signaling pathway and decreased ECM deposition. In summary, these transcriptomic analyses show that SCI induces a transition from acute inflammation to progressive ECM remodeling and fibrosis, while pirfenidone prevents SCI-associated pro-fibrotic transcriptional changes.

**Table 6.** Top ten upregulated and downregulated genes between SCI + Pirf and SCI + CMC bladders 16 days after the surgery.

| Upregulated |  |  | Downregulated |  |  |
| --- | --- | --- | --- | --- | --- |
| GeneID | Log2_fc | FDR_p_value | GeneID | Log2_fc | FDR_p_value |
| Gstm3 | 1.201883 | 2.67E-06 | Col8a1 | -1.85897 | 7.62E-20 |
| Trdn | 2.366513 | 3.81E-06 | Wisp2 | -3.57672 | 1.33E-13 |
| Rasgrf1 | 1.34833 | 0.000473 | Ptprz1 | -3.13235 | 1.33E-13 |
| Hhatl | 1.909963 | 0.000971 | Prex2 | -1.42483 | 1.65E-09 |
| Phlda2 | 1.350018 | 0.001192 | Actg2 | -1.2535 | 1.65E-09 |
| Ttr | 1.849511 | 0.001192 | Gm43278 | -2.5324 | 1.86E-09 |
| Ngb | 1.867633 | 0.00123 | Mir6950 | -2.22069 | 3.70E-09 |
| Gm28729 | 1.683396 | 0.004327 | Pcsk5 | -1.22124 | 9.56E-09 |
|  |  |  | Ltbp2 | -2.95312 | 1.44E-08 |
|  |  |  | F13a1 | -1.50479 | 1.70E-08 |

### Pirfenidone improves bladder function after SCI

To determine whether pirfenidone improved bladder function at d16, void spot assays were performed (Fig. 7C). Pirfenidone-treated mice showed a significant decrease in voiding events, compared to SCI mice (Fig. 7A). While total volume voided (total spot area) was not significantly different between groups (Fig 7B), the volume per void (measured as the average spot area) was decreased in SCI mice and increased in pirfenidone-treated and sham mice (Fig. 7D), suggesting that pirfenidone restores bladder capacity. While the ECM composition of the bladder after pirfenidone did reveal less collagen III, the assays reflecting total collagen composition did not show significant differences in pirfenidone-treated versus SCI mice. This suggested that preservation in bladder function could be due to improved neuronal function. Indeed, our transcriptomic analysis showed a downregulation of *Bdnf* and *Chrm2* and *Adra1d* in pirfenidone-treated compared to SCI mice, which suggested less axonal sprouting and/or less neurotransmission (17, 18). Taken together with the GO terms that suggested changes in vascularization in pirfenidone-treated mice, we examined vascular and neuronal architecture in the bladder. Whole bladder antibody labeling was performed to detect neuronal (TUJ1) and vascular (CD31) structures using an adapted iDISCO protocol and light sheet microscopy (Fig. 7C, second row). CD31 staining (green) revealed an increase in the vascularization of SCI bladders compared to sham, but the bladders treated with pirfenidone had less-developed vascular trees than vehicle-treated bladders (Fig 7C, second row). Concomitantly, CD31 protein levels appeared increased in bladders from SCI mice, while pirfenidone-treated mice showed less CD31 protein expression. These trends were observed in western blot, but the differences were not statistically significant in densitometric analysis (Fig. 3SC, D). Nerve structure by TUJ1 staining was better visualized using confocal microscopy. TUJ1 staining revealed long and continuous axonal processes in the bladders of sham mice which appeared disrupted and swollen in SCI mice. In contrast, pirfenidone treatment preserved neurons as shown by long and intact processes (Fig. 7C, third row). Interestingly, TUJ1 protein levels on western blot were decreased in SCI mice and a partial recovery was observed in the pirfenidone-treated mice (Fig. 3SB, D). This data suggests that early pirfenidone treatment prevents vascular and neuronal changes associated with SCI.

**Figure 7.**
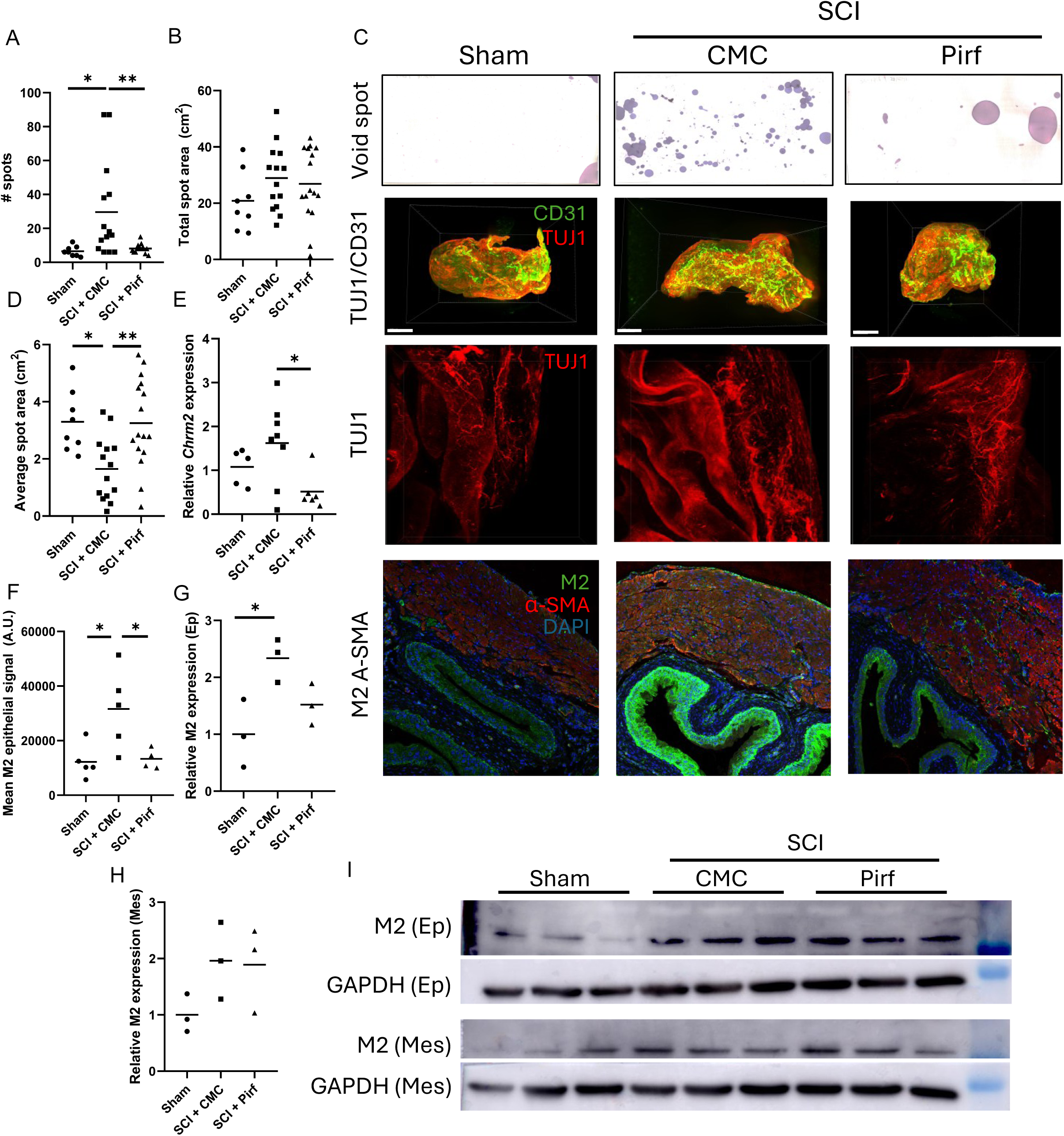
Pirfenidone improves voiding function and modulates uroepithelial muscarinic receptor 2, M2, expression at d16. (A) Number of void spots in sham, SCI+CMC, and SCI+Pirf mice (ANOVA; n=8 sham, n=14 SCI+CMC, n=16 SCI+Pirf). (B) Total void spot area for same groups. (C) Representative void spot assay images (top), whole-bladder TUJ1/CD31 protein expression shown by lightsheet microscopy (second row), scale bars: 1 mm; whole-bladder TUJ1 expression shown by confocal microscopy (third row), scale bars: 200 µm; and M2/α-SMA protein expression of bladder sections shown by confocal microscopy (bottom), scale bars: 200 µm. (D) Average void spot area. (E) Relative *Chrm2* mRNA expression by RT-qPCR. (F) Quantification of immunofluorescent signal quantification of M2 staining in the epithelium in arbitrary units. (G) Relative M2 protein expression in uroepithelial lysates (n=3/group). (H) Relative M2 protein expression in mesenchymal/detrusor lysates (not significant, n=3/group). (I) Representative western blots of M2 in epithelial (Ep) and mesenchymal (Mes) fractions with corresponding GAPDH loading controls.

To determine if the improved bladder function was due to changes in neurotransmitter sensitivity or signaling, we assessed muscarinic receptor expression. We found that the *Chrm2* gene that encodes the muscarinic receptor 2 was significantly upregulated in SCI mice but downregulated in pirfenidone-treated mice by RT-qPCR (Fig. 7E). There were no changes in *Chrm3* gene expression in SCI compared to sham mice (Fig. 3SA). The muscarinic acetylcholine receptor 2 (M2) is expressed in the uroepithelium and muscle wall and is upregulated in the denervated bladder (19). To examine whether M2 upregulation was global or layer-specific, M2 localization and abundance was assessed by immunofluorescence and western blot analysis. Interestingly, there was increased expression of the M2 receptor in the uroepithelium of SCI mice, while levels remained comparable among sham and pirfenidone-treated mice (Fig. 7C lower panel, F). No overt differences were detected in the muscle layer. Western blot analyses were performed on bladder lysates in which the uroepithelium (Ep) was separated from the lamina propria and muscle (Mes). Consistent with what was observed in the immunofluorescent labeling, there was an increase in M2 expression in the uroepithelium of SCI mice (Fig. 7G and I, M2 (Ep)) and a decrease in pirfenidone-treated mice. This pattern was not observed in the combined lamina propria and muscle samples (Fig. 7H). This decrease in M2 expression could reflect decreased cholinergic responsiveness within the uroepithelial layer that results in a decrease in the number of voiding events.

### Structural and functional changes elicited by pirfenidone are maintained at day 45

To determine whether the effects of short-term pirfenidone treatment were sustained, bladders were analyzed at day 45. Bladder weights were increased in all SCI mice but were lower in the pirfenidone-treated compared with SCI mice (Fig. 8A). Histological examination by H&E staining revealed no obvious differences between SCI and pirfenidone-treated mice (Fig. 8B, upper panel). However, Masson trichrome staining suggested reduced collagen deposition in the pirfenidone-treated mice (Fig. 8C). M2 expression was examined at d45 by immunofluorescence. A similar pattern was observed: uroepithelial M2 expression was elevated in SCI mice compared to sham and pirfenidone-treated mice, while the level of M2 protein in sham and pirfenidone-treated mice appeared comparable (Fig. 8B third row, and 8D). These findings also correlated with changes in voiding patterns evaluated by the void spot assay. The number of voiding spots was increased in SCI mice compared to pirfenidone-treated and sham mice (Fig. 8B bottom panels). Similarly, the average voiding volume was increased in pirfenidone-treated mice compared to SCI mice at d45, as was seen at d16 (Fig. 8F). These results suggest that short and preventive pirfenidone treatment can dampen the inflammation and extracellular matrix changes observed in SCI and improve bladder function by modulating M2 expression within the uroepithelium.

**Figure 8.**
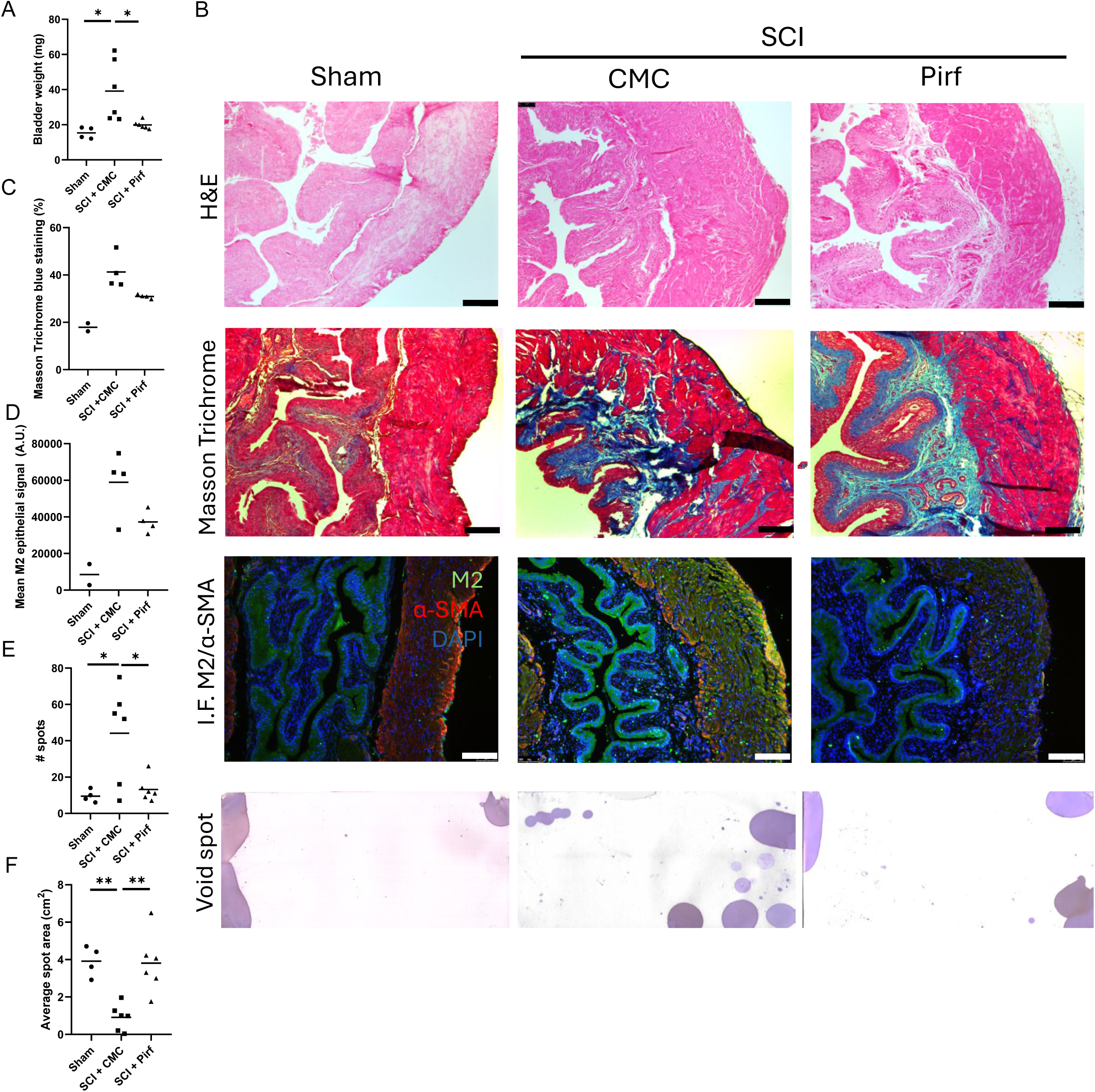
Pirfenidone’s effects on bladder fibrosis and voiding function are sustained at d45. (A) Bladder weight at d45 in sham, SCI+CMC, and SCI+Pirf groups (ANOVA, n=2 sham, n=4 SCI+CMC and SCI+Pirf). (B) Representative bladder sections stained by hematoxylin and eosin (top), Masson trichrome (second row), and immunofluorescent detection of M2/α-SMA protein expression (third row) from bladder sections and representative void spot assays (fourth row). (C) Quantification of Masson trichrome, blue-stained area (% of total area). Scale bars, 150 µm. (D) Quantification immunofluorescent signal of M2 staining in the epithelium in arbitrary units. (E) Number of void spots. (F) Average void spot area. (ANOVA, n=4 sham, n=6 SCI+CMC and SCI+Pirf).

## Discussion

We used a well-established contusion model of spinal cord injury in the mouse (2, 6) that simulates traumatic injury in humans. In humans, during spinal shock, the bladder is inflamed and hypotonic and requires catheterization for drainage. When spinal shock ends, detrusor overactivity is observed during urine storage. In contrast, during voiding, ineffective bladder emptying occurs due to detrusor-sphincter dyssynergia (20). These impairments result in distension and ischemia that progresses to fibrosis. Our mouse contusion model showed this progression but in a shorter timespan. During the period of spinal shock from day 0-7, the mice underwent manual compression of the bladder. Early during spinal shock at day 2, bladders from SCI mice exhibited neutrophil infiltration and upregulation of inflammatory pathways with increased cell proliferation in the uroepithelium, the lamina propria and the muscle layer. By day 7 after spinal cord injury when spontaneous voiding had resumed, the muscle layer was enlarged, indicating progression to detrusor-sphincter dyssynergia. Muscle hypertrophy was accompanied by persistent upregulation of inflammatory pathways and an increase in expression of extracellular matrix genes. Comparison of gene expression at day 2 and 7, revealed that TGFβ signalling was upregulated and central to the progression from inflammation to fibrosis (21). To target inflammation and fibrosis, we treated mice with pirfenidone from day 2 to day 7 and noted a marked and sustained improvement in bladder function up to 45 days post-injury. Pirfenidone decreased gene expression associated with inflammation, fibrosis, and neural signaling. The expression of muscarinic receptor 2 was decreased in the uroepithelium suggesting that pirfenidone improves bladder function by blunting acetylcholine-mediated neural signaling and reflex voiding.

Our transcriptomic and histological analysis revealed that neutrophils were recruited in the first 48 hours. Activated neutrophils contribute to dysfunction of the uroepithelium and release neutrophilic extracellular traps, reactive oxygen species, and cytokines that sustain inflammation and attract additional immune cell lineages including macrophages, dendritic cells, and T cells (Fig. 3S) (22). This shares similarities to other models of the neurogenic bladder. Dong et al. investigated the transcriptomic changes in the female rat bladder after complete transection of the spinal cord at d3, d7 and d25 and also found an early strong immune signature with ECM remodeling (23). Their analysis of all time points revealed 10 genes as potential hub genes including *Serpine1, Mmp3, Fn1* and *Edn1* that were also strongly expressed in our dataset. Similarly, ECM remodeling was also observed in a study that compared the bladder transcriptomes after complete transection of the spinal cord versus partial bladder outlet obstruction in female mice at 7–8 weeks after intervention. Notably, the TGFβ signaling pathway was a common signature in both bladder diseases (24). After severing spinal nerves in female rats to induce a neurogenic bladder, Qi et al. (2022) found that hypoxia, inflammation, and TNF-α signaling provoked ECM remodeling (25). More recently, a transcriptomic study from day 4 to day 28 after complete spinal transection in female rats also identified a temporal transition from acute inflammatory pathways, including leukocyte migration and cytokine production, to a chronic fibrotic program enriched for extracellular matrix organization, fibroblast proliferation and TGFβ signaling (26). Based on our findings and others, the neurogenic bladder is initiated by a strong immune reaction resulting in activation of TGFβ signaling during the transition between inflammation and fibrosis. Stromal and urothelial cells can secrete TGFβ in response to stretching in an injury-repair response and/or to cytotoxic exposures (27). Taken together, the immune infiltration observed at d2, and the events occurring between d2 and d7 are crucial to the development of bladder dysfunction in SCI.

Most studies of bladder dysfunction from SCI have focused on fibrosis as the major cause of impaired expansion and contractility. However, inflammation also drives neuronal re-arrangement as well as ECM deposition, and both could contribute to aberrant bladder contractility. In the early shock phase, the lack of communication with the central nervous system as well as the oxidative stress associated with inflammation result in emergence of autonomous circuits that control micturition. NGF and BDNF are secreted by the bladder to re-establish innervation as observed in our dataset at day 2. At day 7 and 16, BDNF was the major neurotrophin identified. BDNF has been shown to significantly upregulate genes involved in cholinergic transmission in the neurogenic bladder including cholinergic receptor muscarinic 2, *Chrm2*, that encodes the M2 muscarinic acetylcholine receptor 2 (28). Consistent with the function of BDNF, we observed significant upregulation of M2 in our model. Kwon et al. also observed marked upregulation of M2 and M3 mRNA transcripts after SCI in a mouse model of spinal cord transection (29). M3 is the main muscarinic receptor that mediates bladder contraction, however in the neurogenic bladder, M2 is upregulated and contributes to dysfunction (30–32). The paradigm of a pro-inflammatory environment that can upregulate muscarinic receptor expression to modify an organ’s neural signaling is also observed in the lungs. In pulmonary fibroblasts and epithelial cells, upregulation of muscarinic receptor expression is postulated to enhance clearance of bacteria by mediating the contraction of airway smooth muscle cells (33).

Pirfenidone is an anti-fibrotic agent that is a treatment for idiopathic pulmonary fibrosis. Pirfenidone inhibits TGFβ, TNF-α and PDGF signaling pathways to decrease inflammation and limit ECM deposition. Here, pirfenidone was evaluated as a potential modulator of bladder inflammation early after SCI, as opposed to its use after injury is fully established. Indeed, pirfenidone was able to decrease transcription of cytokines *Tgfb2/3*, *Il1b*, *Il6* and *Tnf-α* in our model. The prominence of *Tgfb2/3* and not *Tgfb1* in this process is consistent with pro-inflammatory signaling events in a genetic mouse model of cardiomyopathy (34). Pirfenidone attenuated bladder overactivity by decreasing the number of voiding events and increasing the volume per void, consistent with improved bladder function. This implies an interplay between inflammation, TGFβ signaling and voiding, where pirfenidone can positively intervene. Pirfenidone also appears to blunt the increase in uroepithelial cellularity after SCI(35, 36) although the role of the uroepithelium in bladder dysfunction is unclear. Our results are consistent with Ko et al. who reported improved bladder function in a rat model with an underactive bladder. They started treatment with pirfenidone at a later timepoint, 6 weeks after crushing injury to the pelvic ganglia (13). In another rat model of partial bladder outlet obstruction, daily pirfenidone administration was initiated one week after obstruction and continued for 5 weeks (14) and resulted in a decrease in ECM deposition. Our model is one of detrusor overactivity and this may be related to the endpoint selected for our study. The neurogenic bladder from SCI in rodents begins as detrusor overactivity in the sub-acute phase of the injury (1-4 weeks) and progresses to a detrusor underactivity in the chronic phase of the injury. ECM deposition contributes to detrusor underactivity in the chronic phase, whereas the contribution of ECM deposition in the subacute phase of the injury is less clear.

Our oral pirfenidone regime is the shortest and earliest treatment reported with noticeable results in bladder function. This is important given that long-term treatment with pirfenidone can result in significant gastrointestinal side effects that limit adherence. Zhang et al. demonstrated that intravesical instillation of pirfenidone using polydopamine nanoparticles (PDA) as a delivery method during the early shock phase was also effective at preserving bladder function and histology in a rat model with complete transection of the spinal cord. They demonstrated decreased immune infiltration after pirfenidone treatment, with pronounced effects on neutrophils and monocyte recruitment that prevented ECM accumulation. Zhang et al. contrasted the effect of intravesical particle instillation with oral pirfenidone administration starting from day 0 and continuing up to day 28 and reported a much milder improvement in bladder function with oral pirfenidone compared to our findings. We showed a decrease in bladder weight of approximately 40% from pirfenidone treatment compared to ∼12% in the Zhang study. They also noted that PDA particles themselves induce an antioxidant effect that may synergize with pirfenidone to prevent SCI-induced bladder dysfunction. While this was not examined in our studies, pirfenidone may also improve bladder function through its antioxidant effects. In both our study and Zhang et al., it is possible that pirfenidone is limiting the spinal cord injury itself with secondary benefits on bladder function, given its anti-inflammatory and anti-oxidant effects.

Beyond its use in idiopathic pulmonary fibrosis, pirfenidone has been studied in a number of human trials for acute lung injury, lung cancer, sarcoidosis, cirrhosis and chronic kidney disease (https://clinicaltrials.gov/search?term=pirfenidone&viewType=Card&page=1). Thus far, its use in bladder disease is limited. In a phase II randomized, double-blind human study in secondary progressive multiple sclerosis, Walker et al. reported a marked improvement in bladder dysfunction including urinary urgency, incontinence and need for catheterization in the pirfenidone group as compared to placebo (40.0% improved on PFD vs 16.7% on placebo).

In conclusion, the mouse model of spinal contusion recapitulates the molecular and cellular events observed in neurogenic bladder in humans. Short-term treatment with pirfenidone ameliorates bladder dysfunction when administered early after spinal cord contusion by decreasing inflammation and fibrosis and limiting upregulation of M2 receptors in the uroepithelium.

---

Figure S1. Transcriptomic differences and neutrophil invasion in mouse bladders 2 or 7 days after surgery. (A) Sample-to-sample distance analyses from sham and SCI samples subjected to bulk RNA-sequencing. Transcript reads were analyzed with DESeq2 and distances were calculated from variance-stabilized transformed data. Gradient ranges from low distances (dark blue/blue, high similarity) to high distances (light blue/white, indicating greater dissimilarity). Axes are ordered via hierarchical clustering to group samples with similar transcription patterns. (B) Representative bladder sections showing histology with hematoxylin and eosin staining, protein expression of Ly6G and Ki67, and the TUNEL assay (bottom row) of sham, SCI, and SCI neutrophilic node regions (n=5 sham, n=5 SCI). Dotted lines outline nodes of polymorphonuclear cell aggregates. Arrowheads indicate filament-like TUNEL signal. Scale bars: 150 µm, inset: 50 µm. (C) Sample-to-sample distance analyses from sham and SCI samples subjected to bulk RNA-sequencing at d7. (D) KEGG terms of overlapping genes between d2 and d7 analyses. Circle size indicates number of genes present from each term. Color scale indicates p-value (-log10-FDR corrected) (E) Ly6G immunohistochemistry of sham and SCI bladder sections at d7. Scale bars: 150 µm

Figure S2. Time-dependent transcriptomic remodeling of the bladder following spinal cord injury (SCI). (A) Principal component analysis (PCA) of variance-stabilized RNA-sequencing data from sham and SCI bladders collected at 2 (d2), 7 (d7), and 16 (d16) days post-injury. Samples cluster according to both injury status and timepoint, demonstrating progressive transcriptional changes following SCI. (B) Heatmap of genes identified as significantly differentially expressed by likelihood ratio test (LRT; DESeq2), genes highlighted vary in function of injury and time. Gene expression values are row-scaled (Z-score), and genes are grouped into co-expression modules based on hierarchical clustering. Functional annotations of representative modules include cell cycle, inflammatory response, mast cell response, muscle hypertrophy, protein maturation, and T cell differentiation. Sample annotations indicate timepoint and experimental group. (C) Heatmap of immune cell signature scores inferred from transcriptomic data, showing relative enrichment of neutrophil, monocyte, macrophage, dendritic cell, T cell, and B cell-associated associated gene. Color scales represent standardized expression (B) or immune cell signature scores (C). Replicates are shown individually.

Figure 3S. Analysis of neuro-vascular proteins in response to SCI and pirfenidone treatment. (A) Relative mRNA expression of *Chrm3*. Gene expression was quantified by RT-qPCR and normalized to housekeeping gene *Gapdh*. (B, C) Densitometric analysis of relative protein expression of TUJ1 (B) and CD31 (C) in bladders from Sham, SCI treated with vehicle (SCI+CMC), and SCI treated with pirfenidone (SCI+Pirf) mice. (D) Representative immunoblot showing protein expression of the endothelial marker CD31 and the neuronal marker TUJ1 in bladder tissue from Sham, SCI+CMC, and SCI+Pirf mice. GAPDH was used as a loading control. Individual data points represent biological replicates, and horizontal lines indicate the mean (ANOVA, n=3 sham, n=5 SCI+CMC, n=4 SCI+Pirf). **: p < 0.01.

## Acknowledgements

We thank Dr. L. O’ Brien for helpful advice and comments on lightsheet microscopy.

## Disclosures

### Disclaimers

The content is solely the responsibility of the authors and does not necessarily represent the official views of the RI MUHC.

## Author contributions

Conceived and designed research: CAIA, IRG, SD. Performed experiments: CAIA, VM, LC. Analyzed data: CAIA, LR, AB. Interpreted results of experiments: CAIA, IRG, SD. Prepared figures: CAIA. Drafted manuscript: CAIA. Edited revised manuscript: IRG, WK, JJ, CAIA, VM. Approved final version of manuscript: CAIA, VM, LC, LR, AB, JJ, WK, SD, IRG.

