## Supplemental Tables for "Early Treatment with Oral Pirfenidone Improves Bladder Function after Contusive Spinal Cord Injury in Mice"

Fig. S1

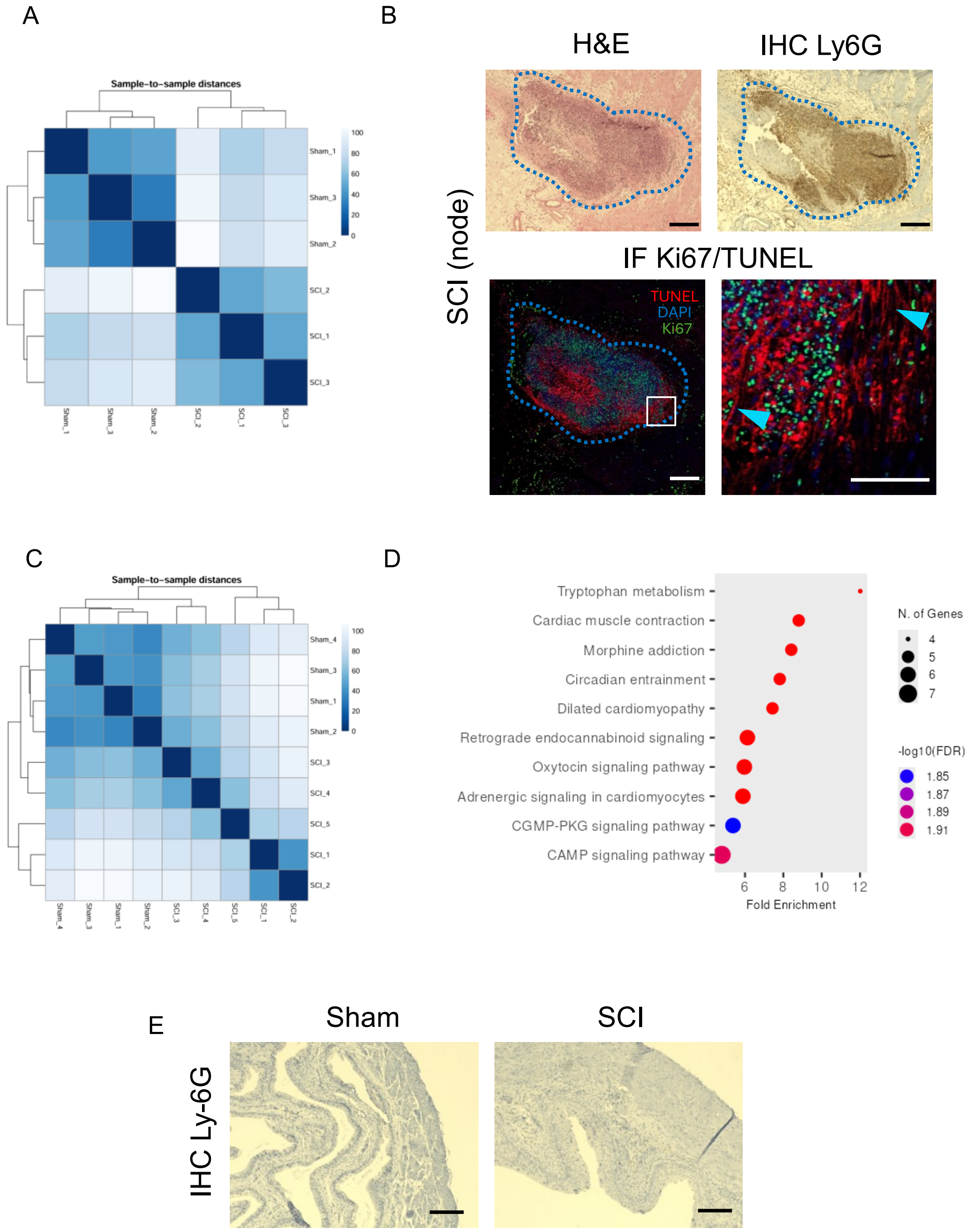
A
Sample-to-sample distances
Sham_1
Sham_3
Sham_2
SCI_2
SCI_1
SCI_3
100
80
60
40
20
0
B
H&E
IHC Ly6G
SCI (node)
IF Ki67/TUNEL
TUNEL
DAPI
Ki67
C
Sample-to-sample distances
Sham_4
Sham_3
Sham_1
Sham_2
SCI_3
SCI_4
SCI_5
SCI_1
SCI_2
D
Tryptophan metabolism
Cardiac muscle contraction
Morphine addiction
Circadian entrainment
Dilated cardiomyopathy
Retrograde endocannabinoid signaling
Oxytocin signaling pathway
Adrenergic signaling in cardiomyocytes
CGMP-PKG signaling pathway
CAMP signaling pathway
6
8
10
12
Fold Enrichment
N. of Genes
4
5
6
7
-log10(FDR)
1.85
1.87
1.89
1.91
E
Sham
SCI
IHC Ly-6G

Fig. S2

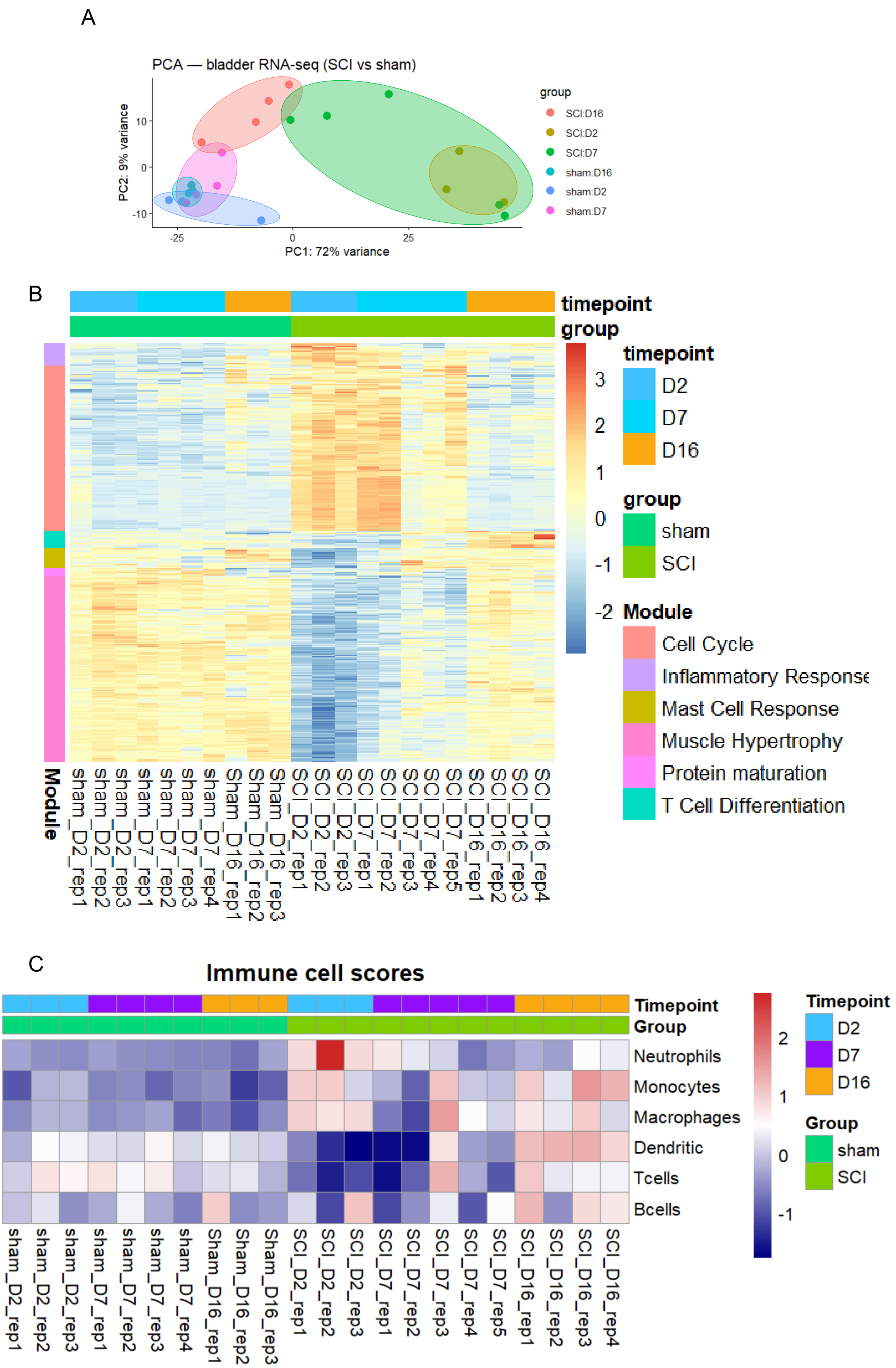
A
PCA — bladder RNA-seq (SCI vs sham)
PC1: 72% variance
PC2: 9% variance
group
SCI:D16
SCI:D2
SCI:D7
sham:D16
sham:D2
sham:D7
B
timepoint
group
D2
D7
D16
sham
SCI
Module
Cell Cycle
Inflammatory Response
Mast Cell Response
Muscle Hypertrophy
Protein maturation
T Cell Differentiation
sham_D2_rep1
sham_D2_rep2
sham_D2_rep3
sham_D7_rep1
sham_D7_rep2
sham_D7_rep3
sham_D7_rep4
Sham_D16_rep1
Sham_D16_rep2
Sham_D16_rep3
SCI_D2_rep1
SCI_D2_rep2
SCI_D2_rep3
SCI_D7_rep1
SCI_D7_rep2
SCI_D7_rep3
SCI_D7_rep4
SCI_D7_rep5
SCI_D16_rep1
SCI_D16_rep2
SCI_D16_rep3
SCI_D16_rep4
C
Immune cell scores
Timepoint
Group
Neutrophils
Monocytes
Macrophages
Dendritic
Tcells
Bcells

Fig. S3

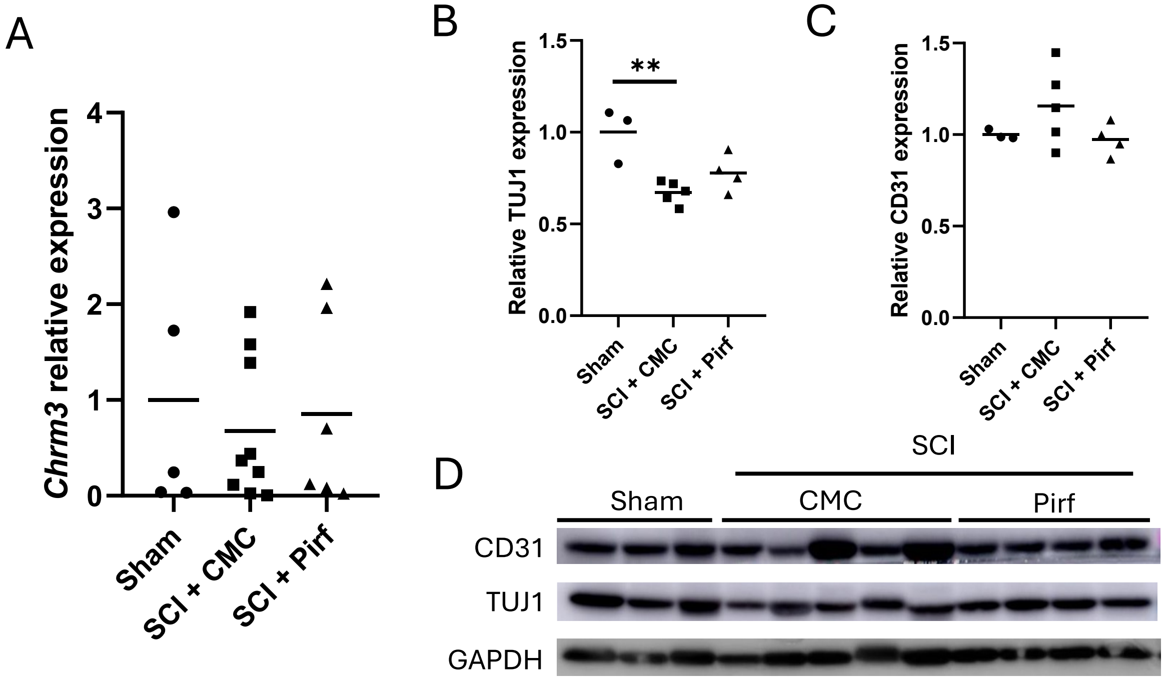

A
Chrm3 relative expression
4
3
2
1
0
Sham
SCI + CMC
SCI + Pirf
B
Relative TUJ1 expression
1.5
1.0
0.5
0.0
**
Sham
SCI + CMC
SCI + Pirf
C
Relative CD31 expression
1.5
1.0
0.5
0.0
Sham
SCI + CMC
SCI + Pirf
D
SCI
Sham
CMC
Pirf
CD31
TUJ1
GAPDH

Supplemental Table 1. d2 DEGs

| GeneID | Log2 fc | SE | p-value | FDR p-value |
|---|---|---|---|---|
| Serpine1 | 6.621968 | 0.226899 | 3.04E-187 | 6.93E-183 |
| Ptgs2 | 4.973526 | 0.233786 | 1.98E-100 | 2.26E-96 |
| Gm15270 | 5.519216 | 0.272625 | 3.96E-91 | 3.01E-87 |
| Wisp1 | 3.632947 | 0.191561 | 3.33E-80 | 1.90E-76 |
| Thbs1 | 3.048535 | 0.178384 | 1.77E-65 | 8.07E-62 |
| Fam150a | 8.28091 | 0.48517 | 2.57E-65 | 9.76E-62 |
| Ptx3 | 6.375357 | 0.374464 | 5.33E-65 | 1.74E-61 |
| Adamts4 | 5.246155 | 0.310384 | 4.34E-64 | 1.24E-60 |
| Tnfrsf12a | 4.856062 | 0.292947 | 1.03E-61 | 2.61E-58 |
| Socs3 | 4.960543 | 0.302962 | 2.96E-60 | 6.75E-57 |
| Ctgf | 5.542263 | 0.342338 | 5.99E-59 | 1.24E-55 |
| Hist1h2ap | 5.731842 | 0.358566 | 1.61E-57 | 3.07E-54 |
| Bmper | 5.02545 | 0.316148 | 6.77E-57 | 1.19E-53 |
| Bdkrb1 | 6.411508 | 0.404019 | 1.03E-56 | 1.68E-53 |
| Chil3 | 6.532391 | 0.431702 | 1.00E-51 | 1.52E-48 |
| Kcnc4 | -6.04914 | 0.405957 | 3.25E-50 | 4.63E-47 |
| Ier3 | 2.356341 | 0.160425 | 7.68E-49 | 9.73E-46 |
| Cemip | 4.914224 | 0.334502 | 7.34E-49 | 9.73E-46 |
| Mamdc2 | -4.90432 | 0.335712 | 2.47E-48 | 2.97E-45 |
| Nes | 2.92896 | 0.200574 | 2.69E-48 | 3.07E-45 |
| Gm44466 | 6.285416 | 0.430843 | 3.32E-48 | 3.60E-45 |
| Fhl3 | 3.678756 | 0.252596 | 4.77E-48 | 4.95E-45 |
| Dusp10 | 2.572147 | 0.176888 | 6.65E-48 | 6.60E-45 |
| Gadd45g | 2.814562 | 0.195275 | 4.27E-47 | 4.05E-44 |
| Lox | 4.387867 | 0.312091 | 6.73E-45 | 6.14E-42 |
| Bdnf | 4.537321 | 0.32715 | 9.73E-44 | 8.54E-41 |
| Trib1 | 2.234822 | 0.161937 | 2.53E-43 | 2.14E-40 |
| Ccl2 | 5.057653 | 0.367074 | 3.44E-43 | 2.81E-40 |
| Cyr61 | 2.948823 | 0.215653 | 1.45E-42 | 1.14E-39 |
| Ccl7 | 4.94383 | 0.361753 | 1.61E-42 | 1.23E-39 |
| Hspb7 | 2.928336 | 0.215271 | 3.84E-42 | 2.83E-39 |
| Sv2b | -3.54745 | 0.263234 | 2.15E-41 | 1.53E-38 |
| Has1 | 5.485181 | 0.409039 | 5.29E-41 | 3.66E-38 |
| Egr2 | 4.609185 | 0.344236 | 6.95E-41 | 4.66E-38 |
| Lrrc32 | 2.636977 | 0.198159 | 2.10E-40 | 1.37E-37 |
| Uchl1 | -2.63878 | 0.200567 | 1.56E-39 | 9.61E-37 |
| Akap2 | 2.753204 | 0.20926 | 1.55E-39 | 9.61E-37 |
| Pakap | 2.73317 | 0.209537 | 6.89E-39 | 4.14E-36 |
| Ccl9 | 2.93338 | 0.225112 | 8.18E-39 | 4.78E-36 |
| Fam212b | -2.66882 | 0.20484 | 8.39E-39 | 4.78E-36 |
| Spock3 | -4.25782 | 0.328786 | 2.35E-38 | 1.31E-35 |
| Wnk2 | -3.90339 | 0.306554 | 3.87E-37 | 2.10E-34 |

| | | | | |
|---|---|---|---|---|
| Hspb1 | 2.483881 | 0.196054 | 8.74E-37 | 4.64E-34 |
| Hrh3 | -4.91972 | 0.390907 | 2.54E-36 | 1.32E-33 |
| Foxs1 | 4.670645 | 0.371268 | 2.71E-36 | 1.37E-33 |
| Syt14 | -4.13758 | 0.329355 | 3.39E-36 | 1.68E-33 |
| Ung | 3.221026 | 0.259477 | 2.21E-35 | 1.07E-32 |
| Has2 | 3.283638 | 0.265496 | 3.90E-35 | 1.85E-32 |
| Ccl6 | 3.41426 | 0.277439 | 8.37E-35 | 3.90E-32 |
| Il11 | 4.754358 | 0.387087 | 1.13E-34 | 5.14E-32 |
| Hist1h2ao | 6.70295 | 0.552924 | 8.00E-34 | 3.58E-31 |
| Nxph3 | 2.564391 | 0.213084 | 2.34E-33 | 1.03E-30 |
| F3 | 2.481799 | 0.206273 | 2.42E-33 | 1.04E-30 |
| Ipcef1 | -4.31121 | 0.358609 | 2.72E-33 | 1.15E-30 |
| Nkd2 | -2.40214 | 0.200753 | 5.38E-33 | 2.23E-30 |
| Smad9 | -3.40931 | 0.286819 | 1.39E-32 | 5.66E-30 |
| Il6 | 6.815453 | 0.574491 | 1.83E-32 | 7.33E-30 |
| Tnc | 2.537665 | 0.214192 | 2.21E-32 | 8.71E-30 |
| Edn1 | 2.389022 | 0.201865 | 2.58E-32 | 9.98E-30 |
| Gm29233 | 2.966243 | 0.251122 | 3.38E-32 | 1.29E-29 |
| Maff | 2.091365 | 0.177945 | 6.82E-32 | 2.55E-29 |
| Gdf15 | 5.693301 | 0.484652 | 7.30E-32 | 2.69E-29 |
| Ddx26b | -2.25381 | 0.192477 | 1.14E-31 | 4.13E-29 |
| Rtn4rl2 | 3.547466 | 0.306911 | 6.68E-31 | 2.38E-28 |
| Gbp5 | -3.12603 | 0.27243 | 1.77E-30 | 6.12E-28 |
| Akap6 | -3.00111 | 0.261659 | 1.88E-30 | 6.39E-28 |
| Gm37804 | 2.526962 | 0.2241 | 1.72E-29 | 5.70E-27 |
| Nfil3 | 3.081519 | 0.273258 | 1.71E-29 | 5.70E-27 |
| Noct | 2.664172 | 0.236592 | 2.05E-29 | 6.69E-27 |
| Pcsk5 | 3.101077 | 0.275735 | 2.41E-29 | 7.73E-27 |
| Mcm3 | 3.193767 | 0.284495 | 3.04E-29 | 9.62E-27 |
| Kit | -3.42391 | 0.305038 | 3.09E-29 | 9.65E-27 |
| Gm26885 | 2.18744 | 0.195049 | 3.45E-29 | 1.06E-26 |
| Xirp1 | 3.024511 | 0.271444 | 7.81E-29 | 2.37E-26 |
| Dbp | -4.17748 | 0.377267 | 1.70E-28 | 5.02E-26 |
| Clec4d | 4.961014 | 0.449594 | 2.61E-28 | 7.62E-26 |
| Rgs16 | 2.885202 | 0.262814 | 4.87E-28 | 1.41E-25 |
| Timeless | 2.798275 | 0.256148 | 8.81E-28 | 2.48E-25 |
| Snai1 | 2.318747 | 0.213005 | 1.35E-27 | 3.74E-25 |
| B4galt5 | 2.030903 | 0.187214 | 2.04E-27 | 5.60E-25 |
| Dusp5 | 3.041813 | 0.282522 | 4.95E-27 | 1.34E-24 |
| Tcap | -5.06918 | 0.47141 | 5.72E-27 | 1.54E-24 |
| Ccne1 | 3.103269 | 0.289301 | 7.62E-27 | 2.02E-24 |
| Ciart | -2.72953 | 0.255573 | 1.26E-26 | 3.31E-24 |
| Kcng1 | -3.76823 | 0.358882 | 8.64E-26 | 2.22E-23 |
| Rab17 | -4.62081 | 0.440915 | 1.07E-25 | 2.70E-23 |

| | | | | |
|---|---|---|---|---|
| Kcnn2 | -4.16432 | 0.397999 | 1.28E-25 | 3.20E-23 |
| Msn | 2.43744 | 0.233263 | 1.48E-25 | 3.66E-23 |
| Gnao1 | -2.0039 | 0.191936 | 1.62E-25 | 3.93E-23 |
| Adam12 | 3.049403 | 0.292559 | 1.94E-25 | 4.66E-23 |
| Mcm2 | 2.717158 | 0.260982 | 2.20E-25 | 5.23E-23 |
| Myoc | -4.06906 | 0.390882 | 2.23E-25 | 5.25E-23 |
| Rgs5 | -3.07135 | 0.29713 | 4.81E-25 | 1.12E-22 |
| Slc9a2 | -3.09416 | 0.301537 | 1.05E-24 | 2.43E-22 |
| Tpm4 | 2.087721 | 0.203675 | 1.18E-24 | 2.67E-22 |
| Tgfb3 | 2.71919 | 0.265687 | 1.39E-24 | 3.10E-22 |
| Ms4a6d | 3.226627 | 0.315548 | 1.52E-24 | 3.38E-22 |
| Nasp | 2.254651 | 0.220691 | 1.68E-24 | 3.64E-22 |
| Mir6950 | 2.378376 | 0.23293 | 1.78E-24 | 3.82E-22 |
| Mcm6 | 2.784343 | 0.273515 | 2.44E-24 | 5.15E-22 |
| Loxl2 | 2.548504 | 0.250462 | 2.56E-24 | 5.35E-22 |
| Fst | 3.33625 | 0.328459 | 3.08E-24 | 6.32E-22 |
| Serpina3m | 4.270365 | 0.420427 | 3.08E-24 | 6.32E-22 |
| Dock3 | -2.25344 | 0.22205 | 3.37E-24 | 6.86E-22 |
| Gria1 | -4.73076 | 0.468134 | 5.22E-24 | 1.04E-21 |
| Msr1 | 2.740685 | 0.271899 | 6.79E-24 | 1.35E-21 |
| Il4ra | 2.377387 | 0.237338 | 1.29E-23 | 2.53E-21 |
| Itga8 | -2.66456 | 0.26778 | 2.51E-23 | 4.85E-21 |
| Pim1 | 2.304028 | 0.232 | 3.05E-23 | 5.84E-21 |
| Hells | 4.24798 | 0.428117 | 3.32E-23 | 6.32E-21 |
| Fbxo32 | -4.08287 | 0.414008 | 6.09E-23 | 1.15E-20 |
| Ccr1 | 2.764392 | 0.281137 | 8.12E-23 | 1.51E-20 |
| Mt2 | 2.08721 | 0.212468 | 8.91E-23 | 1.64E-20 |
| Flnc | 2.506193 | 0.256493 | 1.50E-22 | 2.72E-20 |
| Gdf6 | 2.868062 | 0.293758 | 1.62E-22 | 2.90E-20 |
| Hk2 | 2.38386 | 0.245509 | 2.74E-22 | 4.84E-20 |
| Lrp8 | 3.950029 | 0.407088 | 2.92E-22 | 5.13E-20 |
| Tle2 | -2.33159 | 0.241459 | 4.62E-22 | 7.99E-20 |
| Cbx7 | -2.01223 | 0.208669 | 5.25E-22 | 9.01E-20 |
| Tenm2 | -3.8959 | 0.404161 | 5.45E-22 | 9.27E-20 |
| Tex15 | -2.16174 | 0.224338 | 5.63E-22 | 9.45E-20 |
| Tagln2 | 2.167741 | 0.224963 | 5.63E-22 | 9.45E-20 |
| Ereg | 4.076154 | 0.423708 | 6.57E-22 | 1.09E-19 |
| Arg1 | 7.844143 | 0.815482 | 6.65E-22 | 1.10E-19 |
| Galr2 | -4.02619 | 0.418993 | 7.31E-22 | 1.19E-19 |
| 11-Sep | 2.021371 | 0.210343 | 7.26E-22 | 1.19E-19 |
| Wdhd1 | 2.915868 | 0.303803 | 8.16E-22 | 1.32E-19 |
| Bcat1 | 2.808848 | 0.294346 | 1.39E-21 | 2.24E-19 |
| Inhba | 3.177207 | 0.333407 | 1.58E-21 | 2.52E-19 |
| Gm26586 | 2.06091 | 0.217541 | 2.70E-21 | 4.25E-19 |

| | | | | |
|---|---|---|---|---|
| C5ar1 | 2.128012 | 0.225046 | 3.20E-21 | 5.00E-19 |
| Chaf1a | 2.798723 | 0.29621 | 3.44E-21 | 5.34E-19 |
| Filip1l | 2.139859 | 0.227887 | 6.00E-21 | 9.19E-19 |
| Dio3 | 4.461 | 0.476624 | 8.00E-21 | 1.22E-18 |
| Cdt1 | 2.75518 | 0.294893 | 9.37E-21 | 1.41E-18 |
| Ogn | -2.71225 | 0.290657 | 1.04E-20 | 1.57E-18 |
| Dok7 | -2.45319 | 0.262958 | 1.07E-20 | 1.59E-18 |
| Rgs11 | -2.16921 | 0.233239 | 1.40E-20 | 2.07E-18 |
| Hcn1 | -3.03951 | 0.327451 | 1.66E-20 | 2.41E-18 |
| Gm13446 | -3.25321 | 0.352414 | 2.68E-20 | 3.77E-18 |
| Ctla2b | 2.23814 | 0.243006 | 3.25E-20 | 4.55E-18 |
| Tc2n | -2.23147 | 0.242723 | 3.80E-20 | 5.29E-18 |
| Gm9817 | 2.464509 | 0.268384 | 4.20E-20 | 5.81E-18 |
| 6030408B1 | 2.80341 | 0.305957 | 5.06E-20 | 6.87E-18 |
| Ankrd1 | 8.615118 | 0.940802 | 5.33E-20 | 7.19E-18 |
| Bst1 | 2.351102 | 0.25717 | 6.12E-20 | 8.21E-18 |
| Galnt9 | -3.89753 | 0.427235 | 7.33E-20 | 9.78E-18 |
| Bcl3 | 2.520643 | 0.276348 | 7.43E-20 | 9.85E-18 |
| Gfpt2 | 2.508237 | 0.27513 | 7.75E-20 | 1.02E-17 |
| Fstl3 | 2.734901 | 0.300308 | 8.47E-20 | 1.11E-17 |
| Lig1 | 2.418701 | 0.265715 | 8.82E-20 | 1.15E-17 |
| Selp | 2.233288 | 0.246009 | 1.10E-19 | 1.43E-17 |
| Fam111a | 2.939025 | 0.324086 | 1.20E-19 | 1.55E-17 |
| Vwa5b2 | -3.66171 | 0.405205 | 1.61E-19 | 2.04E-17 |
| Mcm5 | 3.657124 | 0.405199 | 1.79E-19 | 2.24E-17 |
| Ch25h | 2.874641 | 0.319336 | 2.22E-19 | 2.75E-17 |
| Hlf | -3.8571 | 0.430692 | 3.38E-19 | 4.14E-17 |
| Serpina3g | 2.369695 | 0.264944 | 3.75E-19 | 4.55E-17 |
| Chl1 | 2.172043 | 0.242977 | 3.92E-19 | 4.73E-17 |
| Kif5a | -2.19868 | 0.245981 | 3.95E-19 | 4.74E-17 |
| Prss57 | 2.661519 | 0.298112 | 4.34E-19 | 5.18E-17 |
| Chrdl1 | -2.47443 | 0.277213 | 4.41E-19 | 5.24E-17 |
| Cebpd | 2.245062 | 0.25176 | 4.77E-19 | 5.61E-17 |
| Gm10143 | 2.853915 | 0.320572 | 5.46E-19 | 6.38E-17 |
| Pdpn | 2.092749 | 0.235874 | 7.16E-19 | 8.34E-17 |
| Hpse2 | -3.57751 | 0.403734 | 7.93E-19 | 9.18E-17 |
| Ripk3 | 3.053174 | 0.34494 | 8.65E-19 | 9.96E-17 |
| Plaur | 3.062788 | 0.346814 | 1.04E-18 | 1.18E-16 |
| Cmya5 | -3.47251 | 0.393866 | 1.18E-18 | 1.34E-16 |
| Tgm1 | 3.573732 | 0.40571 | 1.27E-18 | 1.43E-16 |
| Rgs2 | -2.5233 | 0.286564 | 1.30E-18 | 1.46E-16 |
| Rgs7bp | -2.53033 | 0.287671 | 1.42E-18 | 1.58E-16 |
| Adam19 | 2.346969 | 0.267031 | 1.51E-18 | 1.66E-16 |
| Fen1 | 2.810169 | 0.320096 | 1.65E-18 | 1.80E-16 |

| | | | | |
|---|---|---|---|---|
| Slc5a1 | -3.92244 | 0.447561 | 1.88E-18 | 2.02E-16 |
| Car11 | -2.44625 | 0.279739 | 2.23E-18 | 2.37E-16 |
| Rnd1 | 4.55058 | 0.520655 | 2.33E-18 | 2.46E-16 |
| Tmem200c | -3.39505 | 0.38874 | 2.47E-18 | 2.60E-16 |
| Slc6a2 | 2.674525 | 0.307777 | 3.63E-18 | 3.80E-16 |
| Atad2 | 2.181255 | 0.251395 | 4.08E-18 | 4.25E-16 |
| Itgam | 2.271391 | 0.261875 | 4.19E-18 | 4.34E-16 |
| Stk32c | -2.69981 | 0.311597 | 4.54E-18 | 4.68E-16 |
| Gmnn | 2.74632 | 0.318789 | 7.00E-18 | 7.20E-16 |
| Cldn10 | -2.14636 | 0.249423 | 7.61E-18 | 7.75E-16 |
| Lmnb1 | 2.452335 | 0.285083 | 7.82E-18 | 7.86E-16 |
| Mybl2 | 3.585629 | 0.416889 | 7.91E-18 | 7.91E-16 |
| 6330403A( | -2.17752 | 0.253264 | 8.12E-18 | 8.09E-16 |
| Pde5a | -2.17902 | 0.253494 | 8.26E-18 | 8.19E-16 |
| Bzrap1 | -3.11409 | 0.363317 | 1.02E-17 | 1.01E-15 |
| Gm11947 | -3.11147 | 0.363009 | 1.02E-17 | 1.01E-15 |
| Hmcn2 | -3.89323 | 0.455469 | 1.26E-17 | 1.22E-15 |
| Crebrf | -2.19576 | 0.256939 | 1.28E-17 | 1.23E-15 |
| Apold1 | 2.919955 | 0.341687 | 1.28E-17 | 1.23E-15 |
| Adam8 | 3.900327 | 0.456638 | 1.33E-17 | 1.27E-15 |
| Cited4 | -2.20027 | 0.257686 | 1.36E-17 | 1.30E-15 |
| Gkn3 | -4.22287 | 0.494876 | 1.42E-17 | 1.35E-15 |
| Ryr2 | -3.37502 | 0.395523 | 1.43E-17 | 1.35E-15 |
| Trdn | -3.69579 | 0.433524 | 1.53E-17 | 1.43E-15 |
| Mturn | -2.66146 | 0.312177 | 1.52E-17 | 1.43E-15 |
| Fibin | -2.71088 | 0.318041 | 1.55E-17 | 1.44E-15 |
| Clec4e | 5.692065 | 0.667926 | 1.57E-17 | 1.46E-15 |
| Marco | 6.386108 | 0.750108 | 1.69E-17 | 1.56E-15 |
| Mical2 | 2.007552 | 0.235844 | 1.71E-17 | 1.57E-15 |
| Esyt3 | -2.04287 | 0.240515 | 2.00E-17 | 1.83E-15 |
| Tnfsf18 | 6.5388 | 0.771091 | 2.25E-17 | 2.05E-15 |
| Tnfaip6 | 3.849932 | 0.454224 | 2.33E-17 | 2.11E-15 |
| Stil | 3.482941 | 0.411027 | 2.38E-17 | 2.14E-15 |
| Rmi2 | 2.792033 | 0.32971 | 2.49E-17 | 2.24E-15 |
| Cacna1b | -4.29785 | 0.508196 | 2.74E-17 | 2.45E-15 |
| Pla2g2d | -3.62143 | 0.429307 | 3.30E-17 | 2.93E-15 |
| 1700019D( | -2.60493 | 0.308961 | 3.42E-17 | 3.02E-15 |
| Mmp3 | 3.703475 | 0.44034 | 4.08E-17 | 3.55E-15 |
| Aunip | 4.318927 | 0.513618 | 4.14E-17 | 3.59E-15 |
| Cyp26b1 | -2.63832 | 0.31534 | 5.93E-17 | 5.10E-15 |
| Chaf1b | 3.259829 | 0.391019 | 7.64E-17 | 6.55E-15 |
| Syt12 | 3.445811 | 0.413729 | 8.18E-17 | 6.96E-15 |
| Brinp2 | -3.64616 | 0.440418 | 1.24E-16 | 1.04E-14 |
| Tuba1c | 2.205831 | 0.266448 | 1.25E-16 | 1.04E-14 |

| | | | | |
|---|---|---|---|---|
| Tlr13 | 2.354345 | 0.284505 | 1.28E-16 | 1.07E-14 |
| Gm9821 | -2.92199 | 0.353354 | 1.35E-16 | 1.11E-14 |
| Edn3 | -3.70024 | 0.447569 | 1.37E-16 | 1.13E-14 |
| Foxj1 | 2.981525 | 0.360783 | 1.41E-16 | 1.16E-14 |
| Ly6a | 2.234647 | 0.271413 | 1.82E-16 | 1.47E-14 |
| Mybl1 | 2.006603 | 0.244907 | 2.54E-16 | 2.04E-14 |
| Mall | 2.870741 | 0.350383 | 2.54E-16 | 2.04E-14 |
| Glp2r | -2.32161 | 0.283558 | 2.67E-16 | 2.13E-14 |
| Prss46 | 6.846882 | 0.837782 | 3.02E-16 | 2.39E-14 |
| Gm16033 | 2.272138 | 0.278222 | 3.17E-16 | 2.50E-14 |
| Ncapg2 | 2.890815 | 0.354128 | 3.26E-16 | 2.57E-14 |
| Nptx1 | -3.57062 | 0.43826 | 3.72E-16 | 2.92E-14 |
| Fhod3 | -2.54369 | 0.31227 | 3.77E-16 | 2.94E-14 |
| Cdc45 | 2.74229 | 0.337254 | 4.25E-16 | 3.27E-14 |
| Txlnb | -3.55915 | 0.43786 | 4.35E-16 | 3.34E-14 |
| Pik3ip1 | -2.33143 | 0.286881 | 4.41E-16 | 3.37E-14 |
| Atp2a3 | -2.01219 | 0.248059 | 4.99E-16 | 3.81E-14 |
| Hhipl1 | 2.328445 | 0.287188 | 5.16E-16 | 3.92E-14 |
| Prim1 | 2.269013 | 0.279917 | 5.23E-16 | 3.96E-14 |
| Grem1 | 5.254929 | 0.648506 | 5.36E-16 | 4.05E-14 |
| Dut | 2.378176 | 0.293642 | 5.55E-16 | 4.18E-14 |
| Rasd2 | -2.72263 | 0.336595 | 6.03E-16 | 4.49E-14 |
| Amotl2 | 2.0168 | 0.249763 | 6.76E-16 | 4.99E-14 |
| Pygm | -2.34731 | 0.290824 | 6.96E-16 | 5.12E-14 |
| Dhfr | 2.40296 | 0.299184 | 9.61E-16 | 6.96E-14 |
| Apitd1 | 2.550595 | 0.317614 | 9.71E-16 | 7.01E-14 |
| Nr1d1 | -2.13047 | 0.265803 | 1.10E-15 | 7.89E-14 |
| Gm17180 | -5.46951 | 0.682733 | 1.14E-15 | 8.12E-14 |
| Serpina3i | 4.565151 | 0.570254 | 1.19E-15 | 8.46E-14 |
| Klhl38 | -4.45756 | 0.558607 | 1.47E-15 | 1.03E-13 |
| Hsp25-ps1 | 2.355612 | 0.295577 | 1.59E-15 | 1.11E-13 |
| Cdh4 | -2.19992 | 0.2769 | 1.94E-15 | 1.35E-13 |
| B230312C | -3.8899 | 0.490011 | 2.05E-15 | 1.42E-13 |
| Dusp8 | 2.451073 | 0.308818 | 2.07E-15 | 1.43E-13 |
| Mcm7 | 2.155083 | 0.271726 | 2.17E-15 | 1.49E-13 |
| Cnnm1 | -3.21811 | 0.40599 | 2.25E-15 | 1.53E-13 |
| Pvr | 2.174471 | 0.274388 | 2.29E-15 | 1.55E-13 |
| Gpr176 | 3.017738 | 0.381884 | 2.74E-15 | 1.84E-13 |
| Nxpe5 | 2.160364 | 0.27395 | 3.12E-15 | 2.08E-13 |
| Igsf5 | -2.15848 | 0.273821 | 3.20E-15 | 2.12E-13 |
| Apobec2 | -3.72569 | 0.472957 | 3.34E-15 | 2.21E-13 |
| 9330159F1 | -2.64185 | 0.335968 | 3.74E-15 | 2.46E-13 |
| Il33 | 2.569261 | 0.327035 | 3.96E-15 | 2.59E-13 |
| AI480526 | -2.15947 | 0.275541 | 4.61E-15 | 2.96E-13 |

| | | | | |
|---|---|---|---|---|
| Lsamp | -3.5931 | 0.458958 | 4.92E-15 | 3.15E-13 |
| Angptl7 | 2.562998 | 0.327488 | 5.03E-15 | 3.20E-13 |
| Relt | 2.505298 | 0.32021 | 5.12E-15 | 3.25E-13 |
| Rgs6 | -2.32251 | 0.296948 | 5.23E-15 | 3.31E-13 |
| Il1b | 3.685733 | 0.471398 | 5.34E-15 | 3.37E-13 |
| A330023F2 | -3.28574 | 0.420706 | 5.72E-15 | 3.60E-13 |
| Chek1 | 3.262153 | 0.417721 | 5.75E-15 | 3.61E-13 |
| Gins1 | 2.530742 | 0.324397 | 6.12E-15 | 3.84E-13 |
| Fmo2 | -2.3697 | 0.303831 | 6.22E-15 | 3.88E-13 |
| Psat1 | 2.356242 | 0.302274 | 6.44E-15 | 3.99E-13 |
| Wfdc17 | 2.67187 | 0.343345 | 7.15E-15 | 4.41E-13 |
| Snhg11 | -3.0665 | 0.394093 | 7.19E-15 | 4.42E-13 |
| Kpna2 | 2.37007 | 0.304862 | 7.59E-15 | 4.64E-13 |
| Gm2788 | 4.59153 | 0.590749 | 7.70E-15 | 4.68E-13 |
| Gm15943 | -3.79963 | 0.489973 | 8.85E-15 | 5.37E-13 |
| Npl | 2.494313 | 0.321962 | 9.39E-15 | 5.67E-13 |
| Tbx5 | -2.02415 | 0.261622 | 1.02E-14 | 6.13E-13 |
| Dtx1 | -2.3984 | 0.310648 | 1.16E-14 | 6.93E-13 |
| Ncald | -2.18952 | 0.2843 | 1.35E-14 | 8.01E-13 |
| Plk4 | 2.390243 | 0.310698 | 1.44E-14 | 8.53E-13 |
| Fgl1 | 4.017258 | 0.522301 | 1.45E-14 | 8.62E-13 |
| Ppp1r18os | 2.084669 | 0.271113 | 1.48E-14 | 8.72E-13 |
| Rrm1 | 2.085805 | 0.271337 | 1.50E-14 | 8.84E-13 |
| Mlkl | 2.744283 | 0.357118 | 1.54E-14 | 9.00E-13 |
| Mcm4 | 2.324951 | 0.303709 | 1.93E-14 | 1.12E-12 |
| Kcnj12 | -3.37342 | 0.440768 | 1.96E-14 | 1.13E-12 |
| Ngb | -5.95206 | 0.779272 | 2.21E-14 | 1.26E-12 |
| Mansc4 | -3.81349 | 0.499617 | 2.30E-14 | 1.30E-12 |
| Vcan | 2.00366 | 0.262496 | 2.29E-14 | 1.30E-12 |
| Atcayos | -2.40108 | 0.314918 | 2.45E-14 | 1.38E-12 |
| Cdkl1 | -2.78204 | 0.36527 | 2.61E-14 | 1.47E-12 |
| Cldn2 | 4.218844 | 0.555765 | 3.17E-14 | 1.77E-12 |
| Ctla2a | 2.047136 | 0.26977 | 3.24E-14 | 1.81E-12 |
| Cdc6 | 2.754261 | 0.363096 | 3.31E-14 | 1.84E-12 |
| Galnt16 | -2.14572 | 0.282999 | 3.40E-14 | 1.89E-12 |
| Stc1 | -2.49716 | 0.329398 | 3.43E-14 | 1.90E-12 |
| Lilr4b | 2.31887 | 0.30614 | 3.60E-14 | 1.99E-12 |
| Gm42793 | 5.655802 | 0.748776 | 4.24E-14 | 2.31E-12 |
| Tet1 | -2.20277 | 0.292197 | 4.75E-14 | 2.57E-12 |
| Ankdd1a | -2.7171 | 0.361166 | 5.35E-14 | 2.88E-12 |
| Gm43278 | 3.495408 | 0.465151 | 5.71E-14 | 3.07E-12 |
| Sparcl1 | -2.21697 | 0.29525 | 5.97E-14 | 3.17E-12 |
| Ddah1 | 3.018429 | 0.402066 | 6.04E-14 | 3.19E-12 |
| Hnmt | -2.26702 | 0.30208 | 6.16E-14 | 3.25E-12 |

| | | | | |
|---|---|---|---|---|
| Gm12960 | 2.107939 | 0.28098 | 6.28E-14 | 3.31E-12 |
| A330093E2 | -3.99489 | 0.532717 | 6.43E-14 | 3.36E-12 |
| Cxcl1 | 4.273883 | 0.570501 | 6.81E-14 | 3.56E-12 |
| Ankrd63 | -5.26009 | 0.702468 | 6.99E-14 | 3.63E-12 |
| Actr3b | -2.4906 | 0.334169 | 9.12E-14 | 4.67E-12 |
| Cks1b | 2.415101 | 0.324268 | 9.49E-14 | 4.81E-12 |
| Gm19658 | 8.925547 | 1.200438 | 1.04E-13 | 5.26E-12 |
| Dna2 | 2.727749 | 0.367077 | 1.08E-13 | 5.40E-12 |
| Sirpb1c | 3.472728 | 0.468015 | 1.17E-13 | 5.84E-12 |
| A730049H( | 2.665167 | 0.359442 | 1.22E-13 | 6.07E-12 |
| 7420461P1 | -4.37616 | 0.590503 | 1.25E-13 | 6.23E-12 |
| Lif | 2.6339 | 0.355641 | 1.30E-13 | 6.42E-12 |
| Pole2 | 2.482245 | 0.335443 | 1.36E-13 | 6.71E-12 |
| Fam167b | 2.353334 | 0.318104 | 1.38E-13 | 6.79E-12 |
| Nr3c2 | -2.337 | 0.316189 | 1.46E-13 | 7.11E-12 |
| 4930427A( | 2.872114 | 0.388661 | 1.47E-13 | 7.17E-12 |
| Rapgef4 | -2.02479 | 0.274375 | 1.59E-13 | 7.66E-12 |
| Chst11 | 2.425361 | 0.328961 | 1.67E-13 | 8.02E-12 |
| Cenpm | 2.643728 | 0.358713 | 1.71E-13 | 8.17E-12 |
| Slfn9 | 2.105025 | 0.286 | 1.84E-13 | 8.76E-12 |
| Zwilch | 2.846771 | 0.386882 | 1.86E-13 | 8.87E-12 |
| Trim36 | -2.74556 | 0.373616 | 2.00E-13 | 9.52E-12 |
| Gfra3 | -2.67496 | 0.364144 | 2.04E-13 | 9.67E-12 |
| Itgam | 2.177621 | 0.29658 | 2.10E-13 | 9.90E-12 |
| Odc1 | 2.020683 | 0.275555 | 2.25E-13 | 1.05E-11 |
| 9330175E1 | -3.68779 | 0.503223 | 2.33E-13 | 1.09E-11 |
| Abca8a | -2.59949 | 0.354969 | 2.42E-13 | 1.12E-11 |
| Tfrc | 2.125483 | 0.290754 | 2.67E-13 | 1.23E-11 |
| 4631405K( | -2.02184 | 0.276911 | 2.85E-13 | 1.30E-11 |
| Trip13 | 3.339993 | 0.457475 | 2.86E-13 | 1.31E-11 |
| 4932422M | -2.01454 | 0.276002 | 2.90E-13 | 1.32E-11 |
| Dscc1 | 3.559745 | 0.488195 | 3.06E-13 | 1.39E-11 |
| Vsig4 | 2.965973 | 0.40779 | 3.51E-13 | 1.57E-11 |
| Hbegf | 2.429202 | 0.334007 | 3.52E-13 | 1.57E-11 |
| Kl | -3.93928 | 0.542203 | 3.72E-13 | 1.66E-11 |
| Asic2 | -3.61658 | 0.49984 | 4.64E-13 | 2.05E-11 |
| Pole | 3.159458 | 0.43723 | 4.97E-13 | 2.18E-11 |
| Tnfrsf23 | 2.456639 | 0.340066 | 5.05E-13 | 2.21E-11 |
| Cacna2d3 | -2.25286 | 0.311948 | 5.13E-13 | 2.24E-11 |
| Mms22l | 2.467171 | 0.341752 | 5.23E-13 | 2.29E-11 |
| Fkbp11 | 2.131343 | 0.295893 | 5.89E-13 | 2.55E-11 |
| Arhgef26 | -2.04951 | 0.284605 | 5.97E-13 | 2.58E-11 |
| Cpxm2 | -2.3925 | 0.33245 | 6.17E-13 | 2.66E-11 |
| Kcna2 | -2.78011 | 0.386877 | 6.67E-13 | 2.85E-11 |

| | | | | |
|---|---|---|---|---|
| Rab36 | -2.15941 | 0.300726 | 6.94E-13 | 2.96E-11 |
| Gm16548 | 2.36294 | 0.32911 | 6.98E-13 | 2.98E-11 |
| 1810041L1 | -3.07424 | 0.430042 | 8.76E-13 | 3.69E-11 |
| Acrbp | -2.08435 | 0.291615 | 8.83E-13 | 3.71E-11 |
| Nsg2 | -2.91914 | 0.408492 | 8.93E-13 | 3.74E-11 |
| Kcnk3 | -2.15947 | 0.302275 | 9.06E-13 | 3.79E-11 |
| Ager | -2.79332 | 0.391149 | 9.24E-13 | 3.85E-11 |
| Areg | 5.679055 | 0.795218 | 9.23E-13 | 3.85E-11 |
| Hmgb2 | 2.11067 | 0.295609 | 9.33E-13 | 3.88E-11 |
| Uchl1os | -2.60237 | 0.364552 | 9.43E-13 | 3.92E-11 |
| Ttr | -8.37372 | 1.181137 | 1.35E-12 | 5.48E-11 |
| Tubb6 | 2.03795 | 0.287564 | 1.37E-12 | 5.57E-11 |
| Tk1 | 3.005115 | 0.424138 | 1.39E-12 | 5.63E-11 |
| Frzb | 2.178534 | 0.307508 | 1.40E-12 | 5.65E-11 |
| B3gnt3 | 2.279458 | 0.322102 | 1.47E-12 | 5.95E-11 |
| Itga5 | 2.837212 | 0.401106 | 1.51E-12 | 6.09E-11 |
| Islr | -2.38261 | 0.336878 | 1.52E-12 | 6.12E-11 |
| Bdkrb2 | 2.124587 | 0.300742 | 1.61E-12 | 6.46E-11 |
| Inmt | -5.09845 | 0.721757 | 1.62E-12 | 6.48E-11 |
| Arid3c | -4.08833 | 0.578848 | 1.63E-12 | 6.50E-11 |
| E030013I1 | -3.54942 | 0.50393 | 1.87E-12 | 7.44E-11 |
| Fn3k | -4.25932 | 0.604925 | 1.91E-12 | 7.55E-11 |
| Dmrtc1a | -4.5441 | 0.645417 | 1.91E-12 | 7.57E-11 |
| Pclo | -2.23378 | 0.318005 | 2.15E-12 | 8.46E-11 |
| Eya1 | -2.36726 | 0.337351 | 2.26E-12 | 8.87E-11 |
| Tyms | 2.079251 | 0.296558 | 2.36E-12 | 9.21E-11 |
| Ngf | 2.491202 | 0.355692 | 2.49E-12 | 9.69E-11 |
| Kifc2 | -2.74486 | 0.391933 | 2.50E-12 | 9.71E-11 |
| Kcnmb4os | -3.06605 | 0.43787 | 2.52E-12 | 9.76E-11 |
| Rad18 | 2.300475 | 0.328533 | 2.52E-12 | 9.76E-11 |
| Gfra1 | -2.25385 | 0.322178 | 2.64E-12 | 1.02E-10 |
| Gm16175 | 2.701347 | 0.386574 | 2.79E-12 | 1.08E-10 |
| Steap1 | 3.076761 | 0.440474 | 2.85E-12 | 1.09E-10 |
| Gm16034 | 2.199856 | 0.315442 | 3.08E-12 | 1.18E-10 |
| Plat | 2.360541 | 0.338614 | 3.14E-12 | 1.20E-10 |
| Ms4a4a | 2.355067 | 0.339299 | 3.89E-12 | 1.45E-10 |
| Tgfbr3l | -2.51484 | 0.362933 | 4.23E-12 | 1.57E-10 |
| Nt5dc2 | 2.391449 | 0.345137 | 4.24E-12 | 1.57E-10 |
| Gm29491 | 2.915851 | 0.420895 | 4.28E-12 | 1.58E-10 |
| Gm26777 | -2.40979 | 0.347876 | 4.29E-12 | 1.58E-10 |
| Agtr2 | -4.22888 | 0.610998 | 4.48E-12 | 1.64E-10 |
| P4ha3 | 3.052321 | 0.441201 | 4.57E-12 | 1.68E-10 |
| Cys1 | -2.84166 | 0.411066 | 4.75E-12 | 1.74E-10 |
| Gm38102 | -2.15453 | 0.312 | 5.00E-12 | 1.82E-10 |

| | | | | |
|---|---|---|---|---|
| Pnck | -2.02579 | 0.293387 | 5.03E-12 | 1.82E-10 |
| Fbxl22 | -2.42379 | 0.351394 | 5.29E-12 | 1.90E-10 |
| Kcne3 | 3.358343 | 0.487051 | 5.38E-12 | 1.93E-10 |
| Fbn1 | 2.028268 | 0.294338 | 5.54E-12 | 1.98E-10 |
| Smc2 | 2.45135 | 0.356287 | 5.97E-12 | 2.13E-10 |
| Mapt | -2.16073 | 0.314366 | 6.27E-12 | 2.23E-10 |
| Bche | -2.78347 | 0.40615 | 7.22E-12 | 2.53E-10 |
| Ctsf | -2.1166 | 0.308964 | 7.35E-12 | 2.57E-10 |
| Dnajc6 | -2.40235 | 0.351855 | 8.63E-12 | 3.00E-10 |
| Gm44335 | 2.284547 | 0.334709 | 8.76E-12 | 3.03E-10 |
| Wfdc15b | -2.20651 | 0.323657 | 9.27E-12 | 3.20E-10 |
| Mmp8 | 7.504088 | 1.102875 | 1.02E-11 | 3.50E-10 |
| Kif4 | 2.250025 | 0.331055 | 1.07E-11 | 3.68E-10 |
| Gm16285 | -2.57232 | 0.378515 | 1.08E-11 | 3.69E-10 |
| Slc16a3 | 2.182074 | 0.32156 | 1.15E-11 | 3.94E-10 |
| Gm13415 | 5.097621 | 0.751344 | 1.16E-11 | 3.97E-10 |
| Baiap2l2 | -3.13668 | 0.462968 | 1.24E-11 | 4.22E-10 |
| Col20a1 | -2.05605 | 0.303806 | 1.31E-11 | 4.44E-10 |
| Pacsin1 | -2.89936 | 0.429 | 1.40E-11 | 4.71E-10 |
| Tmem200a | 2.395952 | 0.354825 | 1.45E-11 | 4.90E-10 |
| Gpr35 | 2.036974 | 0.301886 | 1.50E-11 | 5.05E-10 |
| Cdc7 | 2.357911 | 0.349645 | 1.54E-11 | 5.16E-10 |
| Cpm | -2.42429 | 0.360002 | 1.65E-11 | 5.48E-10 |
| Cda | 2.815665 | 0.419064 | 1.83E-11 | 6.04E-10 |
| AA467197 | 4.09185 | 0.610742 | 2.09E-11 | 6.81E-10 |
| Csgalnact1 | 2.307636 | 0.345113 | 2.28E-11 | 7.40E-10 |
| Tcea3 | -2.41087 | 0.360933 | 2.40E-11 | 7.74E-10 |
| Gm12840 | 3.511273 | 0.526428 | 2.56E-11 | 8.22E-10 |
| Ddias | 2.260809 | 0.339096 | 2.61E-11 | 8.35E-10 |
| Fbxw27 | -4.13714 | 0.620805 | 2.66E-11 | 8.50E-10 |
| Orc1 | 3.611614 | 0.542015 | 2.68E-11 | 8.54E-10 |
| Calcoco1 | -2.0873 | 0.313798 | 2.90E-11 | 9.22E-10 |
| Gata5 | -2.12608 | 0.320143 | 3.11E-11 | 9.84E-10 |
| Ncapd2 | 2.149262 | 0.32365 | 3.12E-11 | 9.85E-10 |
| Psd2 | -3.5688 | 0.537656 | 3.19E-11 | 9.98E-10 |
| Kcnmb4 | -3.08119 | 0.464923 | 3.42E-11 | 1.07E-09 |
| Tox2 | -2.23697 | 0.337552 | 3.43E-11 | 1.07E-09 |
| Efcc1 | -2.18555 | 0.330176 | 3.61E-11 | 1.12E-09 |
| Rapgef4os1 | -3.33595 | 0.504122 | 3.66E-11 | 1.13E-09 |
| Pln | -3.3143 | 0.501087 | 3.73E-11 | 1.16E-09 |
| Chtf18 | 2.81385 | 0.425818 | 3.89E-11 | 1.20E-09 |
| Gm44220 | -3.082 | 0.46688 | 4.08E-11 | 1.25E-09 |
| H2afx | 2.093472 | 0.317913 | 4.55E-11 | 1.39E-09 |
| Gm28729 | -4.99829 | 0.759082 | 4.56E-11 | 1.39E-09 |

| | | | | |
|---|---|---|---|---|
| Dnph1 | 2.114568 | 0.321145 | 4.57E-11 | 1.39E-09 |
| Gjb5 | 2.134139 | 0.324347 | 4.71E-11 | 1.43E-09 |
| Brip1 | 2.636949 | 0.400839 | 4.75E-11 | 1.44E-09 |
| Kcnmb4os | -3.27881 | 0.499124 | 5.06E-11 | 1.52E-09 |
| Stab2 | -2.31794 | 0.353095 | 5.22E-11 | 1.57E-09 |
| Adam33 | -2.33653 | 0.356125 | 5.35E-11 | 1.60E-09 |
| Gm6161 | 2.708328 | 0.412897 | 5.41E-11 | 1.62E-09 |
| Traip | 2.864785 | 0.437986 | 6.12E-11 | 1.81E-09 |
| Per3 | -2.16747 | 0.331427 | 6.16E-11 | 1.82E-09 |
| Gm16185 | -2.72139 | 0.416426 | 6.36E-11 | 1.87E-09 |
| Gm6397 | 3.540044 | 0.541695 | 6.36E-11 | 1.87E-09 |
| E130012A1 | 3.21347 | 0.492283 | 6.68E-11 | 1.96E-09 |
| Gprasp2 | -3.04738 | 0.467498 | 7.10E-11 | 2.08E-09 |
| Tubb5 | 2.009369 | 0.30911 | 8.01E-11 | 2.32E-09 |
| Incenp | 2.855568 | 0.439753 | 8.38E-11 | 2.42E-09 |
| Klf15 | -2.96564 | 0.457584 | 9.11E-11 | 2.61E-09 |
| Dbf4 | 2.617357 | 0.403929 | 9.19E-11 | 2.63E-09 |
| Srrm4 | 2.369153 | 0.36574 | 9.31E-11 | 2.66E-09 |
| Mad2l1 | 2.522044 | 0.389519 | 9.50E-11 | 2.71E-09 |
| Cxcl3 | 6.838259 | 1.056805 | 9.76E-11 | 2.78E-09 |
| 4933406C1 | -2.82255 | 0.436355 | 9.90E-11 | 2.82E-09 |
| Tfpi2 | 2.042069 | 0.316043 | 1.04E-10 | 2.94E-09 |
| Klhl30 | -2.25054 | 0.348685 | 1.09E-10 | 3.06E-09 |
| Pbld2 | -2.49811 | 0.388152 | 1.23E-10 | 3.43E-09 |
| Mir1906-1 | -2.22375 | 0.345618 | 1.24E-10 | 3.47E-09 |
| Prr36 | -2.00018 | 0.311274 | 1.31E-10 | 3.65E-09 |
| Ppil6 | -2.7802 | 0.432806 | 1.33E-10 | 3.69E-09 |
| Gm44876 | 2.434289 | 0.379037 | 1.34E-10 | 3.72E-09 |
| H2-K2 | -3.072 | 0.479563 | 1.50E-10 | 4.10E-09 |
| Bcl11a | -2.09313 | 0.326855 | 1.52E-10 | 4.15E-09 |
| Slc46a2 | -2.31456 | 0.361715 | 1.57E-10 | 4.28E-09 |
| Gm28221 | -2.30019 | 0.359609 | 1.59E-10 | 4.33E-09 |
| E2f7 | 3.26425 | 0.510569 | 1.62E-10 | 4.41E-09 |
| Bglap2 | -4.1086 | 0.64354 | 1.72E-10 | 4.66E-09 |
| Ncaph | 2.87157 | 0.450188 | 1.79E-10 | 4.82E-09 |
| E2f1 | 2.448131 | 0.383952 | 1.82E-10 | 4.89E-09 |
| Cenpn | 2.802919 | 0.440134 | 1.91E-10 | 5.13E-09 |
| Cxcl5 | 5.930596 | 0.931854 | 1.96E-10 | 5.25E-09 |
| Bmp8a | -2.27057 | 0.357148 | 2.05E-10 | 5.47E-09 |
| Lrg1 | 2.397782 | 0.377577 | 2.15E-10 | 5.70E-09 |
| 2310002F0 | -2.73592 | 0.431539 | 2.30E-10 | 6.08E-09 |
| St6gal2 | -4.20417 | 0.663374 | 2.33E-10 | 6.16E-09 |
| Dgki | -2.56907 | 0.405379 | 2.34E-10 | 6.16E-09 |
| Sgca | -2.3312 | 0.36881 | 2.60E-10 | 6.83E-09 |

| | | | | |
|---|---|---|---|---|
| Ifit3b | -2.40246 | 0.380753 | 2.79E-10 | 7.28E-09 |
| Akr1b8 | 2.377467 | 0.377208 | 2.92E-10 | 7.60E-09 |
| Hs6st2 | 3.1731 | 0.503813 | 3.01E-10 | 7.82E-09 |
| Lgi2 | 2.141922 | 0.340454 | 3.15E-10 | 8.14E-09 |
| Ppp1r3c | -2.4033 | 0.382039 | 3.16E-10 | 8.15E-09 |
| Ncapg | 2.781577 | 0.442176 | 3.16E-10 | 8.15E-09 |
| Skap1 | -2.97195 | 0.475774 | 4.20E-10 | 1.06E-08 |
| Gsg2 | 3.381889 | 0.541652 | 4.27E-10 | 1.08E-08 |
| Spp1 | 5.132535 | 0.822157 | 4.30E-10 | 1.09E-08 |
| Tcf19 | 2.41664 | 0.387816 | 4.62E-10 | 1.16E-08 |
| Lrr1 | 4.08477 | 0.656334 | 4.86E-10 | 1.21E-08 |
| Gm13461 | 2.149522 | 0.345629 | 5.00E-10 | 1.24E-08 |
| Slco5a1 | 2.706661 | 0.435323 | 5.05E-10 | 1.26E-08 |
| Cyp3a57 | -2.42107 | 0.389474 | 5.09E-10 | 1.27E-08 |
| Gpbar1 | 3.163279 | 0.509228 | 5.23E-10 | 1.30E-08 |
| Dpy19l2 | -3.114 | 0.502087 | 5.57E-10 | 1.38E-08 |
| Sgol1 | 3.003092 | 0.48471 | 5.80E-10 | 1.43E-08 |
| Nat8f3 | -3.4299 | 0.554441 | 6.16E-10 | 1.51E-08 |
| Gm42835 | 3.132777 | 0.506562 | 6.23E-10 | 1.53E-08 |
| Ska3 | 2.904655 | 0.470489 | 6.67E-10 | 1.63E-08 |
| Gm15851 | -2.04399 | 0.331231 | 6.79E-10 | 1.66E-08 |
| Fam132b | 3.01665 | 0.489736 | 7.29E-10 | 1.77E-08 |
| Polq | 2.938012 | 0.477218 | 7.44E-10 | 1.80E-08 |
| Hdac11 | -2.34443 | 0.381028 | 7.61E-10 | 1.84E-08 |
| Upb1 | -2.65567 | 0.43238 | 8.15E-10 | 1.96E-08 |
| Retnlg | 4.568778 | 0.744161 | 8.28E-10 | 1.98E-08 |
| Syce2 | 2.347194 | 0.382893 | 8.78E-10 | 2.08E-08 |
| Peg12 | 2.199026 | 0.358743 | 8.80E-10 | 2.09E-08 |
| Plch1 | 3.075186 | 0.501834 | 8.90E-10 | 2.10E-08 |
| Gm17552 | -2.10675 | 0.344244 | 9.36E-10 | 2.19E-08 |
| Olr1 | 5.133672 | 0.840037 | 9.89E-10 | 2.31E-08 |
| Arg2 | -2.31554 | 0.378984 | 9.97E-10 | 2.33E-08 |
| Gm9899 | -2.50035 | 0.40949 | 1.02E-09 | 2.38E-08 |
| Peli3 | -2.24586 | 0.368514 | 1.10E-09 | 2.55E-08 |
| Spdl1 | 3.119074 | 0.511907 | 1.11E-09 | 2.57E-08 |
| Cxcl2 | 8.484326 | 1.393257 | 1.13E-09 | 2.61E-08 |
| 3425401B1 | -2.97229 | 0.488465 | 1.17E-09 | 2.68E-08 |
| Gjb2 | 2.1454 | 0.353002 | 1.22E-09 | 2.80E-08 |
| Opcml | -2.89981 | 0.477777 | 1.28E-09 | 2.93E-08 |
| Ccl12 | 2.564017 | 0.422468 | 1.29E-09 | 2.93E-08 |
| Arhgap11a | 2.439993 | 0.402204 | 1.31E-09 | 2.97E-08 |
| E230032D2 | 2.204923 | 0.363623 | 1.33E-09 | 3.01E-08 |
| Gm26673 | -2.89531 | 0.47766 | 1.35E-09 | 3.05E-08 |
| Il17b | 3.93475 | 0.650069 | 1.42E-09 | 3.20E-08 |

| | | | | |
|---|---|---|---|---|
| Gata5os | -2.55924 | 0.423362 | 1.49E-09 | 3.34E-08 |
| Cd46 | -2.48268 | 0.411322 | 1.58E-09 | 3.51E-08 |
| Slc7a5 | 2.036216 | 0.337675 | 1.64E-09 | 3.63E-08 |
| Gm44002 | -2.57092 | 0.427371 | 1.79E-09 | 3.96E-08 |
| Cd300lf | 3.957247 | 0.657984 | 1.81E-09 | 3.99E-08 |
| Gm14025 | 4.456143 | 0.744505 | 2.16E-09 | 4.67E-08 |
| Lin7b | -2.89041 | 0.483383 | 2.24E-09 | 4.82E-08 |
| Vtcn1 | 2.250393 | 0.376671 | 2.31E-09 | 4.97E-08 |
| Fam214a | -2.06432 | 0.346082 | 2.45E-09 | 5.26E-08 |
| Cenpw | 2.182556 | 0.366042 | 2.48E-09 | 5.32E-08 |
| Cdh3 | 2.125389 | 0.356643 | 2.53E-09 | 5.41E-08 |
| Gm16432 | -2.21835 | 0.372517 | 2.60E-09 | 5.55E-08 |
| Sntg2 | -2.22752 | 0.375014 | 2.85E-09 | 6.05E-08 |
| Alpk2 | 3.189574 | 0.537039 | 2.86E-09 | 6.07E-08 |
| Gm15677 | -2.62668 | 0.442566 | 2.94E-09 | 6.21E-08 |
| Ntf3 | -2.60698 | 0.439489 | 3.00E-09 | 6.32E-08 |
| Lctl | 2.54657 | 0.430346 | 3.27E-09 | 6.80E-08 |
| Cxcr2 | 3.59619 | 0.60796 | 3.32E-09 | 6.87E-08 |
| Brca1 | 2.716192 | 0.459661 | 3.44E-09 | 7.11E-08 |
| Gm4956 | -5.23186 | 0.886908 | 3.66E-09 | 7.54E-08 |
| C430002N | 7.603223 | 1.2899 | 3.76E-09 | 7.73E-08 |
| Ms4a8a | 2.669913 | 0.453369 | 3.88E-09 | 7.97E-08 |
| Gm16701 | -2.35717 | 0.400428 | 3.94E-09 | 8.06E-08 |
| Serpinb9b | 2.750706 | 0.467896 | 4.13E-09 | 8.40E-08 |
| Rims1 | -2.3925 | 0.407277 | 4.24E-09 | 8.61E-08 |
| Fut2 | 2.155646 | 0.367326 | 4.40E-09 | 8.89E-08 |
| Slc26a7 | 2.689576 | 0.458611 | 4.50E-09 | 9.08E-08 |
| Pigz | -3.06705 | 0.523589 | 4.69E-09 | 9.42E-08 |
| Hdc | 2.999085 | 0.512038 | 4.71E-09 | 9.44E-08 |
| 04-Sep | -2.01886 | 0.345114 | 4.92E-09 | 9.83E-08 |
| Nsl1 | 2.775463 | 0.475436 | 5.29E-09 | 1.05E-07 |
| Kcnh2 | -2.07759 | 0.357023 | 5.91E-09 | 1.17E-07 |
| Kifc5b | 2.405118 | 0.413373 | 5.95E-09 | 1.18E-07 |
| Zfp811 | -2.27164 | 0.390515 | 5.99E-09 | 1.18E-07 |
| Gm15792 | 3.020264 | 0.519636 | 6.16E-09 | 1.21E-07 |
| Mastl | 2.52037 | 0.433839 | 6.27E-09 | 1.23E-07 |
| Gm28513 | 2.718072 | 0.467954 | 6.31E-09 | 1.24E-07 |
| 8430408G: | -2.41858 | 0.416914 | 6.59E-09 | 1.29E-07 |
| Fbxo5 | 2.098611 | 0.361782 | 6.60E-09 | 1.29E-07 |
| Mgl2 | -2.50826 | 0.432464 | 6.63E-09 | 1.29E-07 |
| Fcgr4 | 2.45659 | 0.424064 | 6.92E-09 | 1.35E-07 |
| Dzank1 | -2.00947 | 0.346947 | 6.96E-09 | 1.35E-07 |
| Mir675 | 3.400004 | 0.588946 | 7.79E-09 | 1.50E-07 |
| D230017M | -2.29258 | 0.397202 | 7.84E-09 | 1.51E-07 |

| | | | | |
|---|---|---|---|---|
| Hist1h3c | 5.156636 | 0.893523 | 7.87E-09 | 1.51E-07 |
| Otogl | -2.56618 | 0.444885 | 8.01E-09 | 1.53E-07 |
| 4930447N( | -2.40144 | 0.416376 | 8.05E-09 | 1.54E-07 |
| Kif20b | 2.573915 | 0.44628 | 8.05E-09 | 1.54E-07 |
| RP23-462D | -2.12496 | 0.36886 | 8.37E-09 | 1.59E-07 |
| Gen1 | 2.19954 | 0.38217 | 8.64E-09 | 1.64E-07 |
| Mir147 | 4.447163 | 0.773645 | 9.01E-09 | 1.71E-07 |
| Kcnc1 | -5.69722 | 0.991477 | 9.13E-09 | 1.73E-07 |
| Gm15179 | 2.693555 | 0.468795 | 9.15E-09 | 1.73E-07 |
| Tmem25 | -2.19238 | 0.381604 | 9.18E-09 | 1.74E-07 |
| Cldn8 | -2.25849 | 0.393278 | 9.32E-09 | 1.76E-07 |
| Gm12405 | -2.20674 | 0.385097 | 1.00E-08 | 1.89E-07 |
| Rnf183 | 3.375624 | 0.589534 | 1.03E-08 | 1.93E-07 |
| Gm27786 | 3.276372 | 0.572512 | 1.05E-08 | 1.96E-07 |
| Ttc21a | -2.5203 | 0.440422 | 1.05E-08 | 1.96E-07 |
| Fbn2 | 4.501426 | 0.787286 | 1.08E-08 | 2.02E-07 |
| Tmem61 | -6.40815 | 1.123504 | 1.17E-08 | 2.17E-07 |
| Dnah6 | 3.125191 | 0.549389 | 1.28E-08 | 2.36E-07 |
| Edar | -3.42758 | 0.603045 | 1.32E-08 | 2.41E-07 |
| Mybpc1 | -3.15239 | 0.555547 | 1.39E-08 | 2.54E-07 |
| Klhdc8a | 2.025931 | 0.357039 | 1.39E-08 | 2.54E-07 |
| Rgs22 | -3.30917 | 0.583838 | 1.45E-08 | 2.62E-07 |
| F10 | 3.930239 | 0.693416 | 1.45E-08 | 2.62E-07 |
| Gm5069 | -2.39894 | 0.424106 | 1.55E-08 | 2.80E-07 |
| Dhtkd1 | -2.00552 | 0.355561 | 1.70E-08 | 3.04E-07 |
| Dcdc2a | -6.35015 | 1.126322 | 1.72E-08 | 3.08E-07 |
| Mtfr2 | 3.51443 | 0.623852 | 1.77E-08 | 3.15E-07 |
| Gm14573 | -2.27493 | 0.403994 | 1.79E-08 | 3.18E-07 |
| Sell | 2.683763 | 0.476804 | 1.82E-08 | 3.22E-07 |
| Angpt4 | 2.769306 | 0.492064 | 1.82E-08 | 3.23E-07 |
| Cenpk | 2.87001 | 0.510348 | 1.87E-08 | 3.30E-07 |
| Tgm3 | -2.17593 | 0.387024 | 1.89E-08 | 3.32E-07 |
| Kif24 | 2.049969 | 0.365611 | 2.06E-08 | 3.61E-07 |
| Esr2 | -4.5099 | 0.804683 | 2.09E-08 | 3.65E-07 |
| Aox3 | -2.42586 | 0.43285 | 2.09E-08 | 3.65E-07 |
| Acer2 | -2.07132 | 0.370238 | 2.21E-08 | 3.85E-07 |
| Ypel4 | -2.35235 | 0.420504 | 2.22E-08 | 3.85E-07 |
| Epgn | 3.88826 | 0.695446 | 2.26E-08 | 3.91E-07 |
| Carmn | -2.0154 | 0.360565 | 2.28E-08 | 3.94E-07 |
| Myoz2 | 4.275542 | 0.765293 | 2.31E-08 | 3.99E-07 |
| Nupr1l | -2.12545 | 0.380869 | 2.40E-08 | 4.11E-07 |
| 4732414G( | -4.23955 | 0.761068 | 2.54E-08 | 4.33E-07 |
| Maob | -2.13328 | 0.383158 | 2.58E-08 | 4.39E-07 |
| Syt1 | -2.65784 | 0.477405 | 2.59E-08 | 4.39E-07 |

| | | | | |
|---|---|---|---|---|
| Kazald1 | -2.08965 | 0.375415 | 2.60E-08 | 4.41E-07 |
| Abca8b | -2.216 | 0.398334 | 2.65E-08 | 4.49E-07 |
| Espn | -2.00889 | 0.3613 | 2.70E-08 | 4.55E-07 |
| Trim59 | 2.406024 | 0.432877 | 2.73E-08 | 4.60E-07 |
| A730056A( | -2.84049 | 0.512512 | 2.99E-08 | 5.01E-07 |
| Gm23187 | 2.474997 | 0.446663 | 3.01E-08 | 5.04E-07 |
| Nrm | 2.042486 | 0.368992 | 3.11E-08 | 5.19E-07 |
| Hotair | -2.48604 | 0.449197 | 3.12E-08 | 5.21E-07 |
| Cytl1 | -2.67609 | 0.483682 | 3.15E-08 | 5.26E-07 |
| Gm12406 | -2.37969 | 0.430526 | 3.25E-08 | 5.40E-07 |
| BC051019 | -2.32604 | 0.420852 | 3.26E-08 | 5.40E-07 |
| Kcnk12 | 6.563894 | 1.189309 | 3.41E-08 | 5.63E-07 |
| Dnajc28 | -2.09041 | 0.378895 | 3.45E-08 | 5.68E-07 |
| Ulbp1 | 2.761125 | 0.501523 | 3.68E-08 | 6.01E-07 |
| Rgs9bp | -2.9278 | 0.532109 | 3.75E-08 | 6.12E-07 |
| Ecm2 | -2.28861 | 0.416277 | 3.85E-08 | 6.24E-07 |
| Osm | 3.239978 | 0.589816 | 3.95E-08 | 6.39E-07 |
| Saa3 | 6.840413 | 1.246152 | 4.04E-08 | 6.53E-07 |
| Gm13394 | 2.054697 | 0.374938 | 4.25E-08 | 6.85E-07 |
| Gng13 | 2.843363 | 0.520616 | 4.72E-08 | 7.52E-07 |
| Gm11816 | -3.55871 | 0.652759 | 4.99E-08 | 7.90E-07 |
| Nppb | 7.022095 | 1.288663 | 5.06E-08 | 8.01E-07 |
| Astn1 | -2.48428 | 0.456277 | 5.19E-08 | 8.18E-07 |
| Cks2 | 2.138838 | 0.393549 | 5.49E-08 | 8.58E-07 |
| Dynlt1a | 2.019113 | 0.371767 | 5.60E-08 | 8.75E-07 |
| Slc8a3 | -2.52458 | 0.465691 | 5.92E-08 | 9.22E-07 |
| Kbtbd6 | 3.146512 | 0.581525 | 6.27E-08 | 9.73E-07 |
| Gm16587 | 2.056753 | 0.380219 | 6.32E-08 | 9.79E-07 |
| Abcc2 | -5.50314 | 1.018433 | 6.53E-08 | 1.01E-06 |
| Uhrf1 | 4.036864 | 0.74828 | 6.86E-08 | 1.05E-06 |
| Gpr20 | -2.30024 | 0.426549 | 6.94E-08 | 1.06E-06 |
| Fpr2 | 3.771606 | 0.700032 | 7.13E-08 | 1.09E-06 |
| A330074K2 | -2.29785 | 0.426707 | 7.24E-08 | 1.10E-06 |
| Kcna6 | -2.00798 | 0.37306 | 7.35E-08 | 1.12E-06 |
| AI662270 | 2.02341 | 0.376003 | 7.39E-08 | 1.12E-06 |
| Dpyd | -2.09269 | 0.389217 | 7.59E-08 | 1.15E-06 |
| Evx1os | -3.7464 | 0.697535 | 7.83E-08 | 1.18E-06 |
| Pbp2 | 4.548449 | 0.847427 | 7.99E-08 | 1.20E-06 |
| Gm29371 | 5.600534 | 1.043461 | 7.99E-08 | 1.20E-06 |
| Slco4a1 | 2.069978 | 0.385694 | 8.01E-08 | 1.20E-06 |
| Bex4 | -2.72378 | 0.507592 | 8.05E-08 | 1.21E-06 |
| Upk2 | -2.01555 | 0.375622 | 8.06E-08 | 1.21E-06 |
| Bard1 | 2.865423 | 0.534254 | 8.17E-08 | 1.22E-06 |
| Gm27483 | 3.401389 | 0.634859 | 8.43E-08 | 1.26E-06 |

| | | | | |
|---|---|---|---|---|
| Kif23 | 2.260176 | 0.422638 | 8.90E-08 | 1.32E-06 |
| Gm973 | -2.39256 | 0.447421 | 8.92E-08 | 1.32E-06 |
| Nell2 | -3.02254 | 0.566923 | 9.74E-08 | 1.43E-06 |
| Mir5134 | 2.681538 | 0.503 | 9.76E-08 | 1.43E-06 |
| BC025446 | 2.974133 | 0.55864 | 1.02E-07 | 1.48E-06 |
| Car12 | 2.294984 | 0.431575 | 1.05E-07 | 1.52E-06 |
| Gm44985 | 2.016518 | 0.38072 | 1.18E-07 | 1.68E-06 |
| Kif18a | 2.132666 | 0.402974 | 1.21E-07 | 1.72E-06 |
| 5330417C2 | -2.61366 | 0.494051 | 1.22E-07 | 1.73E-06 |
| Cntnap3 | -3.37974 | 0.638964 | 1.23E-07 | 1.74E-06 |
| 4930426D0 | -2.50665 | 0.475285 | 1.33E-07 | 1.88E-06 |
| Hp | 4.082102 | 0.774139 | 1.34E-07 | 1.89E-06 |
| Fpr1 | 4.407164 | 0.837841 | 1.44E-07 | 2.02E-06 |
| C130021I2 | -3.4518 | 0.656454 | 1.45E-07 | 2.03E-06 |
| Foxo6 | -2.42589 | 0.461498 | 1.47E-07 | 2.05E-06 |
| Gm44805 | 3.518711 | 0.670432 | 1.53E-07 | 2.14E-06 |
| Apoc2 | 3.497621 | 0.666954 | 1.57E-07 | 2.18E-06 |
| Kcnt1 | -5.27407 | 1.005818 | 1.58E-07 | 2.18E-06 |
| Gm13807 | -2.86705 | 0.546905 | 1.59E-07 | 2.20E-06 |
| Matn4 | -2.46199 | 0.469668 | 1.59E-07 | 2.20E-06 |
| E2f8 | 4.671557 | 0.891437 | 1.60E-07 | 2.21E-06 |
| Cd177 | 5.713994 | 1.090592 | 1.61E-07 | 2.22E-06 |
| E130102H2 | -3.09424 | 0.591079 | 1.65E-07 | 2.27E-06 |
| Kcnip1 | -3.17111 | 0.606581 | 1.71E-07 | 2.35E-06 |
| Gm26902 | 2.197687 | 0.421536 | 1.85E-07 | 2.51E-06 |
| Dhrs7c | -7.339 | 1.411861 | 2.01E-07 | 2.71E-06 |
| Gm19461 | -3.54052 | 0.682323 | 2.12E-07 | 2.83E-06 |
| Tbx21 | -3.30773 | 0.637841 | 2.15E-07 | 2.88E-06 |
| Pcp4 | -4.17245 | 0.80523 | 2.20E-07 | 2.94E-06 |
| Lrrc7 | -2.02454 | 0.390716 | 2.20E-07 | 2.94E-06 |
| Dlgap2 | -2.98494 | 0.576204 | 2.21E-07 | 2.96E-06 |
| Slc6a18 | -3.38415 | 0.65527 | 2.41E-07 | 3.18E-06 |
| Tmprss11g | 2.410517 | 0.467009 | 2.45E-07 | 3.23E-06 |
| Sdr16c6 | -3.95366 | 0.76626 | 2.47E-07 | 3.26E-06 |
| Coch | -3.5842 | 0.695502 | 2.56E-07 | 3.35E-06 |
| Gm15452 | 2.230718 | 0.434367 | 2.81E-07 | 3.65E-06 |
| Mobp | -2.32067 | 0.453327 | 3.07E-07 | 3.96E-06 |
| Gm14005 | 2.20062 | 0.430398 | 3.17E-07 | 4.08E-06 |
| Reg3g | 5.308775 | 1.038408 | 3.18E-07 | 4.09E-06 |
| 2810408I1 | 3.060088 | 0.59872 | 3.20E-07 | 4.11E-06 |
| Angptl4 | 2.176871 | 0.426178 | 3.26E-07 | 4.17E-06 |
| Dnaic1 | -2.25491 | 0.441541 | 3.27E-07 | 4.19E-06 |
| C530008M | -2.22363 | 0.435634 | 3.32E-07 | 4.24E-06 |
| Ccdc113 | -2.46066 | 0.482197 | 3.34E-07 | 4.27E-06 |

| | | | | |
|---|---|---|---|---|
| Gm15917 | -2.78169 | 0.54555 | 3.42E-07 | 4.35E-06 |
| Col8a2 | -2.1965 | 0.430891 | 3.44E-07 | 4.38E-06 |
| St8sia2 | -2.97322 | 0.584691 | 3.67E-07 | 4.63E-06 |
| 9530026P( | -2.48941 | 0.491254 | 4.03E-07 | 5.03E-06 |
| Cthrc1 | 4.428215 | 0.873876 | 4.03E-07 | 5.03E-06 |
| E2f2 | 2.112776 | 0.417307 | 4.13E-07 | 5.14E-06 |
| Kcna1 | -2.17439 | 0.431356 | 4.64E-07 | 5.70E-06 |
| Tarm1 | 5.458642 | 1.082964 | 4.64E-07 | 5.71E-06 |
| Pla2g4e | -4.70157 | 0.933931 | 4.80E-07 | 5.88E-06 |
| Slc16a8 | -2.47597 | 0.491827 | 4.80E-07 | 5.88E-06 |
| Grip2 | -2.86334 | 0.569896 | 5.05E-07 | 6.17E-06 |
| Cdh8 | -3.4572 | 0.688202 | 5.07E-07 | 6.19E-06 |
| Lmo3 | -4.65536 | 0.927957 | 5.25E-07 | 6.40E-06 |
| Klhl24 | -2.10479 | 0.420186 | 5.47E-07 | 6.63E-06 |
| Crlf1 | 4.860664 | 0.972098 | 5.73E-07 | 6.91E-06 |
| Egr1 | 2.010157 | 0.402135 | 5.77E-07 | 6.96E-06 |
| Trp53inp1 | -2.01128 | 0.403048 | 6.03E-07 | 7.25E-06 |
| Gpr55 | 3.306842 | 0.662822 | 6.07E-07 | 7.28E-06 |
| Nrg1 | 2.316223 | 0.464968 | 6.31E-07 | 7.54E-06 |
| Mcmdc2 | 2.115416 | 0.425466 | 6.63E-07 | 7.88E-06 |
| Dyrk3 | 2.275592 | 0.457685 | 6.63E-07 | 7.88E-06 |
| Bmf | -2.05573 | 0.413932 | 6.82E-07 | 8.08E-06 |
| B230112G | -3.2032 | 0.645135 | 6.86E-07 | 8.12E-06 |
| Tnxb | -2.0697 | 0.417669 | 7.22E-07 | 8.48E-06 |
| Plb1 | -2.01757 | 0.407193 | 7.24E-07 | 8.49E-06 |
| Gjb4 | 3.576279 | 0.721994 | 7.30E-07 | 8.56E-06 |
| Capn11 | -2.40712 | 0.485992 | 7.31E-07 | 8.57E-06 |
| Glt8d2 | -2.00625 | 0.405236 | 7.39E-07 | 8.65E-06 |
| Rrm2 | 4.026934 | 0.813755 | 7.48E-07 | 8.74E-06 |
| Nmur2 | -2.99154 | 0.606195 | 8.02E-07 | 9.31E-06 |
| Kng2 | 3.357631 | 0.680686 | 8.11E-07 | 9.40E-06 |
| Htr1b | 2.473502 | 0.502322 | 8.47E-07 | 9.77E-06 |
| Hist1h4i | 2.000711 | 0.406567 | 8.61E-07 | 9.91E-06 |
| Vgf | 3.333211 | 0.679486 | 9.32E-07 | 1.07E-05 |
| Exo1 | 4.332185 | 0.883712 | 9.47E-07 | 1.08E-05 |
| Ripply1 | 7.39588 | 1.510893 | 9.83E-07 | 1.12E-05 |
| 2410004I0 | -4.08332 | 0.834911 | 1.00E-06 | 1.14E-05 |
| Cr2 | -5.67821 | 1.163593 | 1.06E-06 | 1.20E-05 |
| Tead4 | 2.467164 | 0.505629 | 1.06E-06 | 1.20E-05 |
| Unc13c | -2.7134 | 0.556465 | 1.08E-06 | 1.22E-05 |
| Clic6 | 2.302061 | 0.472827 | 1.12E-06 | 1.26E-05 |
| Timp1 | 4.813671 | 0.988998 | 1.13E-06 | 1.27E-05 |
| Ube2t | 2.069248 | 0.425835 | 1.18E-06 | 1.32E-05 |
| Slc16a12 | -3.36113 | 0.693493 | 1.26E-06 | 1.39E-05 |

| | | | | |
|---|---|---|---|---|
| Fbln7 | -2.33821 | 0.483247 | 1.31E-06 | 1.45E-05 |
| Lonrf2 | -2.19741 | 0.455223 | 1.39E-06 | 1.52E-05 |
| Abcc6 | -4.02729 | 0.834962 | 1.41E-06 | 1.55E-05 |
| Gm27219 | 2.436894 | 0.505977 | 1.46E-06 | 1.60E-05 |
| Shisa7 | -2.75173 | 0.574429 | 1.66E-06 | 1.79E-05 |
| 1700023F( | -3.03659 | 0.635328 | 1.76E-06 | 1.89E-05 |
| Gm44275 | 4.362533 | 0.914864 | 1.86E-06 | 1.98E-05 |
| 6330403K( | -2.24695 | 0.47202 | 1.93E-06 | 2.05E-05 |
| Dgkb | -2.33262 | 0.49052 | 1.98E-06 | 2.10E-05 |
| Mecomos | -2.00343 | 0.421443 | 2.00E-06 | 2.11E-05 |
| G530011O | 2.118231 | 0.44584 | 2.02E-06 | 2.14E-05 |
| Gdf5 | 2.115744 | 0.445967 | 2.09E-06 | 2.20E-05 |
| BC100530 | 7.027052 | 1.482546 | 2.14E-06 | 2.25E-05 |
| Ltb4r1 | 2.408708 | 0.508982 | 2.22E-06 | 2.32E-05 |
| 1700009N' | 5.123055 | 1.086278 | 2.40E-06 | 2.50E-05 |
| Galnt5 | -2.16416 | 0.459388 | 2.47E-06 | 2.55E-05 |
| Ltb4r2 | 4.779648 | 1.014596 | 2.47E-06 | 2.55E-05 |
| Ercc6l | 3.529186 | 0.749238 | 2.47E-06 | 2.56E-05 |
| Cadm2 | -2.06481 | 0.438955 | 2.55E-06 | 2.63E-05 |
| Gm12743 | 2.75438 | 0.586255 | 2.62E-06 | 2.70E-05 |
| Slc6a15 | -2.44515 | 0.520476 | 2.63E-06 | 2.70E-05 |
| Fignl1 | 4.039478 | 0.860483 | 2.67E-06 | 2.74E-05 |
| Gm11992 | -2.8289 | 0.603113 | 2.73E-06 | 2.79E-05 |
| 8430419K( | -2.13252 | 0.454702 | 2.73E-06 | 2.79E-05 |
| 3300005D( | 2.941476 | 0.627487 | 2.76E-06 | 2.82E-05 |
| RP23-218K | -4.53262 | 0.967084 | 2.77E-06 | 2.83E-05 |
| Grin2c | -3.89207 | 0.83328 | 3.00E-06 | 3.03E-05 |
| Foxo6os | -6.45418 | 1.382707 | 3.04E-06 | 3.07E-05 |
| Cyb5r2 | -2.53362 | 0.543874 | 3.19E-06 | 3.20E-05 |
| A930018M | -3.69333 | 0.793848 | 3.28E-06 | 3.29E-05 |
| Omd | -3.12867 | 0.672654 | 3.30E-06 | 3.30E-05 |
| Gm45036 | -2.11952 | 0.455754 | 3.31E-06 | 3.31E-05 |
| Mir6931 | -5.54897 | 1.19368 | 3.34E-06 | 3.34E-05 |
| Zfr2 | -2.01689 | 0.435222 | 3.58E-06 | 3.54E-05 |
| Htr2a | 2.070332 | 0.447732 | 3.76E-06 | 3.70E-05 |
| H2-M5 | -3.29449 | 0.71325 | 3.86E-06 | 3.77E-05 |
| Ces1g | -3.33989 | 0.723626 | 3.92E-06 | 3.82E-05 |
| Gm38165 | 4.202506 | 0.910654 | 3.93E-06 | 3.83E-05 |
| Gm20680 | -2.0731 | 0.449481 | 3.98E-06 | 3.87E-05 |
| Mir29c | -3.74108 | 0.811257 | 4.00E-06 | 3.88E-05 |
| Mthfd2 | 3.24406 | 0.7035 | 4.00E-06 | 3.89E-05 |
| Gm436 | -2.75296 | 0.597539 | 4.08E-06 | 3.95E-05 |
| Dscaml1 | 4.670914 | 1.015833 | 4.26E-06 | 4.11E-05 |
| Pmch | 3.119474 | 0.678791 | 4.31E-06 | 4.16E-05 |

| | | | | |
|---|---|---|---|---|
| Ppp1r1b | 3.471783 | 0.755946 | 4.38E-06 | 4.20E-05 |
| Sox11 | 4.708489 | 1.025963 | 4.45E-06 | 4.26E-05 |
| Mcm10 | 3.98634 | 0.869278 | 4.52E-06 | 4.32E-05 |
| Akap5 | 2.13645 | 0.467577 | 4.90E-06 | 4.65E-05 |
| Pdzrn4 | -3.34345 | 0.731788 | 4.90E-06 | 4.65E-05 |
| 4833427F1 | 2.619628 | 0.573639 | 4.96E-06 | 4.70E-05 |
| Prima1 | -2.57987 | 0.565465 | 5.06E-06 | 4.77E-05 |
| Krt14 | 5.530677 | 1.213433 | 5.17E-06 | 4.86E-05 |
| Ltbp2 | 3.247394 | 0.716556 | 5.84E-06 | 5.43E-05 |
| Gm16141 | -3.07347 | 0.678389 | 5.88E-06 | 5.46E-05 |
| Il1f9 | 4.01776 | 0.887055 | 5.92E-06 | 5.48E-05 |
| Fgf13 | -2.26526 | 0.500558 | 6.03E-06 | 5.56E-05 |
| Treml2 | 2.789203 | 0.617232 | 6.22E-06 | 5.71E-05 |
| Scn4b | -2.20385 | 0.487906 | 6.27E-06 | 5.76E-05 |
| Mir7688 | 4.480164 | 0.991879 | 6.28E-06 | 5.76E-05 |
| Dlx3 | -4.00869 | 0.887755 | 6.32E-06 | 5.79E-05 |
| RP24-82M4 | -2.86048 | 0.635022 | 6.65E-06 | 6.06E-05 |
| Gm5150 | 3.862717 | 0.857829 | 6.70E-06 | 6.10E-05 |
| Sh2d5 | 4.20366 | 0.9344 | 6.83E-06 | 6.21E-05 |
| Clec2f | 2.958427 | 0.660556 | 7.51E-06 | 6.72E-05 |
| Klhl33 | -3.0637 | 0.685155 | 7.77E-06 | 6.92E-05 |
| Gm21451 | 2.117594 | 0.474231 | 7.99E-06 | 7.10E-05 |
| Efnb3 | -2.55478 | 0.572467 | 8.09E-06 | 7.18E-05 |
| Gm44486 | 3.987304 | 0.894783 | 8.34E-06 | 7.37E-05 |
| Hist1h2an | 5.353937 | 1.201789 | 8.39E-06 | 7.41E-05 |
| Gm38037 | 2.939275 | 0.660641 | 8.62E-06 | 7.58E-05 |
| Cpne6 | -3.41269 | 0.767812 | 8.80E-06 | 7.73E-05 |
| Frmpd4 | -2.52368 | 0.570074 | 9.56E-06 | 8.30E-05 |
| Amn | -2.5792 | 0.582878 | 9.65E-06 | 8.37E-05 |
| Ticrr | 3.850174 | 0.870676 | 9.78E-06 | 8.46E-05 |
| Gm9115 | 2.699208 | 0.611465 | 1.01E-05 | 8.74E-05 |
| Gm8066 | -2.02742 | 0.459567 | 1.03E-05 | 8.83E-05 |
| Irf4 | -2.22317 | 0.505089 | 1.07E-05 | 9.21E-05 |
| Dtl | 4.266956 | 0.969903 | 1.09E-05 | 9.28E-05 |
| Klhl40 | 3.375222 | 0.76782 | 1.10E-05 | 9.41E-05 |
| Gm14760 | 2.331051 | 0.531887 | 1.17E-05 | 9.94E-05 |
| 1700095B1 | 2.066502 | 0.471672 | 1.18E-05 | 9.99E-05 |
| Sspo | -2.20127 | 0.502846 | 1.20E-05 | 0.000101 |
| Akp-ps1 | -3.37431 | 0.770993 | 1.21E-05 | 0.000102 |
| Gm17322 | -4.33242 | 0.989937 | 1.21E-05 | 0.000102 |
| Gm7336 | 2.378619 | 0.54359 | 1.21E-05 | 0.000102 |
| Gm2568 | -2.35331 | 0.53797 | 1.22E-05 | 0.000103 |
| Gm12680 | -3.31607 | 0.758229 | 1.22E-05 | 0.000103 |
| Ksr2 | -2.02159 | 0.4629 | 1.26E-05 | 0.000106 |

| | | | | |
|---|---|---|---|---|
| Duox2 | -2.81741 | 0.645733 | 1.28E-05 | 0.000107 |
| Ntm | -2.6882 | 0.616663 | 1.30E-05 | 0.000109 |
| Scube2 | -3.43272 | 0.788886 | 1.35E-05 | 0.000113 |
| Gm44293 | -2.86574 | 0.658698 | 1.36E-05 | 0.000113 |
| D030025P2 | 3.060767 | 0.70362 | 1.36E-05 | 0.000113 |
| Clspn | 4.235732 | 0.976363 | 1.44E-05 | 0.000119 |
| Oprk1 | -2.69168 | 0.621174 | 1.47E-05 | 0.000121 |
| Otof | -6.29286 | 1.452578 | 1.48E-05 | 0.000122 |
| Stmnd1 | -3.82716 | 0.883551 | 1.48E-05 | 0.000122 |
| 2310015D2 | -3.87565 | 0.895101 | 1.49E-05 | 0.000123 |
| Tubb3 | 2.044862 | 0.472559 | 1.51E-05 | 0.000124 |
| Gm15557 | 2.013255 | 0.465892 | 1.55E-05 | 0.000127 |
| Sprr2g | 5.051645 | 1.172535 | 1.65E-05 | 0.000134 |
| Eln | 3.791472 | 0.881042 | 1.68E-05 | 0.000136 |
| Bmp10 | -2.92109 | 0.682931 | 1.89E-05 | 0.000151 |
| Hist1h2aj | 3.863716 | 0.906546 | 2.03E-05 | 0.000161 |
| Gm26682 | 2.785641 | 0.654858 | 2.10E-05 | 0.000166 |
| Fam180a | -2.00406 | 0.472062 | 2.18E-05 | 0.000171 |
| Dmgdh | -2.68327 | 0.632441 | 2.21E-05 | 0.000173 |
| Adarb2 | -3.5441 | 0.835674 | 2.23E-05 | 0.000175 |
| Sox2ot | -3.17388 | 0.748419 | 2.23E-05 | 0.000175 |
| 4930562C1 | -5.13936 | 1.212361 | 2.24E-05 | 0.000176 |
| Depdc1b | 2.892803 | 0.683794 | 2.33E-05 | 0.000181 |
| Rsph4a | -3.75706 | 0.888816 | 2.37E-05 | 0.000184 |
| Serpina3n | 2.827436 | 0.66911 | 2.38E-05 | 0.000184 |
| Gm9767 | 2.382147 | 0.564966 | 2.48E-05 | 0.000191 |
| Gm2115 | -2.19515 | 0.522018 | 2.61E-05 | 0.000199 |
| Cenph | 3.560421 | 0.847653 | 2.67E-05 | 0.000203 |
| Tacc3 | 3.027063 | 0.720687 | 2.67E-05 | 0.000203 |
| Gm12867 | -3.89054 | 0.927197 | 2.72E-05 | 0.000206 |
| Crhbp | -2.49644 | 0.597245 | 2.92E-05 | 0.00022 |
| Hpcal4 | 2.878876 | 0.690352 | 3.04E-05 | 0.000228 |
| Gm23201 | 2.676912 | 0.642062 | 3.06E-05 | 0.000229 |
| Ccdc152 | -2.1741 | 0.521529 | 3.06E-05 | 0.000229 |
| Pigr | -2.16361 | 0.51902 | 3.06E-05 | 0.000229 |
| Fbxw19 | -6.13178 | 1.473254 | 3.15E-05 | 0.000235 |
| Pou2f3 | -2.12522 | 0.511349 | 3.24E-05 | 0.000241 |
| Gm29291 | -2.93931 | 0.707291 | 3.24E-05 | 0.000241 |
| Asf1b | 3.651748 | 0.880287 | 3.35E-05 | 0.000247 |
| Cd5l | 3.415729 | 0.823732 | 3.37E-05 | 0.000249 |
| 4930558J1 | 3.865966 | 0.932539 | 3.39E-05 | 0.00025 |
| Ppbp | 6.566823 | 1.585318 | 3.44E-05 | 0.000253 |
| Tg | 2.048155 | 0.494636 | 3.46E-05 | 0.000255 |
| Gm31406 | -3.33811 | 0.806636 | 3.50E-05 | 0.000257 |

| | | | | |
|---|---|---|---|---|
| Mafa | -3.1627 | 0.764477 | 3.52E-05 | 0.000258 |
| Gjb1 | 4.687611 | 1.134716 | 3.61E-05 | 0.000264 |
| Ttc23l | -2.45232 | 0.593904 | 3.64E-05 | 0.000266 |
| Cdca8 | 3.082368 | 0.74747 | 3.73E-05 | 0.000271 |
| Ccdc187 | -5.75527 | 1.397427 | 3.81E-05 | 0.000276 |
| Gm5067 | -2.72941 | 0.663077 | 3.85E-05 | 0.000279 |
| Siglech | -3.76907 | 0.915941 | 3.87E-05 | 0.00028 |
| Cd28 | -2.69244 | 0.654486 | 3.89E-05 | 0.000281 |
| Gm12603 | 4.246039 | 1.032263 | 3.90E-05 | 0.000281 |
| Snord16a | 2.529041 | 0.615416 | 3.97E-05 | 0.000286 |
| Dmbt1 | -2.71048 | 0.659963 | 4.01E-05 | 0.000288 |
| Hsd17b13 | -2.88147 | 0.701784 | 4.03E-05 | 0.000289 |
| Ltb | -2.46035 | 0.599234 | 4.03E-05 | 0.000289 |
| Efcab11 | 2.753249 | 0.670923 | 4.07E-05 | 0.000292 |
| Ccdc13 | -5.09311 | 1.241934 | 4.11E-05 | 0.000295 |
| Rbfox3 | -2.76248 | 0.673793 | 4.13E-05 | 0.000296 |
| Esrrg | -2.92011 | 0.712449 | 4.15E-05 | 0.000297 |
| Etv4 | 4.440899 | 1.084595 | 4.23E-05 | 0.000302 |
| Serpina3h | 3.045413 | 0.744611 | 4.31E-05 | 0.000307 |
| Gm7809 | 2.119583 | 0.518393 | 4.34E-05 | 0.000308 |
| Tmem240 | -2.05946 | 0.504185 | 4.41E-05 | 0.000313 |
| Aph1c | -2.07369 | 0.507783 | 4.43E-05 | 0.000314 |
| Chd5 | -2.0072 | 0.492625 | 4.61E-05 | 0.000325 |
| Nog | -2.08261 | 0.511252 | 4.63E-05 | 0.000326 |
| Serpina3f | 2.909386 | 0.714213 | 4.63E-05 | 0.000326 |
| Snord52 | 2.374237 | 0.583355 | 4.70E-05 | 0.000331 |
| Slitrk3 | -3.73139 | 0.9179 | 4.80E-05 | 0.000337 |
| Gm28402 | -2.34796 | 0.577585 | 4.80E-05 | 0.000337 |
| Adra1d | 2.292726 | 0.564457 | 4.87E-05 | 0.000341 |
| Pgk1-rs7 | 2.357562 | 0.581271 | 4.99E-05 | 0.000349 |
| Rnf112 | -3.25276 | 0.80219 | 5.02E-05 | 0.00035 |
| Gm45218 | -3.96623 | 0.981816 | 5.35E-05 | 0.00037 |
| Gm7361 | -3.33457 | 0.825537 | 5.36E-05 | 0.000371 |
| Mir8101 | -2.1231 | 0.526194 | 5.46E-05 | 0.000377 |
| Mroh7 | -3.20836 | 0.795512 | 5.51E-05 | 0.000379 |
| Klra2 | 2.212403 | 0.548686 | 5.53E-05 | 0.00038 |
| Gm5514 | -3.01832 | 0.748901 | 5.57E-05 | 0.000383 |
| Gm12961 | 2.138911 | 0.530965 | 5.62E-05 | 0.000385 |
| Plk1 | 2.884304 | 0.7167 | 5.71E-05 | 0.000391 |
| 0610033M | -2.31948 | 0.576421 | 5.72E-05 | 0.000391 |
| Gm44357 | 5.737049 | 1.427439 | 5.84E-05 | 0.000398 |
| Gprc5a | 3.270021 | 0.817463 | 6.33E-05 | 0.000427 |
| Hepacam2 | -3.21039 | 0.804713 | 6.62E-05 | 0.000444 |
| 1700020G | 5.25234 | 1.318859 | 6.82E-05 | 0.000456 |

| | | | | |
|---|---|---|---|---|
| Gm9974 | -2.58477 | 0.649282 | 6.86E-05 | 0.000458 |
| Slfn1 | 2.733603 | 0.687896 | 7.07E-05 | 0.000469 |
| Slco1a4 | -2.81084 | 0.707882 | 7.16E-05 | 0.000475 |
| Gm13270 | -2.05437 | 0.519161 | 7.59E-05 | 0.0005 |
| Slitrk5 | -2.35356 | 0.595246 | 7.69E-05 | 0.000505 |
| S100a3 | 2.114922 | 0.535678 | 7.88E-05 | 0.000516 |
| Gm13237 | 3.309331 | 0.838336 | 7.90E-05 | 0.000517 |
| 4932411E2 | -2.85671 | 0.723891 | 7.94E-05 | 0.000519 |
| Stfa2l1 | 6.232292 | 1.579366 | 7.94E-05 | 0.00052 |
| Gm42487 | -2.54832 | 0.646093 | 8.01E-05 | 0.000523 |
| Gm5483 | 6.36032 | 1.612721 | 8.02E-05 | 0.000524 |
| Diaph3 | 3.353626 | 0.850533 | 8.05E-05 | 0.000525 |
| Gm5648 | 5.620095 | 1.427499 | 8.25E-05 | 0.000536 |
| H2-Ob | -2.27837 | 0.580093 | 8.58E-05 | 0.000556 |
| Gm42706 | -2.73624 | 0.696739 | 8.59E-05 | 0.000557 |
| Gm15845 | 6.213789 | 1.582626 | 8.63E-05 | 0.000558 |
| Gm43136 | -2.26324 | 0.576548 | 8.65E-05 | 0.00056 |
| Bub1b | 2.693143 | 0.686642 | 8.77E-05 | 0.000567 |
| Fam183b | -2.07641 | 0.529762 | 8.87E-05 | 0.000572 |
| Gm37261 | -2.09565 | 0.53478 | 8.90E-05 | 0.000573 |
| Rnf182 | -3.24555 | 0.828776 | 9.00E-05 | 0.000578 |
| Mir7678 | 3.317786 | 0.84833 | 9.19E-05 | 0.000588 |
| D430036J1 | 2.139105 | 0.548495 | 9.62E-05 | 0.000613 |
| Car6 | 5.691911 | 1.460399 | 9.72E-05 | 0.000618 |
| Gm44737 | -5.4804 | 1.406383 | 9.75E-05 | 0.00062 |
| Fosl1 | 3.160357 | 0.811936 | 9.93E-05 | 0.00063 |
| Gm765 | -2.20374 | 0.566747 | 0.000101 | 0.000638 |
| Cnr1 | 2.023445 | 0.522679 | 0.000108 | 0.000679 |
| Gm6089 | -4.28323 | 1.10705 | 0.000109 | 0.000684 |
| A330048O | -2.8538 | 0.737974 | 0.00011 | 0.00069 |
| Gm44421 | -3.07917 | 0.796476 | 0.000111 | 0.000692 |
| AU018091 | 5.787283 | 1.498262 | 0.000112 | 0.000701 |
| Cyp4f15 | -3.31404 | 0.858783 | 0.000114 | 0.00071 |
| Phactr3 | -3.38484 | 0.877309 | 0.000114 | 0.000711 |
| Bsnd | -5.91559 | 1.537537 | 0.000119 | 0.000741 |
| S100a9 | 5.706893 | 1.485112 | 0.000122 | 0.000754 |
| Anln | 2.76062 | 0.720171 | 0.000126 | 0.000779 |
| Prnd | 2.197354 | 0.573936 | 0.000129 | 0.000791 |
| RP23-393E | 6.586395 | 1.720821 | 0.000129 | 0.000794 |
| Ccna2 | 2.754454 | 0.719766 | 0.00013 | 0.000796 |
| Gm15866 | 2.874856 | 0.751781 | 0.000131 | 0.000803 |
| Scg3 | -2.62968 | 0.687962 | 0.000132 | 0.000807 |
| Gm43534 | -4.17501 | 1.092697 | 0.000133 | 0.000812 |
| Gm4869 | -2.3204 | 0.607463 | 0.000134 | 0.000814 |

| | | | | |
|---|---|---|---|---|
| Gm43003 | 5.295327 | 1.391499 | 0.000142 | 0.000857 |
| Car4 | 2.511676 | 0.660165 | 0.000142 | 0.000859 |
| Hmga2 | 7.098209 | 1.871319 | 0.000149 | 0.000894 |
| Adamts17 | -2.01933 | 0.532652 | 0.00015 | 0.000901 |
| Ccl3 | 2.526991 | 0.666727 | 0.000151 | 0.000903 |
| Ntsr1 | -5.87645 | 1.551476 | 0.000152 | 0.000912 |
| Ccnf | 3.070346 | 0.811257 | 0.000154 | 0.000922 |
| Ifi202b | 3.561657 | 0.941336 | 0.000155 | 0.000925 |
| 2810417H1 | 4.11486 | 1.088424 | 0.000156 | 0.000935 |
| Kcnj5 | -4.26618 | 1.132579 | 0.000165 | 0.000981 |
| Lcn2 | 3.606093 | 0.95737 | 0.000165 | 0.000981 |
| Gm44390 | 3.889944 | 1.033373 | 0.000167 | 0.000989 |
| Gm24727 | 2.84308 | 0.756215 | 0.00017 | 0.001006 |
| S100a7a | -2.98366 | 0.79499 | 0.000175 | 0.00103 |
| Gm15984 | -2.00676 | 0.535194 | 0.000177 | 0.001042 |
| Mlf1 | 2.243784 | 0.598938 | 0.000179 | 0.001053 |
| Gm5593 | 3.282298 | 0.876453 | 0.00018 | 0.001058 |
| Hist1h2af | 3.931971 | 1.050002 | 0.000181 | 0.001059 |
| Gm10259 | 2.514571 | 0.673822 | 0.00019 | 0.001106 |
| Ly6i | 2.297386 | 0.616112 | 0.000192 | 0.001119 |
| Csta1 | 2.95684 | 0.79301 | 0.000193 | 0.001119 |
| Gm6377 | 2.270965 | 0.609125 | 0.000193 | 0.001121 |
| Prr11 | 3.078548 | 0.827126 | 0.000198 | 0.001145 |
| Ndp | -2.05851 | 0.553653 | 0.000201 | 0.00116 |
| Gm44514 | 2.477942 | 0.666778 | 0.000202 | 0.001167 |
| Mcemp1 | 2.49562 | 0.67168 | 0.000203 | 0.00117 |
| B430306N0 | 2.146972 | 0.577982 | 0.000204 | 0.001174 |
| Rad51ap1 | 3.613223 | 0.974634 | 0.00021 | 0.001204 |
| Fbxo48 | 2.665382 | 0.719487 | 0.000212 | 0.001214 |
| Catsperg1 | -2.0453 | 0.552161 | 0.000212 | 0.001216 |
| Myom2 | 3.970066 | 1.072195 | 0.000213 | 0.001222 |
| 9530062K0 | -2.39886 | 0.648423 | 0.000216 | 0.001235 |
| Gsdmcl-ps | 2.39849 | 0.648444 | 0.000217 | 0.001238 |
| Atp1a2 | -2.62472 | 0.70977 | 0.000217 | 0.001242 |
| Gpr3 | 3.828252 | 1.036231 | 0.00022 | 0.001257 |
| Gm43181 | 6.156602 | 1.668679 | 0.000225 | 0.00128 |
| Hs3st3b1 | 2.036831 | 0.552219 | 0.000226 | 0.001285 |
| Crocc2 | 2.659599 | 0.721515 | 0.000228 | 0.001295 |
| Gareml | 3.435539 | 0.93242 | 0.000229 | 0.001302 |
| Zfp879 | -2.26232 | 0.615019 | 0.000235 | 0.001325 |
| Ube2c | 3.035476 | 0.826294 | 0.000239 | 0.001349 |
| Espl1 | 2.809375 | 0.76536 | 0.000242 | 0.001362 |
| Sprr2f | 3.744335 | 1.020422 | 0.000243 | 0.001368 |
| Lgals1-ps2 | -3.39465 | 0.926561 | 0.000249 | 0.001396 |

| | | | | |
|---|---|---|---|---|
| Sgol2a | 2.964241 | 0.811717 | 0.00026 | 0.001452 |
| C330027C | 2.696336 | 0.73974 | 0.000267 | 0.001487 |
| Aurka | 2.835094 | 0.777889 | 0.000268 | 0.001488 |
| Mstn | -5.90866 | 1.623285 | 0.000273 | 0.001512 |
| Qrfpr | -3.83387 | 1.053332 | 0.000273 | 0.001513 |
| Crct1 | 6.235672 | 1.714519 | 0.000276 | 0.001527 |
| Myb | 2.557602 | 0.703555 | 0.000278 | 0.001537 |
| Gm15908 | 2.238751 | 0.616454 | 0.000282 | 0.001555 |
| Spdef | 2.079628 | 0.572817 | 0.000283 | 0.001561 |
| RP23-144N | 4.945135 | 1.366265 | 0.000295 | 0.001618 |
| Gm12856 | 5.382017 | 1.48816 | 0.000299 | 0.001633 |
| Gm12671 | 2.065326 | 0.571575 | 0.000302 | 0.001651 |
| H19 | 3.436309 | 0.951506 | 0.000305 | 0.001662 |
| Thsd7b | -4.84828 | 1.342867 | 0.000306 | 0.001668 |
| 1500012K( | -3.29166 | 0.914216 | 0.000318 | 0.001726 |
| A630075F1 | -2.24994 | 0.625816 | 0.000324 | 0.001755 |
| Gm44037 | -5.18282 | 1.444509 | 0.000333 | 0.001797 |
| Hist1h2ai | 3.682773 | 1.028653 | 0.000343 | 0.001846 |
| B230369F2 | -2.06539 | 0.577395 | 0.000347 | 0.001866 |
| Gm14548 | 2.252719 | 0.631922 | 0.000364 | 0.001944 |
| Tmem121 | 2.136589 | 0.599432 | 0.000365 | 0.001946 |
| Gm44346 | 4.942509 | 1.388305 | 0.000371 | 0.001973 |
| Itgb2l | -2.02797 | 0.570687 | 0.00038 | 0.002016 |
| Syt6 | 2.370051 | 0.667158 | 0.000382 | 0.002023 |
| Krt16 | 8.412118 | 2.369829 | 0.000386 | 0.00204 |
| Lrrc2 | 2.750826 | 0.775112 | 0.000387 | 0.002044 |
| Mir6999 | -4.60395 | 1.299998 | 0.000398 | 0.002095 |
| 2010300F1 | -3.69487 | 1.044928 | 0.000406 | 0.002129 |
| Asb10 | -4.6678 | 1.32027 | 0.000407 | 0.002132 |
| Lman1l | 3.045932 | 0.862707 | 0.000415 | 0.002166 |
| Ccl5 | -2.63259 | 0.745655 | 0.000415 | 0.002167 |
| Sbk2 | -3.95691 | 1.125873 | 0.000441 | 0.002279 |
| Cstad | -2.44619 | 0.696715 | 0.000446 | 0.002306 |
| Ect2 | 3.144657 | 0.895819 | 0.000447 | 0.002311 |
| Gpr165 | -5.34587 | 1.523449 | 0.00045 | 0.002322 |
| P2rx1 | -2.55617 | 0.728623 | 0.000451 | 0.002327 |
| Gm37309 | -2.00154 | 0.570524 | 0.000451 | 0.002327 |
| Guca1b | -2.06365 | 0.589374 | 0.000463 | 0.002377 |
| Gm22077 | 2.065024 | 0.590586 | 0.000471 | 0.002412 |
| Cdk1 | 3.62777 | 1.039289 | 0.000482 | 0.00246 |
| Gm27817 | -3.49446 | 1.002075 | 0.000488 | 0.002487 |
| Tmigd1 | 2.120529 | 0.608339 | 0.000491 | 0.0025 |
| Gm16118 | -2.92034 | 0.838586 | 0.000497 | 0.002526 |
| Gm15372 | -5.42346 | 1.563293 | 0.000522 | 0.002635 |

| | | | | |
|---|---|---|---|---|
| 2210418O | 2.492407 | 0.71887 | 0.000526 | 0.002654 |
| Wnt9b | 3.77004 | 1.091816 | 0.000554 | 0.002772 |
| BC030867 | 3.99492 | 1.157448 | 0.000558 | 0.002785 |
| Pgr | -2.12799 | 0.616842 | 0.000561 | 0.002798 |
| Gabrd | -2.48348 | 0.72036 | 0.000566 | 0.002817 |
| Klhl10 | -2.4982 | 0.725115 | 0.000571 | 0.002838 |
| Rnf212 | -4.14377 | 1.203647 | 0.000576 | 0.002862 |
| Aoc1 | 2.943255 | 0.85518 | 0.000578 | 0.002872 |
| Plcxd3 | -2.68703 | 0.780973 | 0.00058 | 0.002882 |
| Gdnf | 2.834073 | 0.823925 | 0.000582 | 0.002891 |
| Slc26a8 | -2.11617 | 0.615559 | 0.000586 | 0.002907 |
| Wfdc6a | -5.29766 | 1.541461 | 0.000589 | 0.002916 |
| Glrp1 | 6.054689 | 1.764279 | 0.0006 | 0.002962 |
| Dlgap5 | 2.870593 | 0.836578 | 0.000601 | 0.002966 |
| Gm27337 | -4.42611 | 1.292086 | 0.000614 | 0.003021 |
| Dclk3 | -2.9847 | 0.873049 | 0.000629 | 0.003088 |
| 0610012D( | 3.110544 | 0.909899 | 0.00063 | 0.003089 |
| Gm14024 | 3.760187 | 1.10084 | 0.000636 | 0.003117 |
| Calca | 2.328101 | 0.682752 | 0.00065 | 0.003174 |
| Zp1 | -3.03025 | 0.891795 | 0.000679 | 0.003299 |
| Tnnc2 | -2.1027 | 0.61976 | 0.000692 | 0.003352 |
| Lrrtm3 | -2.21535 | 0.652995 | 0.000692 | 0.003354 |
| Gm6293 | -2.10994 | 0.622221 | 0.000696 | 0.003369 |
| Gm26580 | -2.61811 | 0.772914 | 0.000706 | 0.003409 |
| Nusap1 | 2.875472 | 0.851545 | 0.000733 | 0.003526 |
| Gm26847 | 2.108163 | 0.624344 | 0.000734 | 0.003526 |
| Gm15271 | 5.512438 | 1.633528 | 0.000739 | 0.00355 |
| Hapln1 | -2.33643 | 0.693758 | 0.000758 | 0.003624 |
| Mei1 | -2.58643 | 0.768453 | 0.000763 | 0.003648 |
| Foxm1 | 3.154709 | 0.940526 | 0.000796 | 0.003779 |
| Alms1-ps1 | -2.59492 | 0.774357 | 0.000805 | 0.003817 |
| Wisp2 | 2.764371 | 0.825264 | 0.000809 | 0.003834 |
| Cdca5 | 3.637401 | 1.086599 | 0.000815 | 0.003859 |
| Kifc1 | 3.33348 | 0.997008 | 0.000827 | 0.003911 |
| Prr32 | 2.406115 | 0.72012 | 0.000834 | 0.003935 |
| Gm1110 | -5.46864 | 1.640665 | 0.000859 | 0.004035 |
| Gm11832 | 2.335962 | 0.701275 | 0.000865 | 0.004062 |
| 3830417A1 | -2.38992 | 0.717605 | 0.000867 | 0.004068 |
| Gm11697 | -2.01409 | 0.605258 | 0.000876 | 0.0041 |
| Tnf | 3.387456 | 1.018751 | 0.000884 | 0.00413 |
| D830026I1 | -2.02696 | 0.609777 | 0.000887 | 0.004143 |
| Gm13111 | -2.46596 | 0.74306 | 0.000905 | 0.004216 |
| Ucn2 | 3.26442 | 0.983652 | 0.000904 | 0.004216 |
| Tcerg1l | -2.65832 | 0.801551 | 0.000912 | 0.004241 |

| | | | | |
|---|---|---|---|---|
| Cyp2c44 | -2.37786 | 0.717193 | 0.000915 | 0.004252 |
| Plxna4os3 | -3.59328 | 1.084574 | 0.000923 | 0.004281 |
| Kntc1 | 3.744814 | 1.130495 | 0.000925 | 0.004287 |
| 2310040G[illegible] | -4.96965 | 1.501952 | 0.000937 | 0.004333 |
| Gm5596 | 5.707808 | 1.726148 | 0.000944 | 0.004365 |
| Hmgcs2 | -2.75526 | 0.833678 | 0.00095 | 0.00439 |
| Gabrp | 2.766139 | 0.84044 | 0.000997 | 0.004583 |

Supplemental Table 2. d7 DEGs

| GeneID | Log2 fc | SE | p-value | FDR p-value |
|---|---|---|---|---|
| Inhba | 3.282637 | 0.304112 | 3.67E-27 | 8.35E-23 |
| Cxcl5 | 5.61521 | 0.536896 | 1.34E-25 | 1.52E-21 |
| Tmem200a | 2.637814 | 0.260047 | 3.54E-24 | 2.68E-20 |
| Clec4d | 3.808857 | 0.380489 | 1.37E-23 | 7.80E-20 |
| Arg1 | 6.196392 | 0.626263 | 4.41E-23 | 2.01E-19 |
| Pcp4 | -4.09649 | 0.424829 | 5.28E-22 | 2.00E-18 |
| Lox | 3.072982 | 0.329907 | 1.22E-20 | 3.48E-17 |
| Sox11 | 4.3975 | 0.471413 | 1.08E-20 | 3.48E-17 |
| AA467197 | 3.653695 | 0.399625 | 6.09E-20 | 1.54E-16 |
| Sulf1 | 1.411077 | 0.156386 | 1.83E-19 | 4.16E-16 |
| Cemip | 4.546743 | 0.52999 | 9.58E-18 | 1.98E-14 |
| Crlf1 | 4.009638 | 0.4694 | 1.32E-17 | 2.50E-14 |
| Ddah1 | 3.001801 | 0.351862 | 1.45E-17 | 2.54E-14 |
| Bmpr1b | 1.188136 | 0.140057 | 2.19E-17 | 3.56E-14 |
| Pmepa1 | 1.314989 | 0.15576 | 3.11E-17 | 4.49E-14 |
| Tmprss11g | 4.628407 | 0.548347 | 3.16E-17 | 4.49E-14 |
| Gm12405 | -2.57547 | 0.308589 | 7.06E-17 | 9.46E-14 |
| 4921507P( | -2.12167 | 0.254556 | 7.77E-17 | 9.82E-14 |
| Psrc1 | 2.36775 | 0.287692 | 1.87E-16 | 2.24E-13 |
| Ms4a6d | 1.989262 | 0.248298 | 1.13E-15 | 1.23E-12 |
| Ckap2 | 3.36914 | 0.422196 | 1.46E-15 | 1.51E-12 |
| Timp1 | 4.209967 | 0.530188 | 2.01E-15 | 1.99E-12 |
| Zfas1 | 1.312882 | 0.166204 | 2.81E-15 | 2.66E-12 |
| Bmper | 3.765587 | 0.481392 | 5.19E-15 | 4.72E-12 |
| 2810417H | 4.812091 | 0.616615 | 6.00E-15 | 5.25E-12 |
| Kcnk12 | 8.854109 | 1.135723 | 6.39E-15 | 5.39E-12 |
| Thbs1 | 3.075611 | 0.394959 | 6.85E-15 | 5.57E-12 |
| Rims1 | -2.66958 | 0.343373 | 7.57E-15 | 5.94E-12 |
| Gjb1 | 4.548505 | 0.585967 | 8.33E-15 | 6.32E-12 |
| Tubb3 | 2.578104 | 0.332346 | 8.68E-15 | 6.37E-12 |
| Wisp1 | 2.754796 | 0.357828 | 1.38E-14 | 9.78E-12 |
| C430002N | 5.60326 | 0.734618 | 2.39E-14 | 1.65E-11 |
| Gdf6 | 2.260494 | 0.297095 | 2.77E-14 | 1.80E-11 |
| Foxj1 | 2.707633 | 0.35571 | 2.70E-14 | 1.80E-11 |
| Clic6 | 2.94691 | 0.388674 | 3.40E-14 | 2.12E-11 |
| Grik3 | -2.37034 | 0.313511 | 4.01E-14 | 2.40E-11 |
| Gm29233 | 2.73133 | 0.36152 | 4.18E-14 | 2.44E-11 |
| Nes | 2.319431 | 0.307155 | 4.31E-14 | 2.45E-11 |
| Kcng3 | -2.3807 | 0.316325 | 5.23E-14 | 2.90E-11 |
| Dio3 | 3.250819 | 0.43242 | 5.57E-14 | 2.95E-11 |
| Ctgf | 4.702876 | 0.625523 | 5.55E-14 | 2.95E-11 |
| Spock3 | -3.71393 | 0.494343 | 5.79E-14 | 2.99E-11 |

| | | | | |
|---|---|---|---|---|
| Gm15270 | 4.508608 | 0.600408 | 5.95E-14 | 3.01E-11 |
| Cpne6 | -4.4746 | 0.599079 | 8.07E-14 | 3.92E-11 |
| Nxph3 | 2.004126 | 0.268332 | 8.09E-14 | 3.92E-11 |
| Fst | 2.948429 | 0.395424 | 8.89E-14 | 4.22E-11 |
| Igsf5 | -1.70987 | 0.229901 | 1.03E-13 | 4.76E-11 |
| Ptx3 | 5.203667 | 0.699886 | 1.05E-13 | 4.76E-11 |
| Gm11816 | -4.16136 | 0.562034 | 1.32E-13 | 5.89E-11 |
| Hs6st2 | 3.154162 | 0.426503 | 1.41E-13 | 6.17E-11 |
| Pitpnm3 | -1.41003 | 0.192065 | 2.11E-13 | 9.08E-11 |
| Ankrd63 | -3.69411 | 0.506463 | 3.01E-13 | 1.27E-10 |
| Bdkrb1 | 5.30107 | 0.727022 | 3.07E-13 | 1.27E-10 |
| Pcsk5 | 2.194761 | 0.30116 | 3.15E-13 | 1.28E-10 |
| Pla1a | 1.596823 | 0.219781 | 3.72E-13 | 1.48E-10 |
| Cdkn2a | 2.005974 | 0.277606 | 4.97E-13 | 1.95E-10 |
| Deptor | -1.04393 | 0.144737 | 5.49E-13 | 2.12E-10 |
| Adamts4 | 2.962296 | 0.411957 | 6.44E-13 | 2.44E-10 |
| Nt5dc2 | 2.158585 | 0.300352 | 6.63E-13 | 2.47E-10 |
| Mybl1 | 2.211923 | 0.308385 | 7.36E-13 | 2.70E-10 |
| Tmem200c | -2.68979 | 0.37618 | 8.66E-13 | 3.13E-10 |
| Enpp2 | -1.19604 | 0.169711 | 1.82E-12 | 6.48E-10 |
| 4932411E2 | -1.96114 | 0.279409 | 2.24E-12 | 7.81E-10 |
| Adam12 | 2.448205 | 0.34889 | 2.26E-12 | 7.81E-10 |
| Anln | 2.734697 | 0.389994 | 2.35E-12 | 7.97E-10 |
| Gm44238 | 3.039664 | 0.436277 | 3.23E-12 | 1.07E-09 |
| Kit | -2.59453 | 0.375128 | 4.63E-12 | 1.51E-09 |
| Ndc80 | 4.274999 | 0.618986 | 4.97E-12 | 1.59E-09 |
| Olr1 | 6.344351 | 0.920312 | 5.44E-12 | 1.72E-09 |
| Cenpe | 4.111552 | 0.598801 | 6.59E-12 | 2.05E-09 |
| Cdh4 | -1.86107 | 0.271728 | 7.43E-12 | 2.29E-09 |
| Aox3 | -3.00633 | 0.439316 | 7.74E-12 | 2.35E-09 |
| Adora1 | -1.36133 | 0.199099 | 8.06E-12 | 2.41E-09 |
| Retnla | -3.30845 | 0.485392 | 9.36E-12 | 2.74E-09 |
| Kif2c | 4.529845 | 0.664617 | 9.38E-12 | 2.74E-09 |
| Nuf2 | 4.254159 | 0.624773 | 9.82E-12 | 2.83E-09 |
| Tpx2 | 3.5604 | 0.52344 | 1.03E-11 | 2.94E-09 |
| Gria1 | -3.90505 | 0.575967 | 1.20E-11 | 3.38E-09 |
| Nptx1 | -2.7857 | 0.411059 | 1.23E-11 | 3.41E-09 |
| Smim3 | 1.596192 | 0.235768 | 1.29E-11 | 3.53E-09 |
| Rasd2 | -1.42956 | 0.213848 | 2.31E-11 | 6.19E-09 |
| Spp1 | 4.386549 | 0.657585 | 2.55E-11 | 6.74E-09 |
| Asic2 | -3.05214 | 0.459061 | 2.96E-11 | 7.74E-09 |
| C7 | -2.55104 | 0.383968 | 3.06E-11 | 7.90E-09 |
| Adam8 | 2.109112 | 0.318344 | 3.47E-11 | 8.87E-09 |
| Scd1 | -1.69728 | 0.256281 | 3.53E-11 | 8.92E-09 |

| | | | | |
|---|---|---|---|---|
| Gm28857 | -1.22325 | 0.184837 | 3.64E-11 | 9.11E-09 |
| Fhl3 | 2.482416 | 0.375196 | 3.68E-11 | 9.11E-09 |
| Csgalnact1 | 2.318974 | 0.350929 | 3.89E-11 | 9.53E-09 |
| Mex3a | 1.07526 | 0.163089 | 4.31E-11 | 1.03E-08 |
| Plscr1 | 1.514743 | 0.229794 | 4.35E-11 | 1.03E-08 |
| Cep55 | 4.272124 | 0.647998 | 4.32E-11 | 1.03E-08 |
| Arl4c | 1.259676 | 0.191175 | 4.42E-11 | 1.04E-08 |
| Lockd | 2.629005 | 0.399505 | 4.68E-11 | 1.09E-08 |
| Lat2 | 1.062633 | 0.161564 | 4.80E-11 | 1.10E-08 |
| Il1b | 4.498586 | 0.684422 | 4.94E-11 | 1.12E-08 |
| Ccl7 | 4.157888 | 0.632827 | 5.02E-11 | 1.13E-08 |
| Serpina3n | 2.08292 | 0.318752 | 6.38E-11 | 1.42E-08 |
| Ltb4r2 | 4.702287 | 0.721471 | 7.14E-11 | 1.58E-08 |
| Cenph | 4.014049 | 0.616813 | 7.63E-11 | 1.67E-08 |
| Gap43 | 1.981578 | 0.305172 | 8.40E-11 | 1.80E-08 |
| Snx7 | 1.276749 | 0.196806 | 8.74E-11 | 1.86E-08 |
| Gm9920 | -1.25497 | 0.193732 | 9.30E-11 | 1.96E-08 |
| Hcrtr1 | -2.31844 | 0.358279 | 9.73E-11 | 2.03E-08 |
| Ngb | -3.51174 | 0.543265 | 1.02E-10 | 2.09E-08 |
| Grip2 | -2.7101 | 0.421104 | 1.23E-10 | 2.45E-08 |
| Acvr1c | -2.28065 | 0.354337 | 1.22E-10 | 2.45E-08 |
| Egr2 | 3.470549 | 0.539414 | 1.24E-10 | 2.46E-08 |
| Mir147 | 5.250792 | 0.816977 | 1.30E-10 | 2.55E-08 |
| Shank2 | -1.22156 | 0.190728 | 1.51E-10 | 2.91E-08 |
| Sostdc1 | -2.11715 | 0.330747 | 1.54E-10 | 2.95E-08 |
| Spsb1 | 1.332737 | 0.208645 | 1.69E-10 | 3.17E-08 |
| Kifc1 | 4.012924 | 0.628232 | 1.68E-10 | 3.17E-08 |
| Reg3g | 5.927924 | 0.928357 | 1.71E-10 | 3.19E-08 |
| Socs3 | 3.928583 | 0.615408 | 1.73E-10 | 3.20E-08 |
| Camk2n2 | 1.363708 | 0.213755 | 1.77E-10 | 3.26E-08 |
| Cxcr2 | 3.426242 | 0.538555 | 1.99E-10 | 3.63E-08 |
| Chst11 | 1.581633 | 0.248885 | 2.09E-10 | 3.77E-08 |
| Hlf | -2.67281 | 0.421167 | 2.21E-10 | 3.94E-08 |
| Ccl2 | 3.517369 | 0.554286 | 2.21E-10 | 3.94E-08 |
| Shcbp1 | 4.323191 | 0.681425 | 2.23E-10 | 3.94E-08 |
| Pdia5 | 1.145737 | 0.180804 | 2.34E-10 | 4.09E-08 |
| Frzb | 2.265489 | 0.35754 | 2.35E-10 | 4.09E-08 |
| Fkbp11 | 2.077221 | 0.328003 | 2.41E-10 | 4.15E-08 |
| Mical2 | 1.400678 | 0.221329 | 2.48E-10 | 4.24E-08 |
| Spc24 | 3.629331 | 0.574107 | 2.59E-10 | 4.39E-08 |
| Tmem173 | 1.744627 | 0.276143 | 2.65E-10 | 4.41E-08 |
| Casc5 | 4.197859 | 0.664381 | 2.64E-10 | 4.41E-08 |
| Il11 | 4.437348 | 0.702313 | 2.65E-10 | 4.41E-08 |
| Gm27483 | 3.153048 | 0.499247 | 2.69E-10 | 4.44E-08 |

| | | | | |
|---|---|---|---|---|
| Nucb2 | 1.215616 | 0.19321 | 3.14E-10 | 5.14E-08 |
| Tgm1 | 3.702336 | 0.588757 | 3.21E-10 | 5.22E-08 |
| Mir675 | 3.101454 | 0.494185 | 3.48E-10 | 5.61E-08 |
| Zfhx2os | -1.99774 | 0.318465 | 3.54E-10 | 5.68E-08 |
| Wif1 | -1.52761 | 0.244341 | 4.05E-10 | 6.45E-08 |
| Pif1 | 3.823768 | 0.612498 | 4.30E-10 | 6.79E-08 |
| RP23-235I: | 1.482816 | 0.237565 | 4.33E-10 | 6.79E-08 |
| Gabrp | 3.757203 | 0.603901 | 4.92E-10 | 7.67E-08 |
| Cdc20 | 3.969962 | 0.638637 | 5.09E-10 | 7.88E-08 |
| Cdc25c | 4.062551 | 0.657035 | 6.28E-10 | 9.66E-08 |
| Scn4b | -1.6388 | 0.265292 | 6.52E-10 | 9.96E-08 |
| Racgap1 | 3.697357 | 0.598854 | 6.66E-10 | 1.01E-07 |
| Htr1b | 2.152733 | 0.35021 | 7.90E-10 | 1.19E-07 |
| Kl | -1.67468 | 0.273371 | 9.01E-10 | 1.34E-07 |
| Exph5 | -1.08048 | 0.176605 | 9.47E-10 | 1.40E-07 |
| Etv4 | 4.654257 | 0.761386 | 9.79E-10 | 1.44E-07 |
| Akap6 | -2.11314 | 0.346313 | 1.05E-09 | 1.53E-07 |
| 11-Sep | 1.374651 | 0.225621 | 1.11E-09 | 1.60E-07 |
| Prc1 | 3.679896 | 0.6039 | 1.10E-09 | 1.60E-07 |
| Dok7 | -2.09674 | 0.345847 | 1.34E-09 | 1.92E-07 |
| Hmmr | 3.951605 | 0.653386 | 1.47E-09 | 2.09E-07 |
| Gpr176 | 2.184919 | 0.361377 | 1.48E-09 | 2.10E-07 |
| Rnd1 | 4.122357 | 0.682641 | 1.55E-09 | 2.18E-07 |
| Diaph3 | 3.447128 | 0.571506 | 1.62E-09 | 2.27E-07 |
| H19 | 2.781019 | 0.461261 | 1.65E-09 | 2.29E-07 |
| Kif5a | -1.7405 | 0.289023 | 1.72E-09 | 2.38E-07 |
| Smad9 | -2.63284 | 0.437516 | 1.77E-09 | 2.43E-07 |
| Evx2 | -1.78836 | 0.298024 | 1.96E-09 | 2.68E-07 |
| Spc25 | 2.944395 | 0.491032 | 2.02E-09 | 2.73E-07 |
| Gm28729 | -3.43647 | 0.574236 | 2.17E-09 | 2.92E-07 |
| Depdc1a | 4.099085 | 0.685449 | 2.23E-09 | 2.98E-07 |
| Serpina3i | 3.691455 | 0.617566 | 2.27E-09 | 3.02E-07 |
| Cenpf | 3.843204 | 0.643816 | 2.38E-09 | 3.15E-07 |
| Cdh6 | -1.90056 | 0.319169 | 2.61E-09 | 3.43E-07 |
| Areg | 4.957474 | 0.833508 | 2.72E-09 | 3.56E-07 |
| Rgs16 | 2.979957 | 0.5017 | 2.86E-09 | 3.69E-07 |
| Gm12406 | -2.52898 | 0.426101 | 2.94E-09 | 3.77E-07 |
| Fam150a | 5.706232 | 0.964042 | 3.24E-09 | 4.09E-07 |
| Osm | 3.146773 | 0.531875 | 3.29E-09 | 4.12E-07 |
| Kif22 | 3.890857 | 0.657633 | 3.29E-09 | 4.12E-07 |
| Cidea | -3.57422 | 0.604395 | 3.34E-09 | 4.16E-07 |
| Ifi202b | 4.866899 | 0.826964 | 3.97E-09 | 4.89E-07 |
| Cldn10 | -2.33085 | 0.396137 | 4.01E-09 | 4.89E-07 |
| Lilr4b | 1.409484 | 0.239572 | 4.02E-09 | 4.89E-07 |

| | | | | |
|---|---|---|---|---|
| E2f1 | 2.318982 | 0.395375 | 4.48E-09 | 5.43E-07 |
| Rragd | -1.3003 | 0.22203 | 4.73E-09 | 5.68E-07 |
| Gm14024 | 3.655852 | 0.624281 | 4.74E-09 | 5.68E-07 |
| Depdc1b | 4.077822 | 0.696774 | 4.84E-09 | 5.77E-07 |
| Cenpi | 4.139287 | 0.710248 | 5.61E-09 | 6.62E-07 |
| Ccne1 | 3.662571 | 0.629 | 5.79E-09 | 6.79E-07 |
| Ctla2b | 1.409242 | 0.242072 | 5.83E-09 | 6.80E-07 |
| Tpm4 | 1.64623 | 0.282918 | 5.93E-09 | 6.89E-07 |
| Nek2 | 3.764144 | 0.647741 | 6.20E-09 | 7.17E-07 |
| 3300005D( | 3.061772 | 0.52781 | 6.60E-09 | 7.58E-07 |
| Cytl1 | -2.27389 | 0.392092 | 6.66E-09 | 7.61E-07 |
| Ctla2a | 1.488291 | 0.257382 | 7.36E-09 | 8.38E-07 |
| Ryr2 | -2.06412 | 0.357224 | 7.55E-09 | 8.55E-07 |
| Myoc | -3.25244 | 0.563374 | 7.78E-09 | 8.72E-07 |
| Il21r | 1.694211 | 0.293447 | 7.77E-09 | 8.72E-07 |
| Spdl1 | 3.059445 | 0.53082 | 8.23E-09 | 9.19E-07 |
| Dhtkd1 | -1.22496 | 0.212944 | 8.79E-09 | 9.76E-07 |
| Mafa | -2.6983 | 0.46943 | 9.03E-09 | 9.98E-07 |
| Fam64a | 3.294535 | 0.573561 | 9.25E-09 | 1.02E-06 |
| Nr3c2 | -1.51362 | 0.263744 | 9.52E-09 | 1.04E-06 |
| Gm14091 | 3.444731 | 0.600477 | 9.66E-09 | 1.05E-06 |
| Atp2a3 | -1.55106 | 0.270441 | 9.73E-09 | 1.05E-06 |
| Cdkn3 | 3.618496 | 0.631008 | 9.78E-09 | 1.06E-06 |
| Il6 | 7.000582 | 1.222088 | 1.01E-08 | 1.09E-06 |
| Gadd45g | 2.238741 | 0.391294 | 1.06E-08 | 1.12E-06 |
| Serpine1 | 5.466161 | 0.955843 | 1.07E-08 | 1.14E-06 |
| Phlda2 | -1.91374 | 0.335316 | 1.15E-08 | 1.21E-06 |
| Ccr5 | 1.503366 | 0.263499 | 1.16E-08 | 1.22E-06 |
| Rhoc | 1.139149 | 0.199897 | 1.21E-08 | 1.24E-06 |
| Foxs1 | 2.802074 | 0.491462 | 1.19E-08 | 1.24E-06 |
| Gm27786 | 3.146631 | 0.552027 | 1.20E-08 | 1.24E-06 |
| Epgn | 5.028176 | 0.882196 | 1.20E-08 | 1.24E-06 |
| Myrip | -1.27228 | 0.223386 | 1.23E-08 | 1.26E-06 |
| Cenpa | 3.162043 | 0.556121 | 1.30E-08 | 1.32E-06 |
| F2r | 1.384279 | 0.243815 | 1.37E-08 | 1.38E-06 |
| Gm28513 | 2.890465 | 0.509161 | 1.37E-08 | 1.38E-06 |
| Knstrn | 3.256233 | 0.573965 | 1.40E-08 | 1.40E-06 |
| Gm20726 | 2.325653 | 0.410047 | 1.41E-08 | 1.41E-06 |
| Gsn | -1.23926 | 0.219328 | 1.60E-08 | 1.59E-06 |
| Ece2 | 1.039889 | 0.184222 | 1.65E-08 | 1.63E-06 |
| Itgam | 1.093509 | 0.194644 | 1.93E-08 | 1.90E-06 |
| Slc41a2 | 1.087969 | 0.193931 | 2.02E-08 | 1.98E-06 |
| Gadd45b | 1.772961 | 0.316123 | 2.04E-08 | 1.99E-06 |
| Tspan4 | 1.492657 | 0.266425 | 2.11E-08 | 2.05E-06 |

| | | | | |
|---|---|---|---|---|
| E2f7 | 3.342037 | 0.597097 | 2.18E-08 | 2.10E-06 |
| Ripk3 | 2.987297 | 0.53415 | 2.24E-08 | 2.15E-06 |
| Ccna2 | 2.984231 | 0.533895 | 2.28E-08 | 2.17E-06 |
| Gm38037 | 3.896149 | 0.696972 | 2.27E-08 | 2.17E-06 |
| Sv2b | -2.76634 | 0.495229 | 2.32E-08 | 2.20E-06 |
| Ccdc64 | -1.66608 | 0.299171 | 2.56E-08 | 2.42E-06 |
| Gm15947 | 2.294394 | 0.412555 | 2.68E-08 | 2.51E-06 |
| Ostc | 1.053377 | 0.18951 | 2.72E-08 | 2.54E-06 |
| Lgmn | 1.132682 | 0.203861 | 2.76E-08 | 2.56E-06 |
| Hmgcs2 | -3.05181 | 0.54946 | 2.79E-08 | 2.58E-06 |
| Zcchc5 | -1.68343 | 0.303162 | 2.81E-08 | 2.59E-06 |
| Neil3 | 3.592068 | 0.647058 | 2.83E-08 | 2.59E-06 |
| Reg1 | 5.349195 | 0.963483 | 2.83E-08 | 2.59E-06 |
| Prss32 | 2.272822 | 0.40963 | 2.88E-08 | 2.62E-06 |
| Stc1 | -1.78402 | 0.322014 | 3.02E-08 | 2.73E-06 |
| Gdf15 | 3.536887 | 0.638479 | 3.03E-08 | 2.73E-06 |
| Saa3 | 3.865078 | 0.698556 | 3.15E-08 | 2.82E-06 |
| Ckap2l | 3.311788 | 0.598734 | 3.18E-08 | 2.84E-06 |
| Sycp3 | -1.34266 | 0.24284 | 3.22E-08 | 2.86E-06 |
| Aspm | 3.725928 | 0.673981 | 3.23E-08 | 2.86E-06 |
| Krt14 | 6.124575 | 1.108664 | 3.31E-08 | 2.92E-06 |
| Gm26586 | 1.724836 | 0.312305 | 3.33E-08 | 2.93E-06 |
| Ccdc34 | 1.684798 | 0.305127 | 3.36E-08 | 2.94E-06 |
| Ckap4 | 1.276399 | 0.231284 | 3.41E-08 | 2.98E-06 |
| Cxcl2 | 4.316149 | 0.78338 | 3.60E-08 | 3.12E-06 |
| Siglech | -2.80152 | 0.509483 | 3.82E-08 | 3.31E-06 |
| Bub1 | 4.686509 | 0.852837 | 3.90E-08 | 3.36E-06 |
| Ear2 | -2.56344 | 0.466644 | 3.94E-08 | 3.39E-06 |
| 1700019D( | -1.62984 | 0.296919 | 4.04E-08 | 3.46E-06 |
| Nusap1 | 3.519385 | 0.641267 | 4.06E-08 | 3.46E-06 |
| Ch25h | 3.390174 | 0.618056 | 4.13E-08 | 3.51E-06 |
| Kcna2 | -2.80581 | 0.511813 | 4.20E-08 | 3.53E-06 |
| Trdn | -2.3862 | 0.435247 | 4.20E-08 | 3.53E-06 |
| Dlgap5 | 3.096675 | 0.566294 | 4.54E-08 | 3.80E-06 |
| Lgals1 | 1.588914 | 0.290631 | 4.57E-08 | 3.81E-06 |
| Ccnb2 | 4.526353 | 0.830273 | 4.99E-08 | 4.15E-06 |
| 1110038B1 | 1.501042 | 0.275692 | 5.19E-08 | 4.30E-06 |
| Mtfr2 | 4.020394 | 0.739312 | 5.39E-08 | 4.44E-06 |
| Cacna2d3 | -1.72561 | 0.317718 | 5.60E-08 | 4.60E-06 |
| Gpr55 | 4.348209 | 0.800777 | 5.64E-08 | 4.61E-06 |
| Slc9a2 | -1.38704 | 0.255829 | 5.90E-08 | 4.81E-06 |
| Cdca8 | 3.379662 | 0.623932 | 6.07E-08 | 4.92E-06 |
| 4930447N( | -1.97259 | 0.364535 | 6.26E-08 | 5.05E-06 |
| Mir6950 | 2.243237 | 0.414683 | 6.32E-08 | 5.08E-06 |

| | | | | |
|---|---|---|---|---|
| Brca1 | 3.535416 | 0.654507 | 6.60E-08 | 5.29E-06 |
| 4930524O( | 1.323146 | 0.244993 | 6.64E-08 | 5.30E-06 |
| Ptrh1 | 1.536212 | 0.284522 | 6.69E-08 | 5.32E-06 |
| Gm42793 | 4.642456 | 0.860803 | 6.92E-08 | 5.49E-06 |
| Relt | 2.348325 | 0.436235 | 7.32E-08 | 5.78E-06 |
| Atcayos | -1.83724 | 0.341698 | 7.58E-08 | 5.95E-06 |
| Tnfsf9 | 2.078599 | 0.386558 | 7.57E-08 | 5.95E-06 |
| 4930579G: | 1.334386 | 0.248258 | 7.66E-08 | 5.99E-06 |
| Cdk1 | 4.311688 | 0.802592 | 7.78E-08 | 6.06E-06 |
| Mrvi1 | -1.13734 | 0.211803 | 7.88E-08 | 6.12E-06 |
| Slc35f1 | -1.4668 | 0.27333 | 8.03E-08 | 6.22E-06 |
| Parpbp | 3.043779 | 0.567339 | 8.09E-08 | 6.25E-06 |
| BC100530 | 6.012075 | 1.123145 | 8.66E-08 | 6.66E-06 |
| Clec4e | 4.633039 | 0.865623 | 8.69E-08 | 6.66E-06 |
| Loxl2 | 2.004593 | 0.374642 | 8.76E-08 | 6.69E-06 |
| Galnt15 | -1.89601 | 0.355114 | 9.34E-08 | 7.11E-06 |
| Galr2 | -2.90612 | 0.545429 | 9.92E-08 | 7.50E-06 |
| Tacc3 | 3.245269 | 0.609084 | 9.92E-08 | 7.50E-06 |
| Cenpk | 3.009219 | 0.565139 | 1.01E-07 | 7.59E-06 |
| Tnfaip6 | 3.375383 | 0.633857 | 1.01E-07 | 7.59E-06 |
| Lrrc15 | 2.59711 | 0.487842 | 1.02E-07 | 7.61E-06 |
| BC025446 | 3.561919 | 0.669177 | 1.02E-07 | 7.62E-06 |
| Mmp7 | 8.004296 | 1.503956 | 1.03E-07 | 7.63E-06 |
| Gm44342 | 6.04281 | 1.135895 | 1.04E-07 | 7.70E-06 |
| Ska1 | 3.807222 | 0.71662 | 1.08E-07 | 7.97E-06 |
| Pbk | 4.893048 | 0.921197 | 1.09E-07 | 7.98E-06 |
| B4galt5 | 1.273786 | 0.239988 | 1.11E-07 | 8.10E-06 |
| Cadm2 | -1.5191 | 0.286501 | 1.14E-07 | 8.32E-06 |
| Rab17 | -2.69686 | 0.509223 | 1.18E-07 | 8.55E-06 |
| Gsg2 | 3.6919 | 0.697555 | 1.21E-07 | 8.68E-06 |
| Ms4a4a | 1.47111 | 0.278027 | 1.21E-07 | 8.70E-06 |
| Esco2 | 4.712329 | 0.890568 | 1.21E-07 | 8.70E-06 |
| Ltbp1 | 1.187809 | 0.224689 | 1.25E-07 | 8.90E-06 |
| Ncapg2 | 2.995852 | 0.566819 | 1.25E-07 | 8.92E-06 |
| Ccnb1 | 4.660775 | 0.881972 | 1.26E-07 | 8.94E-06 |
| Slc8a3 | -1.7783 | 0.336794 | 1.29E-07 | 9.13E-06 |
| Serpina3m | 3.719015 | 0.704551 | 1.30E-07 | 9.18E-06 |
| Lsamp | -2.58643 | 0.490528 | 1.34E-07 | 9.44E-06 |
| Sec61b | 1.035171 | 0.196477 | 1.37E-07 | 9.62E-06 |
| Ccdc18 | 2.074867 | 0.393873 | 1.38E-07 | 9.64E-06 |
| Bub1b | 3.390031 | 0.645517 | 1.51E-07 | 1.05E-05 |
| Ccr1 | 2.261989 | 0.431032 | 1.54E-07 | 1.07E-05 |
| Tex15 | -1.51785 | 0.289709 | 1.61E-07 | 1.11E-05 |
| Krt6a | 4.235995 | 0.808546 | 1.61E-07 | 1.11E-05 |

| | | | | |
|---|---|---|---|---|
| Nsl1 | 3.267115 | 0.623806 | 1.63E-07 | 1.12E-05 |
| Rgs6 | -1.9126 | 0.365268 | 1.64E-07 | 1.12E-05 |
| Rgs11 | -1.15182 | 0.220139 | 1.67E-07 | 1.14E-05 |
| Ttk | 4.764238 | 0.9105 | 1.67E-07 | 1.14E-05 |
| Sprr2d | -3.46412 | 0.662965 | 1.74E-07 | 1.18E-05 |
| Apold1 | 2.124316 | 0.406547 | 1.74E-07 | 1.18E-05 |
| Ces1d | -2.33663 | 0.447672 | 1.79E-07 | 1.21E-05 |
| Gm16033 | 1.398282 | 0.267873 | 1.79E-07 | 1.21E-05 |
| Angpt4 | 3.031546 | 0.581369 | 1.84E-07 | 1.24E-05 |
| Brip1 | 3.394856 | 0.651376 | 1.87E-07 | 1.25E-05 |
| Inmt | -3.2848 | 0.630649 | 1.90E-07 | 1.27E-05 |
| Mgl2 | -2.21263 | 0.424962 | 1.92E-07 | 1.28E-05 |
| Dtl | 4.88615 | 0.938451 | 1.92E-07 | 1.28E-05 |
| Gm29491 | 2.485675 | 0.477549 | 1.94E-07 | 1.28E-05 |
| Itgb1 | 1.101016 | 0.211554 | 1.95E-07 | 1.28E-05 |
| Sprr2f | 2.956737 | 0.568779 | 2.01E-07 | 1.32E-05 |
| Hmcn2 | -2.86812 | 0.552044 | 2.04E-07 | 1.34E-05 |
| Plch1 | 2.921982 | 0.562535 | 2.05E-07 | 1.34E-05 |
| Trim59 | 2.242683 | 0.432434 | 2.15E-07 | 1.40E-05 |
| C330027C | 2.818541 | 0.543905 | 2.19E-07 | 1.42E-05 |
| Hspb6 | -1.35425 | 0.261423 | 2.22E-07 | 1.43E-05 |
| Ska3 | 3.288146 | 0.635129 | 2.25E-07 | 1.45E-05 |
| 4933413G | 3.027433 | 0.584885 | 2.27E-07 | 1.45E-05 |
| Fignl1 | 4.398488 | 0.851347 | 2.39E-07 | 1.52E-05 |
| Rab23 | 1.086292 | 0.210672 | 2.52E-07 | 1.60E-05 |
| Pnpla3 | -3.60152 | 0.699001 | 2.57E-07 | 1.62E-05 |
| Gnao1 | -1.70758 | 0.331505 | 2.59E-07 | 1.62E-05 |
| Gm25745 | 1.096307 | 0.212815 | 2.58E-07 | 1.62E-05 |
| Tmem171 | 6.15756 | 1.195308 | 2.58E-07 | 1.62E-05 |
| Trip13 | 3.431216 | 0.666741 | 2.66E-07 | 1.66E-05 |
| Hcn1 | -2.61434 | 0.508826 | 2.78E-07 | 1.73E-05 |
| Gm16336 | -1.57007 | 0.30564 | 2.79E-07 | 1.73E-05 |
| Filip1l | 1.539228 | 0.299965 | 2.88E-07 | 1.78E-05 |
| Exo1 | 5.098126 | 0.994184 | 2.93E-07 | 1.81E-05 |
| Fbln2 | 1.411423 | 0.275399 | 2.98E-07 | 1.83E-05 |
| AA986860 | -1.01851 | 0.198835 | 3.02E-07 | 1.85E-05 |
| Kcne3 | 3.150255 | 0.615598 | 3.10E-07 | 1.90E-05 |
| Gm4956 | -2.70423 | 0.528579 | 3.12E-07 | 1.90E-05 |
| Adamts1 | 1.329271 | 0.259843 | 3.13E-07 | 1.90E-05 |
| Tnfrsf12a | 3.248286 | 0.635045 | 3.14E-07 | 1.90E-05 |
| Espl1 | 3.39516 | 0.663921 | 3.16E-07 | 1.91E-05 |
| Trem1 | 6.437046 | 1.258869 | 3.16E-07 | 1.91E-05 |
| Lilrb4a | 1.124567 | 0.220236 | 3.29E-07 | 1.97E-05 |
| Arhgap11a | 2.505361 | 0.490626 | 3.28E-07 | 1.97E-05 |

| | | | | |
|---|---|---|---|---|
| Tnxb | -1.90852 | 0.374582 | 3.49E-07 | 2.08E-05 |
| Has2 | 2.450884 | 0.481054 | 3.49E-07 | 2.08E-05 |
| Ppp1r1a | -1.45313 | 0.285819 | 3.69E-07 | 2.19E-05 |
| BC030867 | 4.902659 | 0.964603 | 3.72E-07 | 2.20E-05 |
| Prr11 | 3.30246 | 0.649899 | 3.74E-07 | 2.21E-05 |
| Gm11947 | -2.28267 | 0.449303 | 3.77E-07 | 2.21E-05 |
| Peg10 | 1.851524 | 0.364559 | 3.80E-07 | 2.23E-05 |
| Top2a | 4.116617 | 0.810897 | 3.84E-07 | 2.25E-05 |
| Clspn | 4.890067 | 0.963707 | 3.89E-07 | 2.27E-05 |
| Camk4 | 1.418429 | 0.27969 | 3.95E-07 | 2.30E-05 |
| Fam111a | 2.720101 | 0.536499 | 3.98E-07 | 2.30E-05 |
| Hmga2 | 4.862457 | 0.95907 | 3.98E-07 | 2.30E-05 |
| Gm28661 | -1.48146 | 0.292258 | 4.00E-07 | 2.31E-05 |
| Stxbp6 | -1.3058 | 0.257792 | 4.08E-07 | 2.34E-05 |
| C5ar1 | 1.150303 | 0.227469 | 4.26E-07 | 2.43E-05 |
| Gm13461 | 1.92977 | 0.381601 | 4.26E-07 | 2.43E-05 |
| Stard4 | 1.558251 | 0.308384 | 4.35E-07 | 2.48E-05 |
| Gm15792 | 4.700961 | 0.930637 | 4.39E-07 | 2.49E-05 |
| Cenpn | 3.025275 | 0.599442 | 4.49E-07 | 2.54E-05 |
| Mei1 | -1.67927 | 0.332929 | 4.56E-07 | 2.56E-05 |
| Rtn4rl2 | 2.353625 | 0.466542 | 4.54E-07 | 2.56E-05 |
| Tnc | 2.250931 | 0.446748 | 4.69E-07 | 2.61E-05 |
| Hk2 | 2.012725 | 0.399748 | 4.78E-07 | 2.65E-05 |
| Manf | 1.433853 | 0.284807 | 4.79E-07 | 2.65E-05 |
| Mis18bp1 | 3.292293 | 0.654165 | 4.83E-07 | 2.66E-05 |
| Gm9800 | 1.331417 | 0.264798 | 4.96E-07 | 2.72E-05 |
| AV064505 | 6.075954 | 1.209744 | 5.10E-07 | 2.80E-05 |
| Polq | 3.471341 | 0.692018 | 5.27E-07 | 2.88E-05 |
| B4galnt1 | 1.095354 | 0.218674 | 5.47E-07 | 2.98E-05 |
| Apela | -5.98643 | 1.195938 | 5.57E-07 | 3.02E-05 |
| Steap1 | 2.283098 | 0.45625 | 5.61E-07 | 3.04E-05 |
| Ptgs2 | 4.708209 | 0.941407 | 5.70E-07 | 3.07E-05 |
| Npl | 1.782168 | 0.356459 | 5.74E-07 | 3.09E-05 |
| Pmch | 3.205749 | 0.641369 | 5.78E-07 | 3.10E-05 |
| Enc1 | 1.213575 | 0.242965 | 5.89E-07 | 3.15E-05 |
| Gja1 | 1.667727 | 0.334161 | 6.01E-07 | 3.21E-05 |
| Gm29216 | -1.92199 | 0.385329 | 6.10E-07 | 3.25E-05 |
| Mme | -1.46001 | 0.292785 | 6.14E-07 | 3.27E-05 |
| Hist1h2ao | 4.162836 | 0.835503 | 6.28E-07 | 3.33E-05 |
| Snhg6 | 1.711131 | 0.343511 | 6.32E-07 | 3.34E-05 |
| Kcnc4 | -5.6037 | 1.126912 | 6.61E-07 | 3.48E-05 |
| Gm28439 | -1.37091 | 0.275671 | 6.59E-07 | 3.48E-05 |
| Art3 | -1.58125 | 0.318164 | 6.70E-07 | 3.52E-05 |
| Tnfrsf23 | 1.431822 | 0.288179 | 6.75E-07 | 3.54E-05 |

| | | | | |
|---|---|---|---|---|
| Apobec2 | -2.77566 | 0.559229 | 6.93E-07 | 3.62E-05 |
| Slc29a1 | -1.24893 | 0.25163 | 6.93E-07 | 3.62E-05 |
| Plin4 | -1.63426 | 0.329467 | 7.04E-07 | 3.67E-05 |
| Pcnx2 | -1.54395 | 0.311307 | 7.07E-07 | 3.67E-05 |
| Dnaic1 | -1.80565 | 0.364236 | 7.15E-07 | 3.70E-05 |
| Armcx4 | 1.241302 | 0.250408 | 7.15E-07 | 3.70E-05 |
| Pde5a | -1.54081 | 0.310908 | 7.20E-07 | 3.71E-05 |
| Smc2 | 2.679695 | 0.541116 | 7.34E-07 | 3.77E-05 |
| Melk | 4.612987 | 0.932234 | 7.49E-07 | 3.84E-05 |
| Ppp1r3b | -1.20735 | 0.244319 | 7.74E-07 | 3.95E-05 |
| Cenpp | 2.12513 | 0.43003 | 7.74E-07 | 3.95E-05 |
| Fut2 | 2.698054 | 0.546411 | 7.90E-07 | 4.02E-05 |
| Efemp2 | 1.088367 | 0.220709 | 8.17E-07 | 4.12E-05 |
| Necab1 | -1.28191 | 0.260085 | 8.27E-07 | 4.16E-05 |
| 1810041L1 | -2.63454 | 0.53553 | 8.68E-07 | 4.35E-05 |
| Uchl1 | -1.31753 | 0.26787 | 8.72E-07 | 4.36E-05 |
| Cnnm1 | -1.0727 | 0.218301 | 8.93E-07 | 4.46E-05 |
| Tpbgl | -1.29454 | 0.263519 | 8.99E-07 | 4.46E-05 |
| Ptprn | 2.779472 | 0.56582 | 9.00E-07 | 4.46E-05 |
| Kif20b | 2.920548 | 0.594477 | 8.98E-07 | 4.46E-05 |
| Gm15845 | 5.972666 | 1.215959 | 9.02E-07 | 4.46E-05 |
| Birc5 | 4.779793 | 0.973391 | 9.09E-07 | 4.49E-05 |
| Sfrp5 | -1.95044 | 0.398282 | 9.72E-07 | 4.77E-05 |
| Kcnc1 | -3.99378 | 0.815979 | 9.86E-07 | 4.81E-05 |
| Gpr68 | 1.167366 | 0.238796 | 1.02E-06 | 4.94E-05 |
| Ube2c | 2.955903 | 0.605572 | 1.05E-06 | 5.11E-05 |
| Pde2a | -1.04982 | 0.215156 | 1.06E-06 | 5.13E-05 |
| Mybl2 | 3.331585 | 0.683153 | 1.08E-06 | 5.19E-05 |
| Tsku | -1.3135 | 0.269429 | 1.09E-06 | 5.22E-05 |
| Gm5648 | 6.806904 | 1.398477 | 1.13E-06 | 5.42E-05 |
| Hp | 2.366047 | 0.486169 | 1.13E-06 | 5.43E-05 |
| Ccnf | 3.123225 | 0.64232 | 1.16E-06 | 5.53E-05 |
| Abcg5 | -5.48614 | 1.128628 | 1.17E-06 | 5.55E-05 |
| Gkn3 | -2.07312 | 0.426553 | 1.17E-06 | 5.56E-05 |
| Tgfb2 | 1.580596 | 0.325341 | 1.18E-06 | 5.59E-05 |
| Aatk | -1.7192 | 0.354258 | 1.22E-06 | 5.73E-05 |
| Sgol2a | 3.160664 | 0.651419 | 1.22E-06 | 5.75E-05 |
| Slc16a3 | 1.334087 | 0.274984 | 1.23E-06 | 5.75E-05 |
| Gm28033 | 1.006576 | 0.207662 | 1.25E-06 | 5.85E-05 |
| Cdh24 | 2.049634 | 0.422866 | 1.25E-06 | 5.85E-05 |
| Chil3 | 5.506065 | 1.135838 | 1.25E-06 | 5.85E-05 |
| Tnik | -1.08224 | 0.223526 | 1.29E-06 | 5.97E-05 |
| Stil | 3.527971 | 0.728623 | 1.29E-06 | 5.97E-05 |
| Aurkb | 4.476448 | 0.924551 | 1.29E-06 | 5.97E-05 |

| | | | | |
|---|---|---|---|---|
| Fgl1 | 4.455885 | 0.92091 | 1.31E-06 | 6.05E-05 |
| Myadml2os | -1.97515 | 0.408333 | 1.32E-06 | 6.08E-05 |
| Aurka | 2.919722 | 0.604179 | 1.35E-06 | 6.21E-05 |
| Gm26904 | -1.31439 | 0.272178 | 1.37E-06 | 6.29E-05 |
| Incenp | 2.944377 | 0.610691 | 1.43E-06 | 6.53E-05 |
| 4732414G( | -2.8461 | 0.590436 | 1.43E-06 | 6.55E-05 |
| Tmed9 | 1.006292 | 0.208825 | 1.44E-06 | 6.56E-05 |
| Map1b | 1.024991 | 0.212712 | 1.45E-06 | 6.56E-05 |
| Ammecr1 | 1.18345 | 0.245614 | 1.45E-06 | 6.56E-05 |
| Apln | 1.588083 | 0.330047 | 1.50E-06 | 6.77E-05 |
| Ankle1 | 3.536206 | 0.735032 | 1.50E-06 | 6.78E-05 |
| Plat | 2.18458 | 0.454478 | 1.53E-06 | 6.91E-05 |
| Uhrf1 | 4.793821 | 0.997431 | 1.54E-06 | 6.92E-05 |
| Wnt10b | -2.92244 | 0.608143 | 1.54E-06 | 6.93E-05 |
| Plaur | 2.231944 | 0.464571 | 1.55E-06 | 6.96E-05 |
| Pglyrp1 | 2.03854 | 0.424739 | 1.59E-06 | 7.10E-05 |
| Timp4 | -1.77392 | 0.369755 | 1.61E-06 | 7.14E-05 |
| Mall | 2.374256 | 0.494881 | 1.61E-06 | 7.14E-05 |
| Lrp8 | 3.169602 | 0.66078 | 1.61E-06 | 7.15E-05 |
| Fam132b | 3.107988 | 0.648014 | 1.62E-06 | 7.16E-05 |
| Kcna1 | -2.40571 | 0.501737 | 1.63E-06 | 7.20E-05 |
| Crhbp | -2.62402 | 0.547566 | 1.65E-06 | 7.27E-05 |
| Spag5 | 4.180205 | 0.872346 | 1.65E-06 | 7.27E-05 |
| Gm12498 | -1.31015 | 0.274133 | 1.76E-06 | 7.73E-05 |
| Sgk1 | 1.044545 | 0.21863 | 1.77E-06 | 7.76E-05 |
| 4930427A( | 3.058444 | 0.640248 | 1.78E-06 | 7.76E-05 |
| Htr2a | 2.415125 | 0.506675 | 1.87E-06 | 8.14E-05 |
| 1700056E2 | 1.171905 | 0.245898 | 1.88E-06 | 8.15E-05 |
| Kntc1 | 4.477424 | 0.939746 | 1.89E-06 | 8.19E-05 |
| Foxm1 | 3.025323 | 0.635091 | 1.90E-06 | 8.21E-05 |
| Hbegf | 2.083658 | 0.437516 | 1.91E-06 | 8.24E-05 |
| Rrm2 | 3.882349 | 0.815259 | 1.92E-06 | 8.24E-05 |
| P4ha3 | 2.591987 | 0.544828 | 1.96E-06 | 8.40E-05 |
| Mki67 | 4.305916 | 0.905273 | 1.97E-06 | 8.43E-05 |
| Fam3b | 2.08776 | 0.439137 | 1.99E-06 | 8.51E-05 |
| Gm8319 | 1.276602 | 0.2689 | 2.06E-06 | 8.76E-05 |
| Ube2ql1 | -1.93497 | 0.407778 | 2.08E-06 | 8.82E-05 |
| Lbh | 1.690817 | 0.356305 | 2.08E-06 | 8.82E-05 |
| Sgol1 | 2.886536 | 0.608381 | 2.09E-06 | 8.82E-05 |
| 2810408I1 | 3.347026 | 0.705832 | 2.12E-06 | 8.92E-05 |
| Kif15 | 4.305983 | 0.908147 | 2.12E-06 | 8.93E-05 |
| Adam19 | 1.40766 | 0.297121 | 2.16E-06 | 9.06E-05 |
| Tgm3 | -2.17321 | 0.458787 | 2.17E-06 | 9.08E-05 |
| S100a9 | 4.190863 | 0.885983 | 2.24E-06 | 9.37E-05 |

| | | | | |
|---|---|---|---|---|
| Orc1 | 3.708936 | 0.784352 | 2.26E-06 | 9.41E-05 |
| Has1 | 5.731096 | 1.212037 | 2.26E-06 | 9.41E-05 |
| Pcolce2 | -1.82653 | 0.386513 | 2.29E-06 | 9.48E-05 |
| Avpr1a | -1.2256 | 0.259312 | 2.29E-06 | 9.48E-05 |
| Tcf19 | 2.974851 | 0.629438 | 2.29E-06 | 9.48E-05 |
| Frmpd4 | -2.32539 | 0.492264 | 2.31E-06 | 9.54E-05 |
| Fbxo16 | -1.06165 | 0.224768 | 2.32E-06 | 9.55E-05 |
| Slc6a2 | 1.932588 | 0.409248 | 2.33E-06 | 9.58E-05 |
| Trib1 | 1.815818 | 0.384793 | 2.37E-06 | 9.66E-05 |
| A830012C | 2.961238 | 0.627538 | 2.37E-06 | 9.66E-05 |
| Efcab11 | 3.847858 | 0.815335 | 2.37E-06 | 9.66E-05 |
| Ltbp2 | 3.909413 | 0.828357 | 2.36E-06 | 9.66E-05 |
| 1700061G | -1.63874 | 0.347474 | 2.40E-06 | 9.77E-05 |
| Gm11992 | -1.37597 | 0.291977 | 2.45E-06 | 9.92E-05 |
| Kif18b | 4.042828 | 0.858018 | 2.46E-06 | 9.94E-05 |
| Gpr35 | 1.414483 | 0.300266 | 2.47E-06 | 9.98E-05 |
| Rgma | -1.39853 | 0.297007 | 2.49E-06 | 0.000101 |
| Akap2 | 2.157319 | 0.458246 | 2.50E-06 | 0.000101 |
| Gata4 | -1.49847 | 0.318441 | 2.53E-06 | 0.000102 |
| Dsg3 | 2.940485 | 0.625037 | 2.54E-06 | 0.000102 |
| Cbx7 | -1.13265 | 0.2408 | 2.55E-06 | 0.000102 |
| Pakap | 2.142537 | 0.455872 | 2.60E-06 | 0.000104 |
| Pappa | 1.189932 | 0.253307 | 2.63E-06 | 0.000105 |
| Pim1 | 1.506095 | 0.320593 | 2.63E-06 | 0.000105 |
| Raver2 | -1.41687 | 0.301717 | 2.65E-06 | 0.000105 |
| Plk1 | 2.749978 | 0.586597 | 2.76E-06 | 0.000109 |
| Srl | -1.18401 | 0.252728 | 2.80E-06 | 0.000111 |
| Gmnn | 2.425164 | 0.51784 | 2.82E-06 | 0.000111 |
| Gm28294 | 3.97704 | 0.849242 | 2.83E-06 | 0.000111 |
| Dclk1 | 1.290401 | 0.275713 | 2.87E-06 | 0.000113 |
| Suv39h1 | 1.463752 | 0.313345 | 2.99E-06 | 0.000116 |
| Gm43305 | 1.733896 | 0.371184 | 2.99E-06 | 0.000116 |
| Grem1 | 5.489683 | 1.175038 | 2.98E-06 | 0.000116 |
| Mmp14 | 1.411498 | 0.302286 | 3.02E-06 | 0.000117 |
| Bmp8a | -1.4549 | 0.311749 | 3.06E-06 | 0.000118 |
| Kbtbd6 | 3.059451 | 0.656185 | 3.12E-06 | 0.00012 |
| Ereg | 2.754588 | 0.591691 | 3.23E-06 | 0.000124 |
| Dusp10 | 1.861271 | 0.400153 | 3.30E-06 | 0.000126 |
| Pacsin1 | -1.64627 | 0.354467 | 3.41E-06 | 0.00013 |
| Kifc5b | 2.53813 | 0.546649 | 3.43E-06 | 0.00013 |
| 2310002F | -1.3672 | 0.294504 | 3.44E-06 | 0.00013 |
| Cthrc1 | 3.757984 | 0.809923 | 3.49E-06 | 0.000132 |
| Cks1b | 2.327781 | 0.502559 | 3.62E-06 | 0.000137 |
| Dock3 | -1.27262 | 0.275079 | 3.72E-06 | 0.00014 |

| | | | | |
|---|---|---|---|---|
| Arhgap19 | 1.991043 | 0.430974 | 3.84E-06 | 0.000144 |
| Plk4 | 2.078751 | 0.449995 | 3.85E-06 | 0.000144 |
| Gm10143 | 2.870199 | 0.622443 | 4.00E-06 | 0.000149 |
| Cdca5 | 4.153737 | 0.901659 | 4.09E-06 | 0.000152 |
| Zfp536 | -1.71173 | 0.372067 | 4.21E-06 | 0.000156 |
| Lsm5 | 1.157974 | 0.251691 | 4.21E-06 | 0.000156 |
| Tnfsf18 | 6.052867 | 1.315736 | 4.22E-06 | 0.000156 |
| Npy1r | -2.36264 | 0.51365 | 4.23E-06 | 0.000156 |
| Il4ra | 1.877484 | 0.408199 | 4.24E-06 | 0.000156 |
| Gins1 | 2.715071 | 0.590703 | 4.30E-06 | 0.000158 |
| Sec1 | 3.05468 | 0.664533 | 4.29E-06 | 0.000158 |
| 1700023F( | -2.35365 | 0.51306 | 4.49E-06 | 0.000163 |
| Mapt | -1.99556 | 0.435041 | 4.50E-06 | 0.000163 |
| Msn | 1.716899 | 0.374529 | 4.56E-06 | 0.000165 |
| 4930558J1 | 6.273814 | 1.368566 | 4.56E-06 | 0.000165 |
| Gm9767 | 2.236714 | 0.489153 | 4.82E-06 | 0.000173 |
| Ticrr | 4.557143 | 0.996648 | 4.82E-06 | 0.000173 |
| Eya4 | -1.09205 | 0.238988 | 4.89E-06 | 0.000175 |
| Tagln2 | 1.595469 | 0.349223 | 4.91E-06 | 0.000176 |
| Nr4a1 | 2.018345 | 0.441801 | 4.91E-06 | 0.000176 |
| Syt12 | 3.020901 | 0.66167 | 4.98E-06 | 0.000178 |
| Kif23 | 2.636852 | 0.577651 | 5.00E-06 | 0.000178 |
| Nrg2 | -1.22738 | 0.268967 | 5.04E-06 | 0.000179 |
| Ncaph | 2.8912 | 0.633678 | 5.05E-06 | 0.000179 |
| Gm44397 | 2.138852 | 0.468956 | 5.09E-06 | 0.00018 |
| Sirpb1c | 2.330768 | 0.511419 | 5.18E-06 | 0.000182 |
| Bglap2 | -1.64768 | 0.361587 | 5.19E-06 | 0.000183 |
| Dusp6 | 1.861523 | 0.408708 | 5.25E-06 | 0.000184 |
| Add2 | -1.8862 | 0.414291 | 5.29E-06 | 0.000185 |
| Chaf1a | 2.657036 | 0.583688 | 5.31E-06 | 0.000186 |
| Cda | 2.444545 | 0.537122 | 5.33E-06 | 0.000186 |
| Fen1 | 2.830651 | 0.6224 | 5.42E-06 | 0.000189 |
| Hells | 4.513463 | 0.992874 | 5.47E-06 | 0.00019 |
| Ank1 | -1.27038 | 0.279484 | 5.48E-06 | 0.00019 |
| Fbxw27 | -1.44667 | 0.318387 | 5.53E-06 | 0.000191 |
| Nxpe5 | 1.484682 | 0.32722 | 5.70E-06 | 0.000197 |
| Gm14221 | 1.509309 | 0.332824 | 5.76E-06 | 0.000198 |
| Penk | 2.857233 | 0.63227 | 6.21E-06 | 0.000213 |
| Pdia6 | 1.117057 | 0.247299 | 6.27E-06 | 0.000215 |
| Timeless | 2.597384 | 0.57528 | 6.33E-06 | 0.000216 |
| Reg3b | 7.228678 | 1.60149 | 6.37E-06 | 0.000217 |
| Kif4 | 2.37196 | 0.525662 | 6.41E-06 | 0.000218 |
| Spock2 | -1.65319 | 0.366504 | 6.46E-06 | 0.00022 |
| Adcy9 | -1.10478 | 0.245068 | 6.54E-06 | 0.000222 |

| | | | | |
|---|---|---|---|---|
| Cebpd | 1.912047 | 0.424633 | 6.71E-06 | 0.000227 |
| Gm26674 | -1.0106 | 0.224672 | 6.86E-06 | 0.000231 |
| Aldh1a2 | 1.315307 | 0.292398 | 6.85E-06 | 0.000231 |
| Igfbp5 | -1.2124 | 0.269629 | 6.91E-06 | 0.000232 |
| Tppp3 | -1.58097 | 0.35175 | 6.97E-06 | 0.000234 |
| Dusp5 | 2.086147 | 0.464898 | 7.21E-06 | 0.000241 |
| Il18rap | 2.648432 | 0.590809 | 7.37E-06 | 0.000246 |
| Mcm10 | 4.518609 | 1.007957 | 7.36E-06 | 0.000246 |
| Mastl | 2.678492 | 0.597849 | 7.46E-06 | 0.000248 |
| Atp2b2 | -2.86432 | 0.639488 | 7.50E-06 | 0.000249 |
| Kif20a | 2.08285 | 0.465567 | 7.68E-06 | 0.000255 |
| Wdhd1 | 3.021414 | 0.676049 | 7.85E-06 | 0.00026 |
| B3gnt3 | 1.647081 | 0.368796 | 7.97E-06 | 0.000263 |
| Bex4 | -1.54841 | 0.347217 | 8.22E-06 | 0.000271 |
| D430036J1 | 1.983877 | 0.444909 | 8.23E-06 | 0.000271 |
| 8030451A( | 1.684386 | 0.377868 | 8.29E-06 | 0.000273 |
| Gm35507 | 6.374825 | 1.431121 | 8.41E-06 | 0.000276 |
| Ect2 | 3.677035 | 0.825538 | 8.42E-06 | 0.000276 |
| Msr1 | 1.600412 | 0.359417 | 8.48E-06 | 0.000278 |
| 4930458D( | -1.27243 | 0.285793 | 8.50E-06 | 0.000278 |
| Asb5 | 1.845392 | 0.414837 | 8.65E-06 | 0.000282 |
| Psd2 | -2.17327 | 0.488763 | 8.73E-06 | 0.000284 |
| Gm12868 | 1.044115 | 0.234901 | 8.79E-06 | 0.000285 |
| Gm17315 | -1.45672 | 0.328033 | 8.96E-06 | 0.00029 |
| Hist1h4i | 1.432339 | 0.322553 | 8.97E-06 | 0.00029 |
| Atf3 | 2.729482 | 0.614582 | 8.95E-06 | 0.00029 |
| Gm5150 | 4.944495 | 1.113653 | 9.00E-06 | 0.00029 |
| Gm9887 | 1.383533 | 0.311669 | 9.03E-06 | 0.000291 |
| Pstpip1 | 1.17889 | 0.265612 | 9.06E-06 | 0.000291 |
| Scube2 | -1.76177 | 0.397864 | 9.51E-06 | 0.000304 |
| Lsm7 | 1.080446 | 0.244032 | 9.53E-06 | 0.000304 |
| Tgfbi | 1.067851 | 0.241737 | 9.99E-06 | 0.000317 |
| Lmnb1 | 2.218147 | 0.502188 | 1.00E-05 | 0.000317 |
| Gm14820 | 1.635078 | 0.370495 | 1.02E-05 | 0.000322 |
| Cks2 | 2.059666 | 0.466699 | 1.02E-05 | 0.000322 |
| Rassf1 | 1.485474 | 0.336789 | 1.03E-05 | 0.000325 |
| Gm44123 | -1.24195 | 0.281627 | 1.03E-05 | 0.000325 |
| Nppb | 7.166448 | 1.627877 | 1.07E-05 | 0.000336 |
| Zwilch | 2.705477 | 0.614615 | 1.07E-05 | 0.000336 |
| Synpo | 1.044391 | 0.237276 | 1.07E-05 | 0.000336 |
| Iqgap3 | 3.57596 | 0.8128 | 1.08E-05 | 0.000339 |
| Cldn2 | 3.43604 | 0.781235 | 1.09E-05 | 0.00034 |
| Bard1 | 2.774476 | 0.63124 | 1.11E-05 | 0.000344 |
| Asf1b | 4.34849 | 0.98987 | 1.12E-05 | 0.000346 |

| | | | | |
|---|---|---|---|---|
| Gm14317 | -2.11387 | 0.481497 | 1.13E-05 | 0.00035 |
| Gm10925 | -1.07751 | 0.245608 | 1.15E-05 | 0.000354 |
| Ercc6l | 4.202584 | 0.958269 | 1.16E-05 | 0.000356 |
| Gjb4 | 2.692242 | 0.61403 | 1.16E-05 | 0.000357 |
| Fcgr4 | 1.995701 | 0.45524 | 1.17E-05 | 0.000358 |
| Gm17080 | -1.19678 | 0.273259 | 1.19E-05 | 0.000363 |
| Smc4 | 1.633607 | 0.372987 | 1.19E-05 | 0.000363 |
| Dbf4 | 2.491932 | 0.568907 | 1.19E-05 | 0.000363 |
| Snhg4 | 1.750891 | 0.399896 | 1.20E-05 | 0.000364 |
| E2f8 | 4.697223 | 1.072798 | 1.20E-05 | 0.000364 |
| Kif11 | 3.900507 | 0.891121 | 1.20E-05 | 0.000366 |
| Gpr1 | -1.76741 | 0.404044 | 1.22E-05 | 0.00037 |
| Syt2 | -1.85165 | 0.424109 | 1.27E-05 | 0.000384 |
| Tlr13 | 1.43931 | 0.330121 | 1.30E-05 | 0.000393 |
| Aldh2 | -1.02407 | 0.234923 | 1.31E-05 | 0.000394 |
| Rgs7bp | -1.57603 | 0.361932 | 1.33E-05 | 0.000401 |
| Scnn1g | -1.19301 | 0.274024 | 1.34E-05 | 0.000401 |
| Gm43003 | 6.157146 | 1.41471 | 1.35E-05 | 0.000404 |
| Gm19658 | 3.981999 | 0.91519 | 1.36E-05 | 0.000405 |
| Ncapg | 2.710872 | 0.623405 | 1.37E-05 | 0.000408 |
| Tcf23 | -2.07232 | 0.477047 | 1.40E-05 | 0.000416 |
| Gm6091 | 4.234842 | 0.975133 | 1.41E-05 | 0.000417 |
| Vcan | 1.270903 | 0.292666 | 1.41E-05 | 0.000417 |
| Rad51c | 1.299033 | 0.299232 | 1.42E-05 | 0.000419 |
| Ipcef1 | -1.57173 | 0.362199 | 1.43E-05 | 0.000421 |
| Mir8101 | -1.43828 | 0.331541 | 1.44E-05 | 0.000423 |
| Gm16175 | 2.354499 | 0.54292 | 1.45E-05 | 0.000425 |
| Sorbs2os | -1.21796 | 0.280895 | 1.45E-05 | 0.000426 |
| Spry2 | 1.488277 | 0.343585 | 1.48E-05 | 0.000433 |
| Cdc45 | 2.7063 | 0.624743 | 1.48E-05 | 0.000433 |
| Tfpi2 | 1.336324 | 0.308621 | 1.49E-05 | 0.000436 |
| Notum | -1.82179 | 0.420774 | 1.49E-05 | 0.000436 |
| Htr4 | -1.08517 | 0.250656 | 1.50E-05 | 0.000436 |
| Gm15816 | -1.20273 | 0.27788 | 1.50E-05 | 0.000437 |
| Gm10425 | 1.017182 | 0.23502 | 1.50E-05 | 0.000437 |
| Ccl12 | 2.020383 | 0.466895 | 1.51E-05 | 0.000438 |
| Gm2788 | 4.910077 | 1.135401 | 1.53E-05 | 0.000443 |
| Chpt1 | -1.02012 | 0.23607 | 1.55E-05 | 0.000449 |
| Gm21596 | 1.218824 | 0.28218 | 1.57E-05 | 0.000452 |
| Cdc7 | 2.559295 | 0.592588 | 1.57E-05 | 0.000452 |
| 3425401B1 | -1.84503 | 0.427333 | 1.58E-05 | 0.000453 |
| Pm20d1 | -1.01716 | 0.235594 | 1.58E-05 | 0.000453 |
| Rhox5 | 1.185347 | 0.274765 | 1.60E-05 | 0.000458 |
| Ltb4r1 | 2.770846 | 0.642449 | 1.61E-05 | 0.000459 |

| | | | | |
|---|---|---|---|---|
| Nfkbiz | 1.509252 | 0.350126 | 1.63E-05 | 0.000464 |
| Rrad | 1.043289 | 0.242117 | 1.64E-05 | 0.000467 |
| Ppm1j | 1.573256 | 0.365412 | 1.67E-05 | 0.000474 |
| Foxo6 | -1.71216 | 0.397812 | 1.68E-05 | 0.000475 |
| 9530052E( | -1.47521 | 0.342747 | 1.68E-05 | 0.000475 |
| Troap | 3.020083 | 0.701694 | 1.68E-05 | 0.000475 |
| F3 | 1.770386 | 0.411674 | 1.70E-05 | 0.000482 |
| Tnfrsf9 | 1.622759 | 0.377769 | 1.74E-05 | 0.000492 |
| Gm26673 | -2.06308 | 0.481525 | 1.83E-05 | 0.000515 |
| Nfil3 | 1.930273 | 0.451016 | 1.87E-05 | 0.000524 |
| Creld2 | 1.1691 | 0.273208 | 1.88E-05 | 0.000525 |
| Nmur2 | -1.61161 | 0.37713 | 1.93E-05 | 0.000536 |
| Chtf18 | 2.880575 | 0.674282 | 1.94E-05 | 0.000539 |
| Lig1 | 2.411158 | 0.564467 | 1.94E-05 | 0.000539 |
| Apitd1 | 2.435127 | 0.570174 | 1.95E-05 | 0.000541 |
| Cyp26b1 | -1.44459 | 0.338447 | 1.97E-05 | 0.000546 |
| Fgf23 | 6.22694 | 1.46057 | 2.01E-05 | 0.000557 |
| Runx2os1 | 1.593648 | 0.373912 | 2.03E-05 | 0.000559 |
| Traip | 2.980195 | 0.699733 | 2.05E-05 | 0.000565 |
| Sox2ot | -1.38995 | 0.326403 | 2.06E-05 | 0.000566 |
| Serpinb6b | 1.034681 | 0.243006 | 2.06E-05 | 0.000566 |
| Gm7665 | 1.053488 | 0.247423 | 2.06E-05 | 0.000566 |
| Snrpe | 1.089837 | 0.255989 | 2.07E-05 | 0.000567 |
| Slc25a15 | 1.38672 | 0.325779 | 2.08E-05 | 0.000568 |
| Tm4sf1 | 1.322618 | 0.310843 | 2.09E-05 | 0.000571 |
| Gm10605 | -1.49568 | 0.351558 | 2.10E-05 | 0.000572 |
| Cxcl1 | 3.053917 | 0.718509 | 2.13E-05 | 0.00058 |
| Ppp1r12b | -1.29114 | 0.303924 | 2.15E-05 | 0.000582 |
| Scnn1b | -1.23702 | 0.291181 | 2.15E-05 | 0.000582 |
| Sprr2g | 2.255575 | 0.53107 | 2.16E-05 | 0.000584 |
| Pole | 3.87749 | 0.913164 | 2.17E-05 | 0.000586 |
| Rora | -1.06615 | 0.251264 | 2.20E-05 | 0.000592 |
| Cd177 | 6.370746 | 1.501415 | 2.20E-05 | 0.000592 |
| Gm19705 | 1.730178 | 0.40792 | 2.22E-05 | 0.000594 |
| Slc16a10 | -1.57582 | 0.371673 | 2.24E-05 | 0.000597 |
| Rad51ap1 | 3.794407 | 0.895281 | 2.25E-05 | 0.0006 |
| Hspa5 | 1.471687 | 0.347562 | 2.29E-05 | 0.00061 |
| Cep85 | 1.324225 | 0.313154 | 2.35E-05 | 0.000622 |
| Nasp | 1.811231 | 0.429002 | 2.42E-05 | 0.000638 |
| Cxcl3 | 8.137958 | 1.927577 | 2.42E-05 | 0.000638 |
| Plod2 | 1.583453 | 0.375403 | 2.46E-05 | 0.000648 |
| Kcnn2 | -2.23783 | 0.531098 | 2.51E-05 | 0.00066 |
| E130012A1 | 2.122837 | 0.50428 | 2.56E-05 | 0.00067 |
| Klrb1a | -5.30411 | 1.260672 | 2.58E-05 | 0.000675 |

| | | | | |
|---|---|---|---|---|
| Pthlh | -1.8955 | 0.450611 | 2.59E-05 | 0.000677 |
| Tac2 | 6.473582 | 1.539133 | 2.60E-05 | 0.000678 |
| Hmgb2 | 2.269483 | 0.539899 | 2.63E-05 | 0.000684 |
| Mybph | -1.08558 | 0.258353 | 2.65E-05 | 0.000686 |
| Mad2l1 | 2.354282 | 0.560291 | 2.65E-05 | 0.000686 |
| Gareml | 2.730417 | 0.651189 | 2.75E-05 | 0.000711 |
| Gm44486 | 4.323654 | 1.03117 | 2.75E-05 | 0.000711 |
| Cdh19 | -1.39088 | 0.331763 | 2.76E-05 | 0.000712 |
| Hif1a | 1.78608 | 0.426409 | 2.81E-05 | 0.000722 |
| Pgf | 1.23158 | 0.29419 | 2.83E-05 | 0.000728 |
| Mcm6 | 2.525173 | 0.603226 | 2.84E-05 | 0.000728 |
| P3h1 | 1.074072 | 0.256758 | 2.87E-05 | 0.000736 |
| Gm6793 | 1.48272 | 0.354504 | 2.88E-05 | 0.000737 |
| Prg4 | 1.267253 | 0.30314 | 2.91E-05 | 0.000743 |
| Mansc4 | -2.31309 | 0.553561 | 2.93E-05 | 0.000749 |
| Arid3c | -1.64034 | 0.392914 | 2.98E-05 | 0.000756 |
| Dhfr | 2.537551 | 0.60784 | 2.98E-05 | 0.000756 |
| Gm12960 | 1.47889 | 0.354292 | 2.99E-05 | 0.000757 |
| Kcnq4 | -1.32025 | 0.316354 | 3.00E-05 | 0.000759 |
| Arhgef26 | -1.3888 | 0.332943 | 3.03E-05 | 0.000765 |
| Cfap157 | 1.530353 | 0.366959 | 3.04E-05 | 0.000767 |
| Mthfd2 | 2.622531 | 0.628986 | 3.05E-05 | 0.00077 |
| Fmo2 | -1.78104 | 0.427324 | 3.07E-05 | 0.000773 |
| Cdkn2d | 1.247745 | 0.299748 | 3.15E-05 | 0.00079 |
| Serpinb2 | 5.262832 | 1.264668 | 3.16E-05 | 0.000793 |
| Gm20490 | -1.67441 | 0.40285 | 3.23E-05 | 0.000809 |
| Foxo6os | -3.15815 | 0.760635 | 3.30E-05 | 0.000822 |
| Wfdc17 | 1.334253 | 0.321372 | 3.30E-05 | 0.000822 |
| Lrr1 | 5.059418 | 1.218851 | 3.31E-05 | 0.000822 |
| Gm6377 | 2.291957 | 0.552194 | 3.32E-05 | 0.000822 |
| Plekhb1 | -1.10082 | 0.26531 | 3.34E-05 | 0.000825 |
| Lif | 2.349379 | 0.566233 | 3.34E-05 | 0.000825 |
| Gm26885 | 1.55592 | 0.375152 | 3.36E-05 | 0.00083 |
| Edn3 | -2.9964 | 0.72272 | 3.38E-05 | 0.000833 |
| Hist1h2ag | 4.111941 | 0.991739 | 3.38E-05 | 0.000833 |
| Svopl | -1.9204 | 0.463373 | 3.41E-05 | 0.000838 |
| Fam198b | 1.375333 | 0.33215 | 3.46E-05 | 0.000851 |
| Gm16576 | -1.32911 | 0.321025 | 3.47E-05 | 0.000852 |
| Myef2 | 1.156971 | 0.27948 | 3.48E-05 | 0.000853 |
| Gm28221 | -1.44096 | 0.348107 | 3.48E-05 | 0.000853 |
| Mmp3 | 1.976462 | 0.47757 | 3.49E-05 | 0.000854 |
| Cacna1b | -1.65337 | 0.399676 | 3.52E-05 | 0.000859 |
| Kcnj14 | 3.37099 | 0.81523 | 3.55E-05 | 0.000865 |
| Pde9a | -1.4525 | 0.351313 | 3.56E-05 | 0.000866 |

| | | | | |
|---|---|---|---|---|
| Gm36401 | 2.85394 | 0.690361 | 3.57E-05 | 0.000867 |
| Glipr2 | 1.088473 | 0.263585 | 3.64E-05 | 0.000881 |
| Mms22l | 2.39045 | 0.579137 | 3.67E-05 | 0.000888 |
| Igf1 | 1.221625 | 0.296022 | 3.68E-05 | 0.000889 |
| Kcnj5 | -2.72415 | 0.660652 | 3.73E-05 | 0.0009 |
| Gm16548 | 1.902519 | 0.461377 | 3.73E-05 | 0.0009 |
| Tnxa | -2.52891 | 0.613553 | 3.76E-05 | 0.000905 |
| Sync | 1.261595 | 0.306169 | 3.78E-05 | 0.000907 |
| Ppp1r18os | 1.690847 | 0.41035 | 3.78E-05 | 0.000907 |
| Bcl3 | 2.119981 | 0.514514 | 3.78E-05 | 0.000907 |
| Ddias | 1.957968 | 0.475377 | 3.81E-05 | 0.000913 |
| Gm10222 | -1.23184 | 0.29913 | 3.82E-05 | 0.000914 |
| Gm35801 | -1.21905 | 0.296026 | 3.82E-05 | 0.000914 |
| Klhl14 | -1.03769 | 0.252082 | 3.85E-05 | 0.000919 |
| Cep85l | -1.12421 | 0.273424 | 3.93E-05 | 0.000935 |
| Ung | 3.581827 | 0.871444 | 3.95E-05 | 0.000939 |
| Retnlg | 5.729482 | 1.394073 | 3.96E-05 | 0.00094 |
| Fcgr1 | 1.302081 | 0.316983 | 4.00E-05 | 0.000947 |
| Rxrg | -2.62327 | 0.638733 | 4.01E-05 | 0.000949 |
| Il33 | 1.706942 | 0.415699 | 4.02E-05 | 0.000951 |
| E2f2 | 2.143251 | 0.522135 | 4.05E-05 | 0.000956 |
| Sh2d5 | 3.927202 | 0.957893 | 4.13E-05 | 0.000974 |
| Anp32b | 1.124768 | 0.274377 | 4.14E-05 | 0.000975 |
| Sult1a1 | -1.30781 | 0.319104 | 4.16E-05 | 0.000978 |

Supplemental Table 3. d16 SCI + CMC vs Sham DEGs

| GeneID | Log2 fc | SE | p-value | FDR p-value |
|---|---|---|---|---|
| Pcsk5 | 2.202166 | 0.16158 | 2.70E-42 | 5.48E-38 |
| Cilp | 3.546124 | 0.288539 | 1.03E-34 | 1.04E-30 |
| Cthrc1 | 4.28179 | 0.378844 | 1.28E-29 | 8.67E-26 |
| Eln | 3.415173 | 0.320224 | 1.48E-26 | 7.55E-23 |
| Gm43278 | 3.535602 | 0.369789 | 1.16E-21 | 4.74E-18 |
| Fstl1 | 1.506855 | 0.160422 | 5.83E-21 | 1.98E-17 |
| Wisp2 | 4.00923 | 0.429774 | 1.07E-20 | 3.11E-17 |
| Pla1a | 1.843253 | 0.200926 | 4.57E-20 | 1.16E-16 |
| Crlf1 | 5.758974 | 0.633614 | 9.99E-20 | 1.85E-16 |
| Sfrp1 | 2.676118 | 0.293889 | 8.56E-20 | 1.85E-16 |
| Loxl2 | 2.515645 | 0.276731 | 9.85E-20 | 1.85E-16 |
| Lox | 3.071954 | 0.340644 | 1.91E-19 | 3.24E-16 |
| Frzb | 2.727875 | 0.30329 | 2.38E-19 | 3.72E-16 |
| Mfap4 | 2.031281 | 0.226108 | 2.62E-19 | 3.80E-16 |
| Mchr1 | 1.815355 | 0.207242 | 1.96E-18 | 2.66E-15 |
| P4ha3 | 2.981772 | 0.358191 | 8.47E-17 | 1.08E-13 |
| Bmper | 3.648053 | 0.439314 | 1.01E-16 | 1.20E-13 |
| Slc6a2 | 3.677331 | 0.445699 | 1.57E-16 | 1.78E-13 |
| Bgn | 1.361457 | 0.169466 | 9.45E-16 | 1.01E-12 |
| Cpa3 | 3.388424 | 0.42344 | 1.22E-15 | 1.22E-12 |
| Loxl1 | 1.593005 | 0.199166 | 1.26E-15 | 1.22E-12 |
| Wisp1 | 2.746395 | 0.347118 | 2.53E-15 | 2.34E-12 |
| Rxfp1 | 4.159719 | 0.526533 | 2.78E-15 | 2.46E-12 |
| Itih4 | 2.20447 | 0.28197 | 5.36E-15 | 4.55E-12 |
| Csgalnact1 | 1.895493 | 0.243486 | 6.98E-15 | 5.68E-12 |
| Mfap5 | 1.921479 | 0.247036 | 7.36E-15 | 5.76E-12 |
| Lgi2 | 2.05935 | 0.265263 | 8.27E-15 | 6.23E-12 |
| Chrdl2 | 3.316682 | 0.429188 | 1.09E-14 | 7.95E-12 |
| Ltbp2 | 4.22736 | 0.557586 | 3.41E-14 | 2.40E-11 |
| Atcayos | -2.40528 | 0.320385 | 6.03E-14 | 4.09E-11 |
| Trdn | -3.0469 | 0.412157 | 1.44E-13 | 9.45E-11 |
| Cma1 | 3.128194 | 0.427818 | 2.63E-13 | 1.64E-10 |
| Mmp3 | 2.942072 | 0.402462 | 2.67E-13 | 1.64E-10 |
| Serpine2 | 1.675169 | 0.231138 | 4.25E-13 | 2.54E-10 |
| Ptprz1 | 2.696327 | 0.385571 | 2.69E-12 | 1.56E-09 |
| Grem1 | 9.137743 | 1.31553 | 3.76E-12 | 2.12E-09 |
| Col8a1 | 1.397042 | 0.201821 | 4.45E-12 | 2.44E-09 |
| Mir6950 | 2.511448 | 0.365811 | 6.63E-12 | 3.46E-09 |
| Ccr5 | 2.613707 | 0.388442 | 1.71E-11 | 8.71E-09 |
| Hhipl1 | 2.799558 | 0.416411 | 1.78E-11 | 8.83E-09 |
| 4930478K1 | 3.742138 | 0.55867 | 2.11E-11 | 1.02E-08 |
| 1500009L1 | 2.641693 | 0.396327 | 2.64E-11 | 1.25E-08 |

| | | | | |
|---|---|---|---|---|
| Fst | 2.507363 | 0.376982 | 2.91E-11 | 1.34E-08 |
| Col5a2 | 1.459915 | 0.21959 | 2.96E-11 | 1.34E-08 |
| Chrd | 1.263429 | 0.192008 | 4.70E-11 | 2.08E-08 |
| Adamts2 | 1.388254 | 0.211302 | 5.03E-11 | 2.18E-08 |
| Fbn1 | 1.793447 | 0.277073 | 9.62E-11 | 4.08E-08 |
| Adam12 | 1.816582 | 0.28665 | 2.34E-10 | 9.71E-08 |
| Fcgr4 | 2.859708 | 0.452518 | 2.62E-10 | 1.07E-07 |
| Tpsb2 | 2.273319 | 0.36027 | 2.79E-10 | 1.11E-07 |
| Lilr4b | 2.037013 | 0.324629 | 3.50E-10 | 1.37E-07 |
| Ccr2 | 1.767245 | 0.282087 | 3.73E-10 | 1.43E-07 |
| Runx3 | 1.713068 | 0.27422 | 4.18E-10 | 1.55E-07 |
| Vwa1 | 1.514363 | 0.243351 | 4.88E-10 | 1.77E-07 |
| Ms4a6d | 2.289191 | 0.368275 | 5.10E-10 | 1.82E-07 |
| Mcpt4 | 2.139056 | 0.344346 | 5.23E-10 | 1.84E-07 |
| Gstm3 | -1.46572 | 0.23694 | 6.17E-10 | 2.13E-07 |
| Plek | 1.346742 | 0.217869 | 6.35E-10 | 2.15E-07 |
| Msn | 1.644984 | 0.266294 | 6.52E-10 | 2.17E-07 |
| Mpeg1 | 1.449982 | 0.234938 | 6.75E-10 | 2.22E-07 |
| Heyl | 1.610178 | 0.261976 | 7.93E-10 | 2.56E-07 |
| C1qtnf6 | 1.598617 | 0.260941 | 8.99E-10 | 2.86E-07 |
| Snhg11 | -1.25001 | 0.204598 | 9.99E-10 | 3.13E-07 |
| Arg1 | 4.319687 | 0.709088 | 1.12E-09 | 3.44E-07 |
| Cx3cr1 | 1.901683 | 0.312646 | 1.18E-09 | 3.59E-07 |
| Gm26840 | 1.35093 | 0.223227 | 1.43E-09 | 4.28E-07 |
| Slc5a1 | -1.08471 | 0.180579 | 1.89E-09 | 5.58E-07 |
| D030025P2 | 3.818088 | 0.63797 | 2.17E-09 | 6.30E-07 |
| Plod2 | 1.1103 | 0.186854 | 2.81E-09 | 8.06E-07 |
| Mmp17 | 1.276233 | 0.214969 | 2.91E-09 | 8.21E-07 |
| Htr1b | 2.098923 | 0.357055 | 4.14E-09 | 1.14E-06 |
| Spock3 | -2.62209 | 0.451047 | 6.12E-09 | 1.66E-06 |
| Fxyd6 | 1.774952 | 0.305955 | 6.58E-09 | 1.76E-06 |
| Apbb1ip | 1.091339 | 0.189685 | 8.74E-09 | 2.31E-06 |
| Fcgr1 | 2.133065 | 0.370892 | 8.86E-09 | 2.31E-06 |
| Sfrp2 | 1.340396 | 0.233928 | 1.00E-08 | 2.59E-06 |
| Hdc | 2.134775 | 0.376146 | 1.38E-08 | 3.52E-06 |
| A330023F2 | -1.27876 | 0.22558 | 1.44E-08 | 3.61E-06 |
| Atcay | -2.15275 | 0.380015 | 1.47E-08 | 3.65E-06 |
| Chl1 | 1.902106 | 0.336897 | 1.64E-08 | 4.03E-06 |
| Timp1 | 3.937515 | 0.698207 | 1.71E-08 | 4.13E-06 |
| Tnfrsf1b | 1.179949 | 0.209957 | 1.91E-08 | 4.57E-06 |
| Lgmn | 1.087635 | 0.193769 | 1.99E-08 | 4.70E-06 |
| C6 | 2.294739 | 0.411088 | 2.38E-08 | 5.56E-06 |
| Fn1 | 1.455609 | 0.260937 | 2.43E-08 | 5.61E-06 |
| C1qb | 1.23144 | 0.221169 | 2.58E-08 | 5.89E-06 |

| | | | | |
|---|---|---|---|---|
| Sla | 1.572695 | 0.282847 | 2.69E-08 | 6.09E-06 |
| Fut8 | 1.07425 | 0.193345 | 2.76E-08 | 6.16E-06 |
| Igf1 | 1.724353 | 0.310476 | 2.79E-08 | 6.18E-06 |
| Sparc | 1.250195 | 0.225284 | 2.87E-08 | 6.27E-06 |
| Chst11 | 2.0225 | 0.364929 | 2.99E-08 | 6.46E-06 |
| Mmp2 | 1.358187 | 0.245984 | 3.36E-08 | 7.20E-06 |
| Gdf6 | 2.028124 | 0.367468 | 3.41E-08 | 7.22E-06 |
| Col4a1 | 1.163921 | 0.211713 | 3.85E-08 | 8.07E-06 |
| Aspn | 2.372694 | 0.434257 | 4.66E-08 | 9.58E-06 |
| Olfml3 | 1.454459 | 0.266128 | 4.62E-08 | 9.58E-06 |
| Itgam | 1.212232 | 0.222801 | 5.30E-08 | 1.08E-05 |
| Serpina3n | 2.997798 | 0.557037 | 7.38E-08 | 1.48E-05 |
| Filip1l | 1.461692 | 0.271673 | 7.44E-08 | 1.48E-05 |
| C1qtnf3 | 1.777581 | 0.330617 | 7.59E-08 | 1.50E-05 |
| Crhbp | -1.83938 | 0.34311 | 8.28E-08 | 1.62E-05 |
| Nptx2 | 1.930194 | 0.360941 | 8.91E-08 | 1.71E-05 |
| Col5a1 | 1.390083 | 0.260544 | 9.54E-08 | 1.81E-05 |
| Tlr13 | 1.50455 | 0.283629 | 1.13E-07 | 2.11E-05 |
| A830012C | 2.134503 | 0.402906 | 1.17E-07 | 2.17E-05 |
| Gm12867 | -2.34163 | 0.442792 | 1.23E-07 | 2.26E-05 |
| Cytip | 1.701275 | 0.322089 | 1.28E-07 | 2.32E-05 |
| Ccl6 | 1.719671 | 0.326481 | 1.38E-07 | 2.47E-05 |
| Fam111a | 1.314124 | 0.25015 | 1.49E-07 | 2.62E-05 |
| Igkv8-28 | 20.17584 | 3.841527 | 1.50E-07 | 2.62E-05 |
| Cd53 | 1.264615 | 0.240693 | 1.49E-07 | 2.62E-05 |
| Gm21451 | 2.443183 | 0.465785 | 1.56E-07 | 2.69E-05 |
| Adamts6 | 1.591315 | 0.304351 | 1.71E-07 | 2.92E-05 |
| Adamtsl2 | 1.568626 | 0.300896 | 1.86E-07 | 3.15E-05 |
| Laptm5 | 1.078828 | 0.207084 | 1.89E-07 | 3.18E-05 |
| Mfap2 | 1.338625 | 0.257423 | 1.99E-07 | 3.29E-05 |
| Gm42901 | 1.286532 | 0.247338 | 1.98E-07 | 3.29E-05 |
| Nckap1l | 1.180989 | 0.227848 | 2.18E-07 | 3.58E-05 |
| Igsf5 | -1.08204 | 0.20886 | 2.21E-07 | 3.60E-05 |
| Gda | 1.657901 | 0.32015 | 2.24E-07 | 3.61E-05 |
| Thsd7b | -3.16361 | 0.611206 | 2.27E-07 | 3.63E-05 |
| Il13ra1 | 1.076022 | 0.207979 | 2.29E-07 | 3.65E-05 |
| I830077J0 | 3.816325 | 0.7444 | 2.95E-07 | 4.65E-05 |
| C3ar1 | 1.279165 | 0.251182 | 3.53E-07 | 5.48E-05 |
| Snap91 | -1.47702 | 0.29243 | 4.40E-07 | 6.73E-05 |
| Lbp | 1.049798 | 0.208176 | 4.59E-07 | 6.96E-05 |
| Emb | 1.405174 | 0.278856 | 4.68E-07 | 7.05E-05 |
| Kng2 | 4.693589 | 0.932401 | 4.81E-07 | 7.08E-05 |
| Gm15270 | 2.1181 | 0.420538 | 4.74E-07 | 7.08E-05 |
| Kcnj8 | 1.463508 | 0.290663 | 4.78E-07 | 7.08E-05 |

| | | | | |
|---|---|---|---|---|
| Apobec2 | -2.47516 | 0.491883 | 4.85E-07 | 7.10E-05 |
| Fhl3 | 2.14882 | 0.427438 | 4.98E-07 | 7.23E-05 |
| Sprr2g | 3.616132 | 0.719517 | 5.01E-07 | 7.23E-05 |
| Ms4a6b | 1.653643 | 0.329496 | 5.20E-07 | 7.40E-05 |
| Phyhipl | -1.1476 | 0.228724 | 5.24E-07 | 7.40E-05 |
| Clec4a1 | 1.307379 | 0.261763 | 5.90E-07 | 8.27E-05 |
| Actr3b | -1.47647 | 0.296689 | 6.47E-07 | 8.96E-05 |
| Igkv9-120 | 7.195343 | 1.447543 | 6.67E-07 | 9.17E-05 |
| Clec4e | 4.007003 | 0.812924 | 8.26E-07 | 0.000111 |
| Col3a1 | 1.26829 | 0.257315 | 8.27E-07 | 0.000111 |
| Glis3 | 1.544715 | 0.314152 | 8.78E-07 | 0.000117 |
| Dbn1 | 1.119473 | 0.227666 | 8.78E-07 | 0.000117 |
| Vav3 | 1.029987 | 0.210672 | 1.01E-06 | 0.000133 |
| Gpr68 | 1.818157 | 0.372289 | 1.04E-06 | 0.000135 |
| Gm21975 | 1.418075 | 0.291059 | 1.10E-06 | 0.000142 |
| Masp1 | 2.049077 | 0.42105 | 1.14E-06 | 0.000145 |
| Syk | 1.041954 | 0.214961 | 1.25E-06 | 0.000159 |
| Vgf | 3.651522 | 0.754283 | 1.29E-06 | 0.000163 |
| Tgfb3 | 1.694817 | 0.350337 | 1.31E-06 | 0.000165 |
| Ngb | -2.02045 | 0.419508 | 1.46E-06 | 0.000183 |
| Fmod | 1.263347 | 0.262554 | 1.50E-06 | 0.000184 |
| Ccdc109b | 1.034249 | 0.215072 | 1.52E-06 | 0.000186 |
| Itgam | 1.535901 | 0.319654 | 1.55E-06 | 0.000189 |
| Tmprss11g | 3.259888 | 0.679339 | 1.60E-06 | 0.000193 |
| Plbd1 | 1.10483 | 0.230297 | 1.61E-06 | 0.000193 |
| Sulf1 | 1.049802 | 0.219392 | 1.71E-06 | 0.000202 |
| Ctss | 1.408643 | 0.294474 | 1.72E-06 | 0.000202 |
| Adamts4 | 3.659199 | 0.767226 | 1.85E-06 | 0.000216 |
| Smim3 | 1.235208 | 0.259927 | 2.01E-06 | 0.000234 |
| Rac2 | 1.161715 | 0.245435 | 2.21E-06 | 0.000254 |
| Gm28729 | -1.88367 | 0.398178 | 2.24E-06 | 0.000256 |
| Cxcl5 | 5.105131 | 1.081279 | 2.34E-06 | 0.000256 |
| Sirpb1c | 3.683313 | 0.779924 | 2.33E-06 | 0.000256 |
| Dio2 | 1.629894 | 0.345112 | 2.33E-06 | 0.000256 |
| Ripk3 | 1.614446 | 0.341886 | 2.33E-06 | 0.000256 |
| Pdgfrl | 1.384909 | 0.293338 | 2.34E-06 | 0.000256 |
| Glipr2 | 1.329301 | 0.28153 | 2.34E-06 | 0.000256 |
| Adam19 | 1.142802 | 0.241873 | 2.30E-06 | 0.000256 |
| Ereg | 1.723265 | 0.365612 | 2.44E-06 | 0.000264 |
| Fcgr2b | 1.364746 | 0.289528 | 2.43E-06 | 0.000264 |
| Tpsab1 | 6.561162 | 1.396149 | 2.61E-06 | 0.000276 |
| Adam8 | 1.700682 | 0.361827 | 2.60E-06 | 0.000276 |
| Lrrc75b | 1.334952 | 0.284136 | 2.62E-06 | 0.000276 |
| Fcer1a | 4.003345 | 0.852613 | 2.66E-06 | 0.000279 |

| | | | | |
|---|---|---|---|---|
| Sync | 1.164816 | 0.248115 | 2.67E-06 | 0.000279 |
| Il17b | 3.413045 | 0.728437 | 2.79E-06 | 0.00029 |
| Fam150a | 4.894461 | 1.048231 | 3.02E-06 | 0.000311 |
| Actg2 | 1.212164 | 0.260615 | 3.30E-06 | 0.000337 |
| C4b | 1.051447 | 0.22611 | 3.32E-06 | 0.000337 |
| Stat4 | 2.208692 | 0.475759 | 3.44E-06 | 0.000347 |
| Oprk1 | -1.17647 | 0.254367 | 3.74E-06 | 0.000373 |
| Htra4 | 1.47121 | 0.318195 | 3.77E-06 | 0.000374 |
| Gm37804 | 1.76536 | 0.38226 | 3.87E-06 | 0.00038 |
| Havcr2 | 1.822412 | 0.39508 | 3.97E-06 | 0.000389 |
| Figf | 1.289142 | 0.279683 | 4.04E-06 | 0.000391 |
| Gm266 | -1.8473 | 0.400767 | 4.04E-06 | 0.000391 |
| Ms4a6c | 1.575618 | 0.342134 | 4.12E-06 | 0.000395 |
| Kif5a | -1.15269 | 0.250273 | 4.11E-06 | 0.000395 |
| Angpt4 | 2.058373 | 0.447264 | 4.18E-06 | 0.000399 |
| Ak5 | 1.494464 | 0.324912 | 4.23E-06 | 0.000402 |
| Pirb | 1.412904 | 0.307655 | 4.38E-06 | 0.000414 |
| Ccl9 | 1.574467 | 0.343058 | 4.44E-06 | 0.000418 |
| P2ry6 | 1.153177 | 0.251426 | 4.51E-06 | 0.000422 |
| Gdf11 | 1.154343 | 0.251744 | 4.53E-06 | 0.000423 |
| Gm13372 | 1.381821 | 0.301772 | 4.67E-06 | 0.000434 |
| C530008M | -1.5317 | 0.33525 | 4.90E-06 | 0.000453 |
| Ccr1 | 1.964281 | 0.430064 | 4.94E-06 | 0.000454 |
| Vav1 | 1.390533 | 0.304772 | 5.05E-06 | 0.000459 |
| Gpx2 | 1.371808 | 0.300662 | 5.05E-06 | 0.000459 |
| Postn | 1.012155 | 0.221814 | 5.04E-06 | 0.000459 |
| Gpr176 | 2.620398 | 0.574592 | 5.10E-06 | 0.000461 |
| Tmem173 | 1.304969 | 0.286406 | 5.20E-06 | 0.000468 |
| Maff | 1.404455 | 0.308315 | 5.23E-06 | 0.000469 |
| Fbln2 | 1.4088 | 0.309619 | 5.36E-06 | 0.000478 |
| Ednra | 1.526933 | 0.335657 | 5.39E-06 | 0.000479 |
| Tmem200c | -1.85989 | 0.409251 | 5.50E-06 | 0.000487 |
| Coro1a | 1.036185 | 0.228349 | 5.69E-06 | 0.0005 |
| Mamdc2 | -1.95319 | 0.430506 | 5.71E-06 | 0.0005 |
| Gpbar1 | 4.647466 | 1.02471 | 5.75E-06 | 0.000502 |
| Ism1 | 1.277395 | 0.282664 | 6.21E-06 | 0.00054 |
| Atp8b4 | 2.088026 | 0.462235 | 6.27E-06 | 0.000542 |
| Nxpe5 | 2.150388 | 0.477527 | 6.69E-06 | 0.000577 |
| Aldh1a2 | 1.23218 | 0.274217 | 7.01E-06 | 0.000601 |
| Clec5a | 1.926252 | 0.431293 | 7.96E-06 | 0.000672 |
| Klhl6 | 1.253678 | 0.280599 | 7.90E-06 | 0.000672 |
| Rftn1 | 1.101544 | 0.246571 | 7.92E-06 | 0.000672 |
| Dio3 | 2.16879 | 0.486677 | 8.34E-06 | 0.000701 |
| Ntm | -1.99322 | 0.447589 | 8.46E-06 | 0.000708 |

| | | | | |
|---|---|---|---|---|
| Col4a2 | 1.056174 | 0.237323 | 8.57E-06 | 0.000712 |
| Hs3st3b1 | 2.219278 | 0.498937 | 8.67E-06 | 0.000715 |
| 6030408B: | 2.258272 | 0.508155 | 8.83E-06 | 0.000724 |
| Gm26586 | 1.257055 | 0.282979 | 8.90E-06 | 0.000727 |
| Clip4 | 1.160834 | 0.261587 | 9.09E-06 | 0.000737 |
| Plch1 | 2.377202 | 0.536157 | 9.26E-06 | 0.000747 |
| 2210407C: | 3.434706 | 0.775825 | 9.55E-06 | 0.000765 |
| Medag | 1.319018 | 0.297938 | 9.55E-06 | 0.000765 |
| Fam196b | -1.20772 | 0.273206 | 9.84E-06 | 0.000785 |
| Reg3g | 4.775353 | 1.082802 | 1.03E-05 | 0.000814 |
| Gdf3 | 4.048435 | 0.918362 | 1.04E-05 | 0.000818 |
| Ly86 | 1.342777 | 0.304977 | 1.07E-05 | 0.000832 |
| Adgra1 | 2.036205 | 0.463448 | 1.11E-05 | 0.000866 |
| Fcgr3 | 1.272518 | 0.29025 | 1.16E-05 | 0.0009 |
| 2810430I1 | 1.699216 | 0.387755 | 1.17E-05 | 0.000905 |
| Chil1 | 3.162604 | 0.722246 | 1.19E-05 | 0.000916 |
| Il4ra | 1.317438 | 0.300938 | 1.20E-05 | 0.000917 |
| Tifab | 1.269783 | 0.29081 | 1.26E-05 | 0.000959 |
| Hmcn2 | -1.48689 | 0.340681 | 1.27E-05 | 0.00096 |
| Epha3 | 1.101444 | 0.252805 | 1.32E-05 | 0.000986 |
| AC125167. | 1.679034 | 0.386064 | 1.37E-05 | 0.001018 |
| Trim46 | 1.183331 | 0.272152 | 1.37E-05 | 0.00102 |
| Mnda | 1.461166 | 0.33637 | 1.40E-05 | 0.001035 |
| Cd300lf | 4.026037 | 0.927419 | 1.42E-05 | 0.001036 |
| Epyc | 2.40489 | 0.554115 | 1.42E-05 | 0.001036 |
| Ms4a7 | 1.481881 | 0.341517 | 1.43E-05 | 0.001036 |
| Tpm4 | 1.082871 | 0.249502 | 1.42E-05 | 0.001036 |
| Mmp7 | 10.36886 | 2.392306 | 1.46E-05 | 0.001051 |
| Itgbl1 | 1.677971 | 0.387113 | 1.46E-05 | 0.001051 |
| Arsj | 1.055662 | 0.244514 | 1.58E-05 | 0.001127 |
| Nov | 1.486475 | 0.344532 | 1.60E-05 | 0.001138 |
| Fgr | 2.320537 | 0.538469 | 1.64E-05 | 0.00116 |
| Col17a1 | 1.038516 | 0.241175 | 1.66E-05 | 0.001174 |
| Mefv | 2.21487 | 0.51475 | 1.69E-05 | 0.001187 |
| Was | 1.280141 | 0.29841 | 1.79E-05 | 0.001243 |
| Kdelr3 | 1.27343 | 0.297524 | 1.87E-05 | 0.001288 |
| Evi2b | 1.42363 | 0.332923 | 1.90E-05 | 0.001307 |
| Ly6c2 | 1.918394 | 0.448757 | 1.91E-05 | 0.00131 |
| Lrrc15 | 1.701213 | 0.398093 | 1.93E-05 | 0.001314 |
| Serpina3m | 3.6172 | 0.847276 | 1.96E-05 | 0.001334 |
| P2ry10 | 1.650662 | 0.387722 | 2.07E-05 | 0.001403 |
| Clec4n | 1.758104 | 0.413586 | 2.13E-05 | 0.001439 |
| Sox11 | 3.790325 | 0.892558 | 2.17E-05 | 0.001462 |
| Gm29233 | 1.725338 | 0.406777 | 2.22E-05 | 0.001491 |

| | | | | |
|---|---|---|---|---|
| Plekhh2 | 1.067698 | 0.251863 | 2.24E-05 | 0.001492 |
| Camk2n2 | 1.067554 | 0.251857 | 2.25E-05 | 0.001492 |
| Gm16033 | 1.036422 | 0.244535 | 2.25E-05 | 0.001492 |
| Mmp9 | -1.63748 | 0.38635 | 2.25E-05 | 0.001492 |
| Akap5 | 1.750292 | 0.414124 | 2.37E-05 | 0.001568 |
| Cd84 | 1.310287 | 0.310287 | 2.41E-05 | 0.001588 |
| Pik3ap1 | 1.461281 | 0.346305 | 2.45E-05 | 0.001605 |
| Gm20488 | 1.362094 | 0.323299 | 2.52E-05 | 0.001642 |
| F2r | 1.455587 | 0.345958 | 2.58E-05 | 0.001679 |
| Gm17509 | 1.67309 | 0.398252 | 2.66E-05 | 0.001721 |
| Inpp5d | 1.029571 | 0.245151 | 2.67E-05 | 0.001726 |
| Rtn4rl2 | 2.47589 | 0.589945 | 2.71E-05 | 0.001742 |
| Bin2 | 1.017721 | 0.24257 | 2.72E-05 | 0.001745 |
| Ndnf | 1.750736 | 0.418163 | 2.83E-05 | 0.001799 |
| Tro | 1.676092 | 0.400317 | 2.83E-05 | 0.001799 |
| Penk | 1.764726 | 0.423031 | 3.02E-05 | 0.00191 |
| Trem1 | 3.660018 | 0.877795 | 3.05E-05 | 0.001916 |
| Mt2 | 1.514876 | 0.363302 | 3.05E-05 | 0.001916 |
| Inhba | 1.969156 | 0.47336 | 3.18E-05 | 0.001986 |
| Plk5 | -2.70666 | 0.650559 | 3.18E-05 | 0.001986 |
| Slc26a7 | 1.646853 | 0.396066 | 3.21E-05 | 0.001993 |
| Gm26771 | 1.085403 | 0.261058 | 3.21E-05 | 0.001993 |
| Gpr35 | 1.283599 | 0.309982 | 3.46E-05 | 0.00212 |
| Tspan11 | 1.648604 | 0.398277 | 3.48E-05 | 0.002121 |
| Arhgap30 | 1.007618 | 0.243392 | 3.47E-05 | 0.002121 |
| Fstl3 | 1.344072 | 0.324777 | 3.50E-05 | 0.002123 |
| Slc41a2 | 1.275875 | 0.308473 | 3.53E-05 | 0.002138 |
| Ccl8 | 2.802709 | 0.678332 | 3.60E-05 | 0.002173 |
| Tnnt2 | -1.04707 | 0.254451 | 3.87E-05 | 0.002309 |
| Klhl30 | -1.41703 | 0.344342 | 3.87E-05 | 0.002309 |
| Prnd | 2.590787 | 0.629823 | 3.90E-05 | 0.002314 |
| D430036J1 | 2.063694 | 0.501728 | 3.90E-05 | 0.002314 |
| Clec4d | 2.596786 | 0.631683 | 3.94E-05 | 0.002324 |
| Adra1d | 2.225214 | 0.541216 | 3.93E-05 | 0.002324 |
| Eno3 | -1.77209 | 0.431309 | 3.98E-05 | 0.00234 |
| Tlr1 | 1.666939 | 0.405939 | 4.02E-05 | 0.002356 |
| Gria1 | -2.87681 | 0.700846 | 4.05E-05 | 0.002365 |
| Rgs4 | 2.530733 | 0.616677 | 4.06E-05 | 0.002368 |
| Casp4 | 1.076851 | 0.262459 | 4.08E-05 | 0.002371 |
| Gm11744 | 1.061768 | 0.258884 | 4.11E-05 | 0.00238 |
| S100a8 | 3.535999 | 0.862633 | 4.15E-05 | 0.002397 |
| Igfbp2 | 1.126933 | 0.275235 | 4.23E-05 | 0.002438 |
| Cyth4 | 1.086323 | 0.265402 | 4.26E-05 | 0.002445 |
| Gjb1 | 3.44038 | 0.841641 | 4.36E-05 | 0.002489 |

| | | | | |
|---|---|---|---|---|
| Lrg1 | 1.416938 | 0.347026 | 4.44E-05 | 0.00253 |
| Col12a1 | 1.087936 | 0.267327 | 4.71E-05 | 0.002645 |
| Synpo | 1.043626 | 0.256422 | 4.70E-05 | 0.002645 |
| Gm9733 | 6.204773 | 1.525715 | 4.77E-05 | 0.002671 |
| Fxyd5 | 1.152376 | 0.283947 | 4.94E-05 | 0.002761 |
| Ccdc80 | 1.061535 | 0.261657 | 4.97E-05 | 0.00277 |
| Dlk1 | -2.65321 | 0.654592 | 5.05E-05 | 0.002807 |
| Vcan | 1.598739 | 0.395457 | 5.28E-05 | 0.00292 |
| Lrrc32 | 1.601059 | 0.396205 | 5.32E-05 | 0.00293 |
| 01-Mar | 1.549687 | 0.383561 | 5.34E-05 | 0.00293 |
| Cd38 | 1.167389 | 0.288957 | 5.34E-05 | 0.00293 |
| 4933431K2 | 2.313888 | 0.573107 | 5.40E-05 | 0.002955 |
| Spsb1 | 1.448466 | 0.358855 | 5.43E-05 | 0.00296 |
| Tlr8 | 1.928952 | 0.478006 | 5.45E-05 | 0.002964 |
| Fam179a | 1.936461 | 0.480145 | 5.51E-05 | 0.002976 |
| Gpr65 | 1.516014 | 0.375938 | 5.52E-05 | 0.002976 |
| Wfdc17 | 1.875788 | 0.465562 | 5.60E-05 | 0.003013 |
| C1ql3 | 2.361843 | 0.586354 | 5.63E-05 | 0.003019 |
| Htr7 | 2.835048 | 0.704675 | 5.74E-05 | 0.003061 |
| Hp | 2.541232 | 0.631694 | 5.75E-05 | 0.003061 |
| Thbs1 | 1.557654 | 0.387295 | 5.77E-05 | 0.003066 |
| Kl | -1.02524 | 0.254966 | 5.79E-05 | 0.003068 |
| Padi3 | 1.615187 | 0.40203 | 5.88E-05 | 0.003106 |
| Msln | 1.413599 | 0.351919 | 5.90E-05 | 0.003108 |
| Gng8 | 1.324 | 0.329942 | 6.00E-05 | 0.003153 |
| Tmem200a | 1.77908 | 0.443754 | 6.09E-05 | 0.003194 |
| 1810011H1 | 1.624423 | 0.406986 | 6.57E-05 | 0.003418 |
| Il1rl2 | 1.022964 | 0.257159 | 6.95E-05 | 0.003598 |
| Rhoc | 1.076521 | 0.270763 | 7.01E-05 | 0.0036 |
| Dock3 | -1.06072 | 0.26682 | 7.03E-05 | 0.0036 |
| Ramp3 | 1.588367 | 0.399728 | 7.08E-05 | 0.003618 |
| Lilrb4a | 1.335183 | 0.336072 | 7.10E-05 | 0.003619 |
| Igkj5 | 5.982519 | 1.508783 | 7.34E-05 | 0.00373 |
| Dnm3os | 1.008243 | 0.254441 | 7.41E-05 | 0.003761 |
| Tgm1 | 1.593142 | 0.402795 | 7.65E-05 | 0.003869 |
| Ipcef1 | -1.11502 | 0.282023 | 7.70E-05 | 0.003875 |
| Cbs | -1.34222 | 0.3399 | 7.85E-05 | 0.003943 |
| Rgs16 | 1.301178 | 0.329804 | 7.97E-05 | 0.003985 |
| Dok7 | -1.71612 | 0.43532 | 8.07E-05 | 0.004015 |
| BE692007 | 1.463564 | 0.372397 | 8.49E-05 | 0.004192 |
| Eda2r | 1.28101 | 0.326397 | 8.68E-05 | 0.004277 |
| Selp | 1.58632 | 0.405823 | 9.27E-05 | 0.004544 |
| Rapgef4os1 | -1.1805 | 0.302048 | 9.29E-05 | 0.004544 |
| Chrm2 | 1.495636 | 0.250611 | 6.80E-05 | 0.004549 |

| | | | | |
|---|---|---|---|---|
| Trem2 | 1.856651 | 0.475347 | 9.39E-05 | 0.004579 |
| Ccr7 | 1.80926 | 0.464274 | 9.74E-05 | 0.004717 |
| Samsn1 | 1.590715 | 0.40819 | 9.74E-05 | 0.004717 |
| Clec11a | 1.029045 | 0.264494 | 1.00E-04 | 0.004831 |
| Fam198b | 1.386123 | 0.356531 | 0.000101 | 0.004876 |
| Igsf6 | 1.265916 | 0.325668 | 0.000101 | 0.004878 |
| Il33 | 2.060362 | 0.530705 | 0.000103 | 0.004956 |
| Slc16a3 | 1.1866 | 0.30566 | 0.000104 | 0.004956 |
| Pycr1 | 1.290695 | 0.333066 | 0.000107 | 0.005087 |
| Peg12 | 1.116539 | 0.288222 | 0.000107 | 0.00509 |
| Clec1a | 1.2425 | 0.321165 | 0.000109 | 0.005175 |
| Usp2 | -1.46489 | 0.378786 | 0.00011 | 0.005193 |
| Snx20 | 1.017692 | 0.26359 | 0.000113 | 0.005307 |
| Tmem114 | 5.66948 | 1.469563 | 0.000114 | 0.005359 |
| Serpina3g | 2.381099 | 0.617877 | 0.000116 | 0.005431 |
| Hspb1 | 1.289032 | 0.334506 | 0.000116 | 0.005431 |
| Ly6a | 1.158465 | 0.300717 | 0.000117 | 0.005433 |
| Capn6 | 1.078316 | 0.27988 | 0.000117 | 0.005433 |
| Kcnn2 | -1.6415 | 0.426173 | 0.000117 | 0.005434 |
| Tlr7 | 1.303 | 0.338339 | 0.000118 | 0.005434 |
| Bst1 | 1.025561 | 0.266502 | 0.000119 | 0.005487 |
| Mir6236 | -1.56917 | 0.407889 | 0.00012 | 0.005501 |
| Tyrobp | 1.177322 | 0.306263 | 0.000121 | 0.005542 |
| Slfn1 | 2.504942 | 0.651779 | 0.000121 | 0.00555 |
| Fkbp11 | 1.565118 | 0.407531 | 0.000123 | 0.0056 |
| Lair1 | 1.623197 | 0.423033 | 0.000125 | 0.005657 |
| Aldh1l2 | 1.577585 | 0.411162 | 0.000125 | 0.005657 |
| Txlnb | -1.34878 | 0.351583 | 0.000125 | 0.005658 |
| Clec4a3 | 1.404572 | 0.366732 | 0.000128 | 0.005775 |
| Galnt5 | 1.11946 | 0.292314 | 0.000128 | 0.005775 |
| Arhgap15 | 1.277897 | 0.333784 | 0.000129 | 0.005788 |
| Nav3 | 1.575812 | 0.412143 | 0.000132 | 0.005872 |
| Uhrf1 | 1.320664 | 0.345378 | 0.000131 | 0.005872 |
| Il21r | 1.62463 | 0.425175 | 0.000133 | 0.005914 |
| Adcy7 | 1.096025 | 0.286988 | 0.000134 | 0.005949 |
| Ptger3 | 1.027979 | 0.269702 | 0.000138 | 0.00608 |
| Peg3 | -1.31418 | 0.344975 | 0.000139 | 0.006111 |
| Ryr2 | -1.39905 | 0.367277 | 0.000139 | 0.006111 |
| Cys1 | -1.30569 | 0.343964 | 0.000147 | 0.006405 |
| Fcer1g | 1.189038 | 0.313306 | 0.000148 | 0.006414 |
| Siglece | 1.653547 | 0.435834 | 0.000148 | 0.006416 |
| Lrrc25 | 1.194391 | 0.314793 | 0.000148 | 0.006416 |
| Igkv4-53 | 6.875817 | 1.81335 | 0.00015 | 0.006447 |
| Cgref1 | 1.210583 | 0.31927 | 0.00015 | 0.006447 |

| | | | | |
|---|---|---|---|---|
| Ighv1-53 | 8.445491 | 2.228272 | 0.000151 | 0.006474 |
| Col11a1 | 1.296801 | 0.342204 | 0.000151 | 0.006476 |
| Il6 | 3.351444 | 0.885529 | 0.000154 | 0.006577 |
| Gm13889 | 1.546902 | 0.408871 | 0.000155 | 0.006584 |
| Fbn2 | 4.284433 | 1.134156 | 0.000158 | 0.006709 |
| Bmp10 | -1.23683 | 0.327799 | 0.000161 | 0.006817 |
| Serpinb1a | 1.049782 | 0.279029 | 0.000168 | 0.00705 |
| Rgs6 | -1.10106 | 0.292668 | 0.000168 | 0.00705 |
| Cxcr2 | 2.490822 | 0.66228 | 0.000169 | 0.007069 |
| Stac2 | 2.073058 | 0.551314 | 0.00017 | 0.007076 |
| Msr1 | 1.260302 | 0.335301 | 0.000171 | 0.007104 |
| Rsad2 | 1.715384 | 0.457059 | 0.000175 | 0.007236 |
| Cstad | -1.80294 | 0.481609 | 0.000181 | 0.00747 |
| Pcdh8 | 3.430573 | 0.918014 | 0.000186 | 0.007608 |
| Ighv1-82 | 6.28781 | 1.684225 | 0.000189 | 0.007686 |
| Trac | 2.150497 | 0.576225 | 0.00019 | 0.007711 |
| Mzb1 | 3.66999 | 0.984118 | 0.000192 | 0.007767 |
| Cysltr1 | 1.04231 | 0.280165 | 0.000199 | 0.007959 |
| Pigr | -1.1831 | 0.318005 | 0.000199 | 0.007959 |
| Ptpn22 | 1.633621 | 0.439951 | 0.000205 | 0.008153 |
| Cdr2l | 1.133792 | 0.305692 | 0.000208 | 0.008252 |
| Ly9 | 1.432989 | 0.386647 | 0.00021 | 0.008326 |
| Kcnf1 | -1.25783 | 0.339526 | 0.000212 | 0.008345 |
| Ms4a4a | 1.068635 | 0.288714 | 0.000214 | 0.008437 |
| Themis2 | 1.08003 | 0.291905 | 0.000216 | 0.008468 |
| Ace | 1.165268 | 0.315021 | 0.000216 | 0.00847 |
| Il1rl1 | 1.268565 | 0.344114 | 0.000227 | 0.008827 |
| Alox5ap | 1.052733 | 0.286267 | 0.000236 | 0.009098 |
| Sh2d5 | 2.263577 | 0.616146 | 0.000239 | 0.009207 |
| Qrfpr | -3.44601 | 0.938401 | 0.00024 | 0.009246 |
| B4galt5 | 1.330939 | 0.363512 | 0.000251 | 0.009575 |
| Emp1 | 1.040416 | 0.284583 | 0.000256 | 0.009742 |
| Gm13070 | 1.784515 | 0.488887 | 0.000262 | 0.009915 |
| Pla2g2d | -1.55693 | 0.426818 | 0.000265 | 0.009946 |

Supplemental Table 4. LRT significant DEGs

| GeneID | Log2 fc | SE | p-value | FDR p-value |
|---|---|---|---|---|
| Syt14 | 3.91471 | 0.506127 | 3.30E-14 | 6.33E-10 |
| Hrh3 | 4.338709 | 0.642005 | 3.25E-11 | 3.12E-07 |
| Dcdc2a | 5.833356 | 1.174525 | 1.10E-10 | 7.02E-07 |
| NA | 4.923966 | 0.831859 | 4.44E-10 | 2.13E-06 |
| Bglap2 | 3.96941 | 0.63396 | 1.49E-09 | 5.73E-06 |
| Rgs5 | 2.695646 | 0.446638 | 3.40E-09 | 1.01E-05 |
| Galnt5 | 3.277409 | 0.519004 | 3.67E-09 | 1.01E-05 |
| Cnnm1 | 2.511086 | 0.426744 | 4.23E-09 | 1.01E-05 |
| Rrad | -2.26002 | 0.41611 | 5.08E-09 | 1.08E-05 |
| Ciart | 2.523121 | 0.465919 | 6.37E-09 | 1.22E-05 |
| Slc9a2 | 2.317098 | 0.391383 | 1.56E-08 | 2.73E-05 |
| Ipcef1 | 3.19449 | 0.561668 | 1.93E-08 | 3.08E-05 |
| Fbxw27 | 3.242289 | 0.57906 | 2.36E-08 | 3.48E-05 |
| Slc16a12 | 2.836723 | 0.546306 | 4.14E-08 | 5.67E-05 |
| Ubfd1 | -0.51041 | 0.134484 | 5.09E-08 | 6.51E-05 |
| Tc2n | 2.160142 | 0.380065 | 1.00E-07 | 0.00012 |
| Dusp8 | -4.28813 | 0.799391 | 1.45E-07 | 0.000163 |
| Rxfp1 | 4.434449 | 0.813689 | 1.62E-07 | 0.000172 |
| Esr2 | 4.103346 | 0.859103 | 2.32E-07 | 0.000222 |
| Kl | 2.902566 | 0.540726 | 2.29E-07 | 0.000222 |
| 4930579G[illegible] | -1.72554 | 0.355899 | 6.19E-07 | 0.00054 |
| Faah | 1.333272 | 0.27572 | 6.06E-07 | 0.00054 |
| Enc1 | -1.82294 | 0.35487 | 6.80E-07 | 0.000567 |
| Uchl1 | 1.876199 | 0.356849 | 7.52E-07 | 0.000601 |
| Ager | 2.61143 | 0.523379 | 1.05E-06 | 0.000745 |
| Dhtkd1 | 1.896027 | 0.367978 | 1.02E-06 | 0.000745 |
| Cr2 | 4.314682 | 1.192427 | 1.05E-06 | 0.000745 |
| Skap1 | 2.455692 | 0.52221 | 1.19E-06 | 0.000812 |
| Dmrtc1a | 3.98526 | 0.819197 | 1.32E-06 | 0.000875 |
| Sdr16c6 | 3.311273 | 0.766782 | 1.42E-06 | 0.000909 |
| Cacna1b | 3.399782 | 0.679442 | 1.57E-06 | 0.000944 |
| Slc5a1 | 2.832177 | 0.646726 | 1.57E-06 | 0.000944 |
| Gadd45g | -2.81813 | 0.590585 | 1.66E-06 | 0.000966 |
| Gm13807 | 3.048213 | 0.603146 | 1.85E-06 | 0.001046 |
| Nr1d1 | 2.54013 | 0.497641 | 2.47E-06 | 0.001349 |
| Ccl2 | -4.56448 | 0.910147 | 2.53E-06 | 0.001349 |
| Nes | -2.06072 | 0.427671 | 3.02E-06 | 0.001533 |
| Hnrnpm | -1.35772 | 0.296781 | 3.04E-06 | 0.001533 |
| Wfdc15b | 1.744112 | 0.360512 | 4.04E-06 | 0.001987 |
| Ywhah | -1.3548 | 0.275562 | 4.25E-06 | 0.002006 |
| NA | 5.229367 | 1.111553 | 4.29E-06 | 0.002006 |
| Zfp346 | 0.647319 | 0.159179 | 5.25E-06 | 0.002396 |

| | | | | |
|---|---|---|---|---|
| Ear2 | 2.402089 | 0.72285 | 5.39E-06 | 0.002403 |
| Rasd2 | 2.271317 | 0.454909 | 5.91E-06 | 0.002575 |
| Rab17 | 4.169957 | 0.834048 | 6.47E-06 | 0.002725 |
| Myoc | 4.091965 | 0.85633 | 6.53E-06 | 0.002725 |
| Smoc1 | 1.285371 | 0.26377 | 6.76E-06 | 0.002744 |
| Dbp | 3.873532 | 0.796457 | 6.87E-06 | 0.002744 |
| Arid3c | 3.43146 | 0.725202 | 7.32E-06 | 0.002865 |
| Ccar1 | -0.99798 | 0.244327 | 7.50E-06 | 0.002878 |
| Cpne6 | 3.33614 | 1.042779 | 8.19E-06 | 0.00302 |
| Upb1 | 2.908182 | 0.613972 | 8.06E-06 | 0.00302 |
| Cdh3 | -2.05114 | 0.4261 | 9.02E-06 | 0.003265 |
| NA | -3.57373 | 0.896156 | 9.78E-06 | 0.003474 |
| H2-M5 | 2.450901 | 0.635361 | 1.03E-05 | 0.003585 |
| Ntrk3 | 1.58453 | 0.506647 | 1.19E-05 | 0.00407 |
| Galnt16 | 1.931094 | 0.456906 | 1.28E-05 | 0.004317 |
| Gkn3 | 3.200672 | 0.672751 | 1.31E-05 | 0.004344 |
| Snrnp35 | -0.94129 | 0.199886 | 1.38E-05 | 0.004477 |
| NA | 3.195749 | 0.667747 | 1.57E-05 | 0.005009 |
| Cpm | 2.434804 | 0.509535 | 1.70E-05 | 0.005256 |
| Nr1d2 | 1.45775 | 0.328885 | 1.69E-05 | 0.005256 |
| Kcnmb4os: | 2.623882 | 0.574164 | 1.79E-05 | 0.005462 |
| Suv39h1 | -1.65766 | 0.405454 | 1.98E-05 | 0.005944 |
| Asic2 | 3.129582 | 0.750577 | 2.06E-05 | 0.006083 |
| Gm14412 | 2.665465 | 0.595866 | 2.46E-05 | 0.007154 |
| NA | 1.033379 | 0.555155 | 2.61E-05 | 0.007464 |
| 2310002F( | 2.068709 | 0.458461 | 2.91E-05 | 0.008198 |
| NA | -3.53501 | 0.791325 | 2.96E-05 | 0.00822 |
| Scn4b | 2.254372 | 0.532364 | 3.00E-05 | 0.008222 |
| Mr1 | 0.974635 | 0.221592 | 3.22E-05 | 0.008711 |
| Lrrc15 | 2.622842 | 0.814273 | 3.40E-05 | 0.009065 |
| Grip2 | 2.699638 | 0.700959 | 3.69E-05 | 0.009483 |
| Grap2 | 1.895089 | 0.418831 | 3.71E-05 | 0.009483 |
| Ticrr | -5.37985 | 1.462957 | 3.63E-05 | 0.009483 |
| Rsrp1 | 1.42432 | 0.358833 | 3.85E-05 | 0.00972 |
| Ptgs2 | -4.58929 | 1.156007 | 3.91E-05 | 0.009747 |
| Dnah14 | -1.74956 | 0.41496 | 3.97E-05 | 0.009764 |
| Dusp10 | -2.40837 | 0.558978 | 4.04E-05 | 0.009803 |
| Csnk1e | -0.85363 | 0.219865 | 4.22E-05 | 0.010125 |
| NA | -2.19866 | 0.565138 | 4.30E-05 | 0.010188 |
| NA | 3.283779 | 0.729267 | 4.41E-05 | 0.010306 |
| Zscan26 | 1.061098 | 0.236245 | 4.58E-05 | 0.010455 |
| Magea9 | 1.97217 | 0.674156 | 4.54E-05 | 0.010455 |
| Mex3a | -1.19753 | 0.328784 | 4.65E-05 | 0.010501 |
| Snrnp70 | -1.07613 | 0.283841 | 4.76E-05 | 0.010612 |

| | | | | |
|---|---|---|---|---|
| Cd163 | -1.02533 | 0.331647 | 4.88E-05 | 0.010759 |
| Plk4 | -2.40972 | 0.608975 | 5.20E-05 | 0.011301 |
| 1700019D( | 2.028105 | 0.450957 | 5.24E-05 | 0.011301 |
| Galnt15 | 1.891435 | 0.558168 | 5.37E-05 | 0.011452 |
| Slco4a1 | -3.64443 | 0.963802 | 5.61E-05 | 0.011835 |
| NA | 2.285241 | 0.522629 | 5.72E-05 | 0.011928 |
| Gdf15 | -5.51907 | 1.23408 | 6.10E-05 | 0.012588 |
| Gadd45b | -1.71661 | 0.449983 | 6.37E-05 | 0.012623 |
| Arhgef3 | 1.059507 | 0.252684 | 6.30E-05 | 0.012623 |
| Rgs11 | 1.388218 | 0.323096 | 6.38E-05 | 0.012623 |
| Ckap4 | -1.36348 | 0.315844 | 6.37E-05 | 0.012623 |
| NA | 1.5351 | 0.3638 | 6.66E-05 | 0.013031 |
| Nasp | -2.19818 | 0.545704 | 6.86E-05 | 0.013165 |
| NA | 2.377106 | 0.553532 | 6.86E-05 | 0.013165 |
| Slc6a18 | 2.990496 | 0.726567 | 6.97E-05 | 0.013243 |
| Cdh26 | 4.7345 | 1.206773 | 7.12E-05 | 0.013381 |
| Vipr1 | 1.452738 | 0.359318 | 7.48E-05 | 0.013935 |
| Esyt3 | 2.040582 | 0.461581 | 7.73E-05 | 0.014263 |
| Foxs1 | -3.80152 | 0.86279 | 7.97E-05 | 0.014568 |
| Smc2 | -2.48236 | 0.727802 | 8.17E-05 | 0.014687 |
| Fbxw19 | 6.017022 | 1.487068 | 8.19E-05 | 0.014687 |
| Cpa3 | 3.353492 | 1.012021 | 8.82E-05 | 0.015021 |
| Mybl2 | -3.60361 | 0.952263 | 8.93E-05 | 0.015021 |
| Marcks | -0.85747 | 0.243552 | 8.71E-05 | 0.015021 |
| Zfp872 | 1.772952 | 0.422091 | 8.88E-05 | 0.015021 |
| Gm8355 | -1.61425 | 0.391251 | 8.85E-05 | 0.015021 |
| NA | 3.448044 | 0.97219 | 8.73E-05 | 0.015021 |
| Sox2ot | 2.790834 | 0.685384 | 8.75E-05 | 0.015021 |
| NA | 1.924929 | 0.446812 | 9.30E-05 | 0.015521 |
| Dusp5 | -2.69743 | 0.642873 | 0.000101 | 0.016713 |
| Zfp273 | 1.029455 | 0.263105 | 0.000103 | 0.016836 |
| Haus8 | -0.89398 | 0.250375 | 0.000105 | 0.017133 |
| NA | -0.97102 | 0.323862 | 0.000106 | 0.017156 |
| Ccdc34 | -1.42429 | 0.462668 | 0.000108 | 0.017251 |
| Slc6a4 | 0.908285 | 0.767398 | 0.00011 | 0.017251 |
| Cdca8 | -3.08129 | 0.912012 | 0.000109 | 0.017251 |
| Rap1gap | 1.03707 | 0.24358 | 0.000113 | 0.017456 |
| NA | 2.286745 | 0.76373 | 0.000113 | 0.017456 |
| Myef2 | -1.38614 | 0.357827 | 0.000116 | 0.017858 |
| Taf15 | -1.05234 | 0.313209 | 0.000118 | 0.017868 |
| Gm2762 | 1.368106 | 0.347248 | 0.000118 | 0.017868 |
| Gpr146 | 1.302924 | 0.323846 | 0.00012 | 0.017874 |
| NA | 1.01602 | 0.429602 | 0.00012 | 0.017874 |
| 4933406C: | 2.21431 | 0.522045 | 0.000124 | 0.018259 |

| | | | | |
|---|---|---|---|---|
| Arg2 | 1.969815 | 0.468452 | 0.000132 | 0.019387 |
| Kif20a | -2.19202 | 0.645928 | 0.000135 | 0.019572 |
| NA | 2.043082 | 0.477242 | 0.000136 | 0.019627 |
| Mertk | 0.963315 | 0.241737 | 0.000139 | 0.019798 |
| Ccn2 | -3.71772 | 0.93492 | 0.000139 | 0.019798 |
| Tsc22d2 | -1.80464 | 0.451487 | 0.000142 | 0.019798 |
| Cntnap3 | 3.048916 | 0.722284 | 0.000142 | 0.019798 |
| Tnfaip6 | -3.80297 | 1.004435 | 0.000141 | 0.019798 |
| E2f1 | -2.17685 | 0.604391 | 0.000145 | 0.020027 |
| Scube2 | 3.068617 | 0.708766 | 0.000152 | 0.020275 |
| Lnx1 | 0.776375 | 0.184658 | 0.000152 | 0.020275 |
| Itgax | 2.058114 | 0.512531 | 0.000152 | 0.020275 |
| Smc4 | -1.74207 | 0.490927 | 0.000152 | 0.020275 |
| F13a1 | -0.67683 | 0.321223 | 0.000148 | 0.020275 |
| Erdr1x | -4.21943 | 1.05002 | 0.000156 | 0.020679 |
| Ccna2 | -2.65848 | 0.804611 | 0.000161 | 0.021151 |
| Kcnmb4 | 2.676158 | 0.659509 | 0.000162 | 0.021151 |
| Ttr | 7.562287 | 1.682079 | 0.000165 | 0.021334 |
| Sp4 | 1.477541 | 0.351297 | 0.000169 | 0.021745 |
| Zwilch | -3.01511 | 0.843766 | 0.000172 | 0.022044 |
| Slc12a7 | 0.889329 | 0.235024 | 0.000181 | 0.022801 |
| NA | 1.92374 | 0.48727 | 0.000181 | 0.022801 |
| Apln | -1.55109 | 0.504702 | 0.000183 | 0.02294 |
| Bcl2l2 | -0.84146 | 0.24306 | 0.000186 | 0.023158 |
| Mir147 | -2.99235 | 0.964532 | 0.000187 | 0.023189 |
| NA | -1.83897 | 0.458562 | 0.000189 | 0.023253 |
| Evx1os | 3.802135 | 0.93376 | 0.000202 | 0.024652 |
| Irag1 | 1.197024 | 0.318708 | 0.000206 | 0.024755 |
| C130026I2 | 1.135613 | 0.387782 | 0.000205 | 0.024755 |
| E130102H: | 2.907601 | 0.722683 | 0.000206 | 0.024755 |
| Ezh2 | -1.78192 | 0.485568 | 0.000211 | 0.025082 |
| Tef | 1.527605 | 0.399425 | 0.000216 | 0.025405 |
| Gm14760 | -2.38566 | 0.6157 | 0.000217 | 0.025405 |
| NA | -1.09422 | 0.282538 | 0.000215 | 0.025405 |
| Lmnb1 | -2.34837 | 0.653347 | 0.000219 | 0.025463 |
| Rbak | 0.829781 | 0.20314 | 0.000223 | 0.025786 |
| Fxr1 | -0.55463 | 0.153317 | 0.000227 | 0.025888 |
| Srsf9 | -0.86423 | 0.23971 | 0.000232 | 0.025888 |
| Gtf2e2 | -0.73796 | 0.198449 | 0.00023 | 0.025888 |
| Ccdc113 | 2.065322 | 0.530062 | 0.000233 | 0.025888 |
| Erfe | -3.17752 | 0.939357 | 0.000233 | 0.025888 |
| Rgma | 1.497703 | 0.39733 | 0.000226 | 0.025888 |
| Catspere2 | 1.8894 | 0.458474 | 0.000229 | 0.025888 |
| Fkbp4 | -0.7106 | 0.226937 | 0.000235 | 0.025916 |

| | | | | |
|---|---|---|---|---|
| Gm38416 | 2.571983 | 0.651709 | 0.000239 | 0.026172 |
| Zfp655 | 0.830278 | 0.207288 | 0.000249 | 0.026243 |
| Nptx1 | 3.449444 | 0.887433 | 0.000251 | 0.026243 |
| Ralgds | 1.018268 | 0.263418 | 0.000247 | 0.026243 |
| Itgam | -1.06251 | 0.323282 | 0.000251 | 0.026243 |
| Tpsb2 | 1.977191 | 0.73701 | 0.000241 | 0.026243 |
| Kctd12b | 0.808109 | 0.262026 | 0.000252 | 0.026243 |
| Cbx7 | 1.469841 | 0.358781 | 0.000251 | 0.026243 |
| Gm6161 | -2.53563 | 0.62785 | 0.000244 | 0.026243 |
| NA | 2.43879 | 0.60354 | 0.000251 | 0.026243 |
| Kcnmb4os | 2.896756 | 0.746579 | 0.000266 | 0.027252 |
| Cma1 | 3.808632 | 0.970487 | 0.00027 | 0.027252 |
| Ltb | 3.116431 | 0.776344 | 0.000269 | 0.027252 |
| Polq | -3.02638 | 0.985366 | 0.000269 | 0.027252 |
| Dlgap5 | -2.91415 | 0.898897 | 0.000264 | 0.027252 |
| NA | 1.790332 | 0.46783 | 0.000267 | 0.027252 |
| Rhbdd1 | 0.599577 | 0.163672 | 0.000274 | 0.027431 |
| Chic1 | 1.340923 | 0.330291 | 0.000276 | 0.027431 |
| Ighv10-1 | 9.017881 | 2.157949 | 0.000275 | 0.027431 |
| Rnd1 | -3.68559 | 1.019769 | 0.000279 | 0.027585 |
| Mgl2 | 2.360505 | 0.639883 | 0.000285 | 0.028005 |
| Bdnf | -4.07604 | 0.99264 | 0.000288 | 0.028146 |
| Exosc6 | -1.59598 | 0.468819 | 0.000289 | 0.028146 |
| Sfpq | -1.30433 | 0.361728 | 0.000292 | 0.028267 |
| NA | 3.751711 | 1.03361 | 0.000293 | 0.028267 |
| H4c9 | -2.24151 | 0.589681 | 0.000296 | 0.0284 |
| Rrm2 | -3.64438 | 1.050582 | 0.000298 | 0.028426 |
| NA | 3.838215 | 1.020459 | 0.000306 | 0.02906 |
| Rragd | 1.249643 | 0.34784 | 0.00031 | 0.029328 |
| Cenpn | -2.75498 | 0.858348 | 0.000313 | 0.029426 |
| Tcap | 3.697056 | 0.923296 | 0.000315 | 0.02949 |
| Cnksr3 | -1.21172 | 0.322192 | 0.000325 | 0.030109 |
| Srrm4 | -2.79302 | 0.704216 | 0.000326 | 0.030109 |
| NA | -1.32401 | 0.348089 | 0.000324 | 0.030109 |
| Cdc45 | -2.83063 | 0.841016 | 0.000334 | 0.030124 |
| Ska3 | -2.83126 | 0.912965 | 0.000334 | 0.030124 |
| Tnfrsf12a | -4.2386 | 1.057505 | 0.000336 | 0.030124 |
| Fbxw4 | -0.55374 | 0.200296 | 0.000335 | 0.030124 |
| Lig1 | -2.41793 | 0.727147 | 0.000328 | 0.030124 |
| Tbx3os1 | 1.335059 | 0.350578 | 0.000333 | 0.030124 |
| Ccl7 | -4.32488 | 1.166566 | 0.000342 | 0.03054 |
| Dtx1 | 3.080225 | 0.752892 | 0.000352 | 0.031042 |
| Snhg11 | 1.80957 | 0.556059 | 0.000353 | 0.031042 |
| Gm13394 | -1.70361 | 0.440621 | 0.00035 | 0.031042 |

| | | | | |
|---|---|---|---|---|
| Ccdc184 | 1.520167 | 0.62192 | 0.000356 | 0.031171 |
| Gm1604a | 0.731181 | 0.244562 | 0.000361 | 0.031495 |
| Anp32b | -1.3981 | 0.370357 | 0.000364 | 0.031565 |
| Eya1 | 1.802814 | 0.46682 | 0.000372 | 0.031978 |
| Hgsnat | -0.45092 | 0.122986 | 0.000372 | 0.031978 |
| NA | 3.269694 | 0.817213 | 0.000378 | 0.032404 |
| Cbx3 | -0.79197 | 0.254598 | 0.00038 | 0.032426 |
| Cxcl2 | -7.14872 | 1.843526 | 0.000388 | 0.032928 |
| Kif20b | -2.58518 | 0.84848 | 0.00039 | 0.032972 |
| Meg3 | 1.224001 | 0.540736 | 0.000394 | 0.033058 |
| NA | 2.70798 | 0.735303 | 0.000395 | 0.033058 |
| Aurka | -2.90803 | 0.895832 | 0.000401 | 0.033438 |
| Eepd1 | 1.060702 | 0.275765 | 0.000404 | 0.033447 |
| Haus3 | -1.38525 | 0.412688 | 0.000405 | 0.033447 |
| Tk2 | 0.926997 | 0.244354 | 0.00041 | 0.033752 |
| Leo1 | -1.23424 | 0.313704 | 0.000415 | 0.03393 |
| NA | -1.12661 | 0.344512 | 0.000416 | 0.03393 |
| NA | -2.03389 | 0.536533 | 0.000421 | 0.034226 |
| I830077J02 | 2.849975 | 0.861693 | 0.000425 | 0.034422 |
| Phf19 | -1.66062 | 0.587733 | 0.00043 | 0.034677 |
| Rflnb | -0.85177 | 0.218965 | 0.000435 | 0.034792 |
| NA | 2.131658 | 0.547148 | 0.000434 | 0.034792 |
| Igkj5 | 7.726055 | 1.992884 | 0.000448 | 0.035498 |
| NA | -2.07803 | 0.621533 | 0.000447 | 0.035498 |
| Polm | 0.813901 | 0.214789 | 0.000457 | 0.035891 |
| Acot11 | 0.746754 | 0.313084 | 0.000458 | 0.035891 |
| Rgs9bp | 2.471098 | 0.6976 | 0.000458 | 0.035891 |
| Gmnn | -2.56509 | 0.735412 | 0.000464 | 0.036145 |
| Kdm1b | 1.337275 | 0.357939 | 0.000482 | 0.037422 |
| NA | -1.02528 | 0.559042 | 0.000485 | 0.037547 |
| NA | -0.74072 | 0.27793 | 0.000498 | 0.03838 |
| Aph1a | -0.57669 | 0.148307 | 0.000507 | 0.038787 |
| Cyp3a57 | 1.600958 | 0.468927 | 0.000508 | 0.038787 |
| Stum | 2.359235 | 0.628122 | 0.000514 | 0.039132 |
| Klhl29 | 0.89779 | 0.323351 | 0.000517 | 0.039178 |
| Mphosph8 | -0.81904 | 0.2799 | 0.000525 | 0.039672 |
| Cenph | -2.865 | 0.968586 | 0.00053 | 0.03986 |
| Tardbp | -1.26629 | 0.368427 | 0.000532 | 0.039864 |
| Mtdh | -0.7355 | 0.208204 | 0.000538 | 0.039935 |
| Snx7 | -0.9786 | 0.284226 | 0.000541 | 0.039935 |
| Colgalt1 | -0.62785 | 0.162756 | 0.00054 | 0.039935 |
| Sae1 | -1.08425 | 0.321254 | 0.000536 | 0.039935 |
| Kif23 | -2.38169 | 0.813798 | 0.000547 | 0.040181 |
| Septin11 | -1.23373 | 0.31873 | 0.000551 | 0.040279 |

| | | | | |
|---|---|---|---|---|
| Ankdd1a | 2.183459 | 0.583403 | 0.000552 | 0.040279 |
| Kctd5 | -1.006 | 0.28474 | 0.000562 | 0.040388 |
| Snapc3 | -0.80058 | 0.229522 | 0.000557 | 0.040388 |
| Bmp8a | 1.797386 | 0.46886 | 0.00056 | 0.040388 |
| Cip2a | -2.57865 | 0.823735 | 0.00056 | 0.040388 |
| Dlgap2 | 4.274394 | 1.09956 | 0.000573 | 0.040991 |
| Hdgf | -1.13602 | 0.320909 | 0.000596 | 0.04191 |
| Gabrp | -2.94631 | 1.13308 | 0.000596 | 0.04191 |
| Nlrp10 | 1.341642 | 0.774816 | 0.000592 | 0.04191 |
| Mchr1 | 2.380516 | 0.792021 | 0.000595 | 0.04191 |
| Kcnip1 | 3.162574 | 0.846317 | 0.0006 | 0.04191 |
| Spdl1 | -2.70019 | 0.827724 | 0.000601 | 0.04191 |
| Zfp788 | 0.854361 | 0.22103 | 0.0006 | 0.04191 |
| Chordc1 | -0.71416 | 0.228936 | 0.000618 | 0.042069 |
| Bmp2 | 1.409656 | 0.374032 | 0.000618 | 0.042069 |
| Ptx3 | -5.67417 | 1.531732 | 0.000616 | 0.042069 |
| Gabrd | 3.400279 | 1.040335 | 0.000613 | 0.042069 |
| Fanci | -1.48735 | 0.469904 | 0.000608 | 0.042069 |
| NA | 3.565538 | 0.924467 | 0.000615 | 0.042069 |
| NA | 1.766815 | 0.542745 | 0.000618 | 0.042069 |
| Lin7b | 3.585665 | 0.909258 | 0.000621 | 0.042075 |
| Dab1 | 1.584407 | 0.428858 | 0.000623 | 0.042075 |
| NA | -0.71502 | 0.225222 | 0.000627 | 0.04218 |
| Rsad2 | 2.757717 | 0.743675 | 0.000629 | 0.04222 |
| Slc26a8 | 1.998088 | 0.530528 | 0.000641 | 0.04269 |
| NA | -2.75588 | 0.725534 | 0.000639 | 0.04269 |
| Gnl1 | -0.4266 | 0.152456 | 0.000652 | 0.043195 |
| Gsdmc3 | -2.00847 | 0.584439 | 0.000653 | 0.043195 |
| Ccne1 | -2.25621 | 0.822954 | 0.000664 | 0.043597 |
| Rtkn2 | -2.33265 | 0.658629 | 0.000663 | 0.043597 |
| Jchain | 3.513998 | 1.003238 | 0.000666 | 0.043624 |
| Xlr4a | 3.900693 | 1.211901 | 0.000674 | 0.043842 |
| Uchl1os | 1.818796 | 0.516103 | 0.000673 | 0.043842 |
| NA | -3.8171 | 1.073704 | 0.000677 | 0.043865 |
| Tjap1 | -0.85855 | 0.225356 | 0.000681 | 0.043972 |
| Nox1 | -0.91667 | 0.346082 | 0.000684 | 0.044002 |
| Slc22a21 | 1.299436 | 0.384153 | 0.000686 | 0.044002 |
| NA | 2.067429 | 0.536871 | 0.000691 | 0.044167 |
| Arf6 | -0.99267 | 0.280228 | 0.000695 | 0.044288 |
| Myocos | 4.52193 | 1.227499 | 0.0007 | 0.044468 |
| Gjb6 | 1.187449 | 0.33539 | 0.000704 | 0.044579 |
| Pi4k2b | -1.00481 | 0.272917 | 0.00072 | 0.045436 |
| Pxdc1 | -2.13035 | 0.589759 | 0.000723 | 0.04549 |
| Trip13 | -2.96046 | 0.947678 | 0.000728 | 0.045504 |

| | | | | |
|---|---|---|---|---|
| Ehd1 | -1.31962 | 0.349459 | 0.000727 | 0.045504 |
| Smad9 | 2.770338 | 0.741557 | 0.000731 | 0.045504 |
| Nos1ap | 1.778683 | 0.4637 | 0.000736 | 0.045661 |
| Cmc2 | -1.13693 | 0.409355 | 0.000745 | 0.045925 |
| Fam185a | -0.83801 | 0.325702 | 0.000742 | 0.045925 |
| Syt1 | 2.500809 | 0.661368 | 0.00075 | 0.046098 |
| Phf5a | -1.1211 | 0.30957 | 0.000752 | 0.046098 |
| Chaf1a | -2.50332 | 0.770023 | 0.000755 | 0.046101 |
| Larp7 | -1.1172 | 0.321642 | 0.000762 | 0.046101 |
| Zfp386 | 0.789654 | 0.233273 | 0.00076 | 0.046101 |
| Gm13135 | -2.03064 | 0.557589 | 0.000759 | 0.046101 |
| Aifm1 | -0.82413 | 0.261967 | 0.000788 | 0.047406 |
| Zfp975 | 0.915945 | 0.245412 | 0.000788 | 0.047406 |
| Ndor1 | 0.45536 | 0.126073 | 0.000792 | 0.047474 |
| Plekhb1 | 1.495295 | 0.393742 | 0.000803 | 0.047832 |
| NA | -1.1468 | 0.317429 | 0.000803 | 0.047832 |
| Lipc | 2.019394 | 0.570408 | 0.000813 | 0.048257 |
| Atad2 | -2.22642 | 0.686043 | 0.00083 | 0.048842 |
| Ncapg2 | -2.34842 | 0.770772 | 0.000827 | 0.048842 |
| NA | 1.404183 | 0.371957 | 0.00083 | 0.048842 |
| Ajuba | -2.32135 | 0.655945 | 0.000835 | 0.048943 |
| NA | -2.8588 | 0.981757 | 0.000838 | 0.048943 |
| Zfp712 | 1.311264 | 0.347641 | 0.000839 | 0.048943 |
| Herpud1 | 1.072782 | 0.283961 | 0.000852 | 0.049389 |
| Tacc3 | -2.61563 | 0.878207 | 0.000851 | 0.049389 |
| H2-Aa | 2.67463 | 0.698719 | 0.000857 | 0.049543 |
| Stat4 | 1.848412 | 0.694178 | 0.000861 | 0.049589 |
| Tedc1 | -2.66738 | 0.886087 | 0.000865 | 0.049655 |

Supplemental Table 5. d16 SCI + Pir vs SCI + CMC DEGs

| GeneID | Log2 fc | SE | p-value | FDR p-value |
|---|---|---|---|---|
| Col8a1 | -1.90387 | 0.187219 | 2.72E-24 | 5.53E-20 |
| Wisp2 | -3.43373 | 0.394695 | 3.33E-18 | 3.38E-14 |
| Pcsk5 | -1.21426 | 0.144047 | 3.47E-17 | 2.35E-13 |
| Ptprz1 | -2.97212 | 0.357361 | 9.03E-17 | 4.59E-13 |
| Gm43278 | -2.53053 | 0.333641 | 3.34E-14 | 1.15E-10 |
| Loxl2 | -1.93904 | 0.25572 | 3.39E-14 | 1.15E-10 |
| Eln | -2.23691 | 0.296209 | 4.29E-14 | 1.25E-10 |
| Cilp | -1.77564 | 0.26265 | 1.38E-11 | 3.50E-08 |
| Arnt2 | -2.61027 | 0.391365 | 2.56E-11 | 5.79E-08 |
| Actg2 | -1.6011 | 0.241285 | 3.23E-11 | 6.57E-08 |
| F13a1 | -1.38118 | 0.212976 | 8.86E-11 | 1.64E-07 |
| Loxl1 | -1.17571 | 0.184087 | 1.70E-10 | 2.87E-07 |
| Mir6950 | -2.11416 | 0.333079 | 2.19E-10 | 3.43E-07 |
| Crlf1 | -3.22494 | 0.510623 | 2.69E-10 | 3.91E-07 |
| Mustn1 | -1.27009 | 0.204915 | 5.71E-10 | 7.26E-07 |
| Fcna | -2.18994 | 0.354271 | 6.35E-10 | 7.41E-07 |
| Cmah | -1.83771 | 0.297685 | 6.69E-10 | 7.41E-07 |
| Gstm3 | 1.354015 | 0.219528 | 6.92E-10 | 7.41E-07 |
| Trdn | 2.355002 | 0.383033 | 7.83E-10 | 7.97E-07 |
| Fcnaos | -2.07467 | 0.338168 | 8.51E-10 | 8.25E-07 |
| Lgi2 | -1.47833 | 0.243586 | 1.29E-09 | 1.19E-06 |
| D030025P2 | -3.43586 | 0.571361 | 1.82E-09 | 1.49E-06 |
| Wisp1 | -1.9073 | 0.316942 | 1.77E-09 | 1.49E-06 |
| Col12a1 | -1.48817 | 0.247541 | 1.83E-09 | 1.49E-06 |
| Bmper | -2.35019 | 0.393394 | 2.31E-09 | 1.68E-06 |
| A830012C1 | -2.22674 | 0.372714 | 2.31E-09 | 1.68E-06 |
| Lox | -1.88058 | 0.314378 | 2.21E-09 | 1.68E-06 |
| Vgf | -4.32396 | 0.724909 | 2.45E-09 | 1.72E-06 |
| 1500009L1 | -2.14724 | 0.361364 | 2.82E-09 | 1.91E-06 |
| Fstl3 | -1.77216 | 0.301159 | 3.99E-09 | 2.62E-06 |
| Cd209d | -2.77858 | 0.478776 | 6.49E-09 | 4.13E-06 |
| Cd209f | -1.21322 | 0.212703 | 1.17E-08 | 7.01E-06 |
| Dbn1 | -1.20306 | 0.210788 | 1.15E-08 | 7.01E-06 |
| Msn | -1.38833 | 0.246469 | 1.77E-08 | 1.02E-05 |
| Lbp | -1.08489 | 0.192706 | 1.80E-08 | 1.02E-05 |
| 4930478K1 | -2.37847 | 0.423917 | 2.02E-08 | 1.08E-05 |
| Slc26a7 | -2.06973 | 0.368656 | 1.97E-08 | 1.08E-05 |
| Filip1l | -1.40758 | 0.251488 | 2.18E-08 | 1.11E-05 |
| 4833412C0 | -1.18505 | 0.211555 | 2.12E-08 | 1.11E-05 |
| Cthrc1 | -1.81095 | 0.323922 | 2.26E-08 | 1.12E-05 |
| Hhipl1 | -2.09841 | 0.379173 | 3.13E-08 | 1.45E-05 |
| Sfrp1 | -1.50278 | 0.271424 | 3.08E-08 | 1.45E-05 |

| | | | | |
|---|---|---|---|---|
| Gm15270 | -2.11878 | 0.38922 | 5.22E-08 | 2.23E-05 |
| Igf1 | -1.56401 | 0.287263 | 5.19E-08 | 2.23E-05 |
| P4ha3 | -1.73014 | 0.319291 | 6.00E-08 | 2.44E-05 |
| Mmp3 | -1.93871 | 0.363847 | 9.91E-08 | 3.95E-05 |
| Ltbp2 | -2.7288 | 0.51261 | 1.02E-07 | 3.98E-05 |
| Igfbp2 | -1.33183 | 0.254822 | 1.73E-07 | 6.63E-05 |
| Penk | -2.04394 | 0.391952 | 1.84E-07 | 6.88E-05 |
| Frzb | -1.42836 | 0.274016 | 1.86E-07 | 6.88E-05 |
| Aldh1a2 | -1.32045 | 0.253895 | 1.98E-07 | 7.21E-05 |
| Ankrd1 | -5.80298 | 1.124334 | 2.45E-07 | 8.75E-05 |
| Ngb | 1.975812 | 0.388653 | 3.70E-07 | 0.00013 |
| Cdsn | -1.53015 | 0.303324 | 4.54E-07 | 0.000157 |
| Arg1 | -3.02617 | 0.600751 | 4.72E-07 | 0.000158 |
| Lrg1 | -1.61913 | 0.321455 | 4.73E-07 | 0.000158 |
| Tnc | -1.65398 | 0.329703 | 5.26E-07 | 0.000173 |
| Gm37804 | -1.76805 | 0.353382 | 5.64E-07 | 0.000182 |
| Medag | -1.37725 | 0.275824 | 5.94E-07 | 0.000186 |
| Csgalnact1 | -1.10109 | 0.220503 | 5.93E-07 | 0.000186 |
| Mfap5 | -1.13541 | 0.227792 | 6.21E-07 | 0.000189 |
| Prss57 | -1.60053 | 0.321462 | 6.39E-07 | 0.000191 |
| Itgam | -1.01589 | 0.205653 | 7.82E-07 | 0.00023 |
| Fst | -1.69843 | 0.345247 | 8.68E-07 | 0.000252 |
| Il17b | -3.0863 | 0.627725 | 8.80E-07 | 0.000252 |
| Ccdc80 | -1.19024 | 0.242261 | 8.97E-07 | 0.000253 |
| Chl1 | -1.51072 | 0.309929 | 1.09E-06 | 0.0003 |
| Fbn1 | -1.24476 | 0.256225 | 1.19E-06 | 0.000321 |
| Gm20488 | -1.43928 | 0.299309 | 1.52E-06 | 0.000401 |
| Sfrp2 | -1.03494 | 0.215899 | 1.64E-06 | 0.000427 |
| Sync | -1.08773 | 0.22899 | 2.03E-06 | 0.000517 |
| Lgals1 | -1.43842 | 0.304367 | 2.29E-06 | 0.000575 |
| Tgfb3 | -1.52349 | 0.324212 | 2.61E-06 | 0.000648 |
| Adamts15 | -1.13563 | 0.243843 | 3.20E-06 | 0.000773 |
| Gm42901 | -1.04497 | 0.226436 | 3.93E-06 | 0.000899 |
| Plat | -1.18072 | 0.257279 | 4.45E-06 | 0.000973 |
| Fam150a | -4.41246 | 0.962529 | 4.56E-06 | 0.000986 |
| Pi16 | -1.02112 | 0.224378 | 5.34E-06 | 0.001144 |
| Des | -1.10258 | 0.242979 | 5.69E-06 | 0.001205 |
| Ppp1r18os | -1.22055 | 0.2695 | 5.93E-06 | 0.001243 |
| Fn1 | -1.09205 | 0.241567 | 6.16E-06 | 0.001272 |
| Itgam | -1.3225 | 0.295045 | 7.38E-06 | 0.001487 |
| Ccl6 | -1.35235 | 0.301908 | 7.49E-06 | 0.001493 |
| Cpa3 | -1.58679 | 0.354977 | 7.82E-06 | 0.001544 |
| Hspb1 | -1.38124 | 0.309693 | 8.19E-06 | 0.001603 |
| Cysltr1 | -1.15383 | 0.259977 | 9.07E-06 | 0.001741 |

| | | | | |
|---|---|---|---|---|
| Thbs4 | -2.37499 | 0.537503 | 9.94E-06 | 0.001889 |
| Adcy7 | -1.16684 | 0.265699 | 1.13E-05 | 0.0021 |
| Msln | -1.42852 | 0.32573 | 1.16E-05 | 0.00212 |
| Serpina3n | -2.25312 | 0.515573 | 1.24E-05 | 0.002216 |
| Diaph3 | -1.38636 | 0.31768 | 1.28E-05 | 0.002259 |
| Rgs4 | -2.48459 | 0.570754 | 1.34E-05 | 0.002333 |
| Col5a1 | -1.04846 | 0.241147 | 1.38E-05 | 0.00237 |
| Uhrf1 | -1.38519 | 0.319596 | 1.46E-05 | 0.00248 |
| Phlda2 | 1.185079 | 0.274816 | 1.62E-05 | 0.002694 |
| Rbm24 | 1.077396 | 0.250001 | 1.64E-05 | 0.002705 |
| 9330158H( | -1.7112 | 0.397309 | 1.66E-05 | 0.002715 |
| Synpo | -1.02094 | 0.237367 | 1.70E-05 | 0.00275 |
| Mamdc2 | 1.715609 | 0.398924 | 1.70E-05 | 0.00275 |
| Cwh43 | 1.247211 | 0.291381 | 1.87E-05 | 0.002989 |
| Serpina3i | -3.50877 | 0.821145 | 1.93E-05 | 0.003034 |
| Mcpt4 | -1.30966 | 0.306581 | 1.94E-05 | 0.003034 |
| Cit | -1.42888 | 0.33469 | 1.96E-05 | 0.003045 |
| Casc5 | -1.8851 | 0.441849 | 1.99E-05 | 0.00305 |
| Rxfp1 | -1.4112 | 0.33084 | 1.99E-05 | 0.00305 |
| Gm21451 | -1.79137 | 0.420885 | 2.08E-05 | 0.003103 |
| Vcan | -1.55815 | 0.366037 | 2.07E-05 | 0.003103 |
| Ms4a6d | -1.42986 | 0.335946 | 2.08E-05 | 0.003103 |
| Ccl9 | -1.35019 | 0.317311 | 2.09E-05 | 0.003103 |
| Ctgf | -2.06581 | 0.486595 | 2.18E-05 | 0.003193 |
| Tpsb2 | -1.36275 | 0.321635 | 2.27E-05 | 0.003292 |
| Gda | -1.25174 | 0.296193 | 2.38E-05 | 0.003406 |
| Figf | -1.0888 | 0.257963 | 2.43E-05 | 0.003463 |
| Fbln2 | -1.20381 | 0.286519 | 2.65E-05 | 0.003694 |
| Timp1 | -2.69576 | 0.64401 | 2.84E-05 | 0.00393 |
| Alox5 | -1.32594 | 0.317443 | 2.95E-05 | 0.004061 |
| Abcc2 | 2.50822 | 0.607666 | 3.67E-05 | 0.004937 |
| Folr2 | -1.06485 | 0.258489 | 3.80E-05 | 0.005081 |
| Tpx2 | -1.34998 | 0.328367 | 3.94E-05 | 0.005225 |
| Mki67 | -1.39308 | 0.339207 | 4.01E-05 | 0.005263 |
| Shisa3 | -1.81417 | 0.442768 | 4.18E-05 | 0.005414 |
| 2810430I1 | -1.46012 | 0.356737 | 4.26E-05 | 0.005429 |
| Klhdc8a | -1.06104 | 0.25916 | 4.24E-05 | 0.005429 |
| Timd4 | -2.07639 | 0.507647 | 4.31E-05 | 0.005438 |
| Clec11a | -1.00035 | 0.244642 | 4.33E-05 | 0.005438 |
| Gm44556 | -2.93878 | 0.720089 | 4.48E-05 | 0.005558 |
| 6030408B1 | -1.84632 | 0.456062 | 5.16E-05 | 0.006243 |
| Mthfd2 | -1.5222 | 0.377248 | 5.46E-05 | 0.006533 |
| Lrrc15 | -1.4609 | 0.362538 | 5.59E-05 | 0.006638 |
| Serpinb9b | -2.72965 | 0.678949 | 5.81E-05 | 0.006798 |

| | | | | |
|---|---|---|---|---|
| Rhoc | -1.00621 | 0.250615 | 5.95E-05 | 0.006911 |
| Gm21284 | -1.62144 | 0.40454 | 6.12E-05 | 0.006942 |
| Vash1 | -1.26835 | 0.316164 | 6.03E-05 | 0.006942 |
| Scx | -1.65388 | 0.414257 | 6.54E-05 | 0.00735 |
| Aspn | -1.59764 | 0.401137 | 6.81E-05 | 0.007489 |
| Fcgr2b | -1.0653 | 0.267408 | 6.78E-05 | 0.007489 |
| Sla | -1.02832 | 0.258177 | 6.80E-05 | 0.007489 |
| Cenpi | -2.23707 | 0.562077 | 6.89E-05 | 0.007536 |
| Adamtsl2 | -1.0958 | 0.27546 | 6.95E-05 | 0.007557 |
| Spock3 | 1.655922 | 0.418641 | 7.64E-05 | 0.00818 |
| Prss12 | -1.17232 | 0.297387 | 8.08E-05 | 0.008603 |
| Hspb7 | -1.44484 | 0.366652 | 8.13E-05 | 0.008609 |
| Hsp25-ps1 | -1.33846 | 0.339817 | 8.19E-05 | 0.008631 |
| Chrm3 | -1.1566 | 0.29411 | 8.40E-05 | 0.008812 |
| Tmem200a | -1.60743 | 0.4091 | 8.52E-05 | 0.00889 |
| Ltc4s | -1.12588 | 0.287633 | 9.07E-05 | 0.009409 |
| Iqgap3 | -1.69656 | 0.433799 | 9.19E-05 | 0.009492 |
